# Redox-dependent lipophilicity of phenazine metabolites is modulated by intramolecular hydrogen bonds and controls their biological distribution

**DOI:** 10.64898/2026.04.18.719255

**Authors:** Korbinian O. Thalhammer, Matthew Scurria, Jinyang Li, Inês B. Trindade, Osvaldo Gutierrez, Stuart J. Conway, Dianne K. Newman

## Abstract

Phenazines are redox-active microbial metabolites produced and secreted in diverse ecological contexts from soils to chronic infections. In these disparate environments phenazines can function variously as antibiotics, extracellular electron shuttles, and nutrient scavengers. Key to understanding the impact of these functions is a robust expectation of phenazine retention or diffusion in a given context. But predicting phenazine fate and transport is difficult because of the chemical complexity of their local microenvironments. To address this challenge, we measured the octanol water distribution coefficient (LogD) as a proxy for lipophilicity of three naturally occurring phenazines produced by the opportunistic pathogen *Pseudomonas aeruginosa*: phenazine-1-carboxylic acid, phenazine-1-carboxamide, and pyocyanin. We investigated the behavior of both oxidized and reduced forms of these phenazines across broad ionic strength and pH conditions. While the ionic context exerts only small effects, the pH and redox state contribute strongly and independently to changes in phenazine lipophilicity. The pH trends are expected, but the observed redox dependence is generally missed by existing lipophilicity calculation methods. Additional LogD measurements with 1-hydroxyphenazine and unsubstituted phenazine, together with density functional theory modeling of phenazines in their reduced and oxidized forms, reveal that intramolecular hydrogen bonding contributes significantly to the increased lipophilicity of reduced phenazines that possess H-bond accepting substituents in the 1-position. These results explain phenazine behavior in a biological context: redox state alone significantly alters retention of pyocyanin in planktonic *P. aeruginosa* cells, with the reduced species being predominantly retained by membranes. We propose that the modulation of phenazine lipophilicity in response to the local redox environment has evolved to give a competitive advantage to bacteria by retaining or dispersing these bioactive molecules. Beyond improving our understanding of natural phenazine fate in diverse microbial contexts, our results emphasize an oft-overlooked theme relevant to rational drug and electrochemical shuttle design: redox state matters.

## Introduction

*Pseudomonas aeruginosa* is a common opportunistic human pathogen that colonizes a wide range of habitats and is known to establish both acute and chronic infections in hospitalized and immunodeficient patients. This bacterium exhibits a high level of metabolic flexibility that enables it to exist in the broad range of environments in which it is found^1^. For *P. aeruginosa* and other microbes, secretion of primary and secondary metabolites can confer a competitive advantage through multiple mechanisms including redox balancing^2^ and competitor inhibition^3^. Metabolite secretion is controlled by a variety of systems, including ATP-driven transport systems, antiporters, symporters, uniporters, and passive diffusion^4^. While the selectivity of enzymatic transport systems depends on structural recognition of the metabolite, passive diffusion across the membrane is governed primarily by the metabolite’s physicochemical properties, especially its lipophilicity.

Here we examine the lipophilicity of phenazine metabolites, an important class of secreted, redox-active secondary metabolites produced by *P. aeruginosa* as well as many other bacterial species found in natural soils and waters around the world^5^. Phenazines are known to play critical roles in nutrient scavenging, virulence, competitor inhibition, and signaling^6^. These functions rely on reversible redox transformations that can be triggered by producer cells, host tissue, competing microbes, and abiotic phases like oxygen and iron oxide minerals^7,8^. Despite this, almost nothing is known about the mechanisms by which phenazine concentrations are spatially controlled, and whether the local redox environment affects compound distribution^9^.

In chemically complex environments, lipophilicity is a useful simplifying parameter for understanding and predicting compound distribution. In practice, scientists in disciplines including environmental science and medicinal chemistry lean heavily on the octanol-water partition coefficient (LogP) or the pH-specific distribution coefficient (LogD_pH_) as useful proxies for compound lipophilicity (Equation 1). LogP and LogD correlate strongly with affinity for biological phases, including cell membranes and soil organic matter, which are typically more lipophilic than water^10,11^. As a result, LogP is a useful predictor for the bioaccumulation of anthropogenic toxins such as DDT^12^. By the same token, LogP and LogD feature prominently in drug design because of the correlation between the lipophilicity of a drug, its target promiscuity, and our *in vivo* absorption and retention of the molecule^13,14^. While these values can be measured using the shake flask method or chromatography^14^, they are frequently predicted using a range of different algorithms that report calculated LogP or LogD (cLogP or cLogD). Both measurement and prediction of LogP/LogD tend to neglect changes in molecular redox state that are common in biological settings and are relevant to understanding the behavior of redox active metabolites.

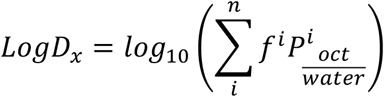

**Equation 1**. The distribution coefficient, LogD_x_, accounts for the partitioning of all ionization forms of a solute between the octanol and the water phase at pH x. The total number of ionization states is given by *n. f*^*i*^ is the mole fraction of each form of the molecule (controlled by the pKa of the solute and the solution pH) and *P*^*i*^ is the partition coefficient for that form. For solutes that do not ionize, LogD_x_ collapses to the familiar partition coefficient LogP, which is simply the ratio of the concentrations of solute in octanol and in water.

To explore the redox sensitivity of phenazine lipophilicity, as a case study through which to elucidate general principles, we examined the three main phenazines produced by *P. aeruginosa*. Like most phenazines, these compounds -- phenazine-1-carboxylic acid (PCA), phenazine-1-carboxamide (PCN), and pyocyanin (PYO) -- undergo reversible, coupled 2e^−^, 2H^+^ redox transformations under physiological conditions (Scheme 1). We systematically subjected oxidized and reduced samples of these three phenazines to environmentally relevant pH and ionic strength conditions and measured their octanol/water distribution coefficients. Our results depended heavily on redox state and deviated substantially from predictions made by four leading cLogP prediction algorithms.

To expand our understanding beyond the subset of *P. aeruginosa* phenazines, we conducted additional experiments with phenazine and 1-hydroxyphenazine and performed density functional theory analysis on the compounds studied. We show that lipophilicity, and therefore the expected biological distribution of some phenazines, is modulated in a redox-dependent manner. We suggest this might be a hitherto unappreciated and more broadly applicable mechanism by which the spatial distribution of bioactive molecules is controlled by certain bacteria. These findings have important ramifications for the prediction of redox-active compound lipophilicity, demonstrating that it is essential that the cLogP or cLogD of the compound is calculated not only at the appropriate pH, but also in the relevant redox state. Even then, care must be taken to ensure the chosen algorithm captures the effects of redox-mediated intramolecular hydrogen bonds. Taken together, our results suggest that redox state exerts a fundamental control on lipophilicity for natural and artificial phenazines, and likely for other similar redox-active molecules, a fact that is both underappreciated experimentally and not accounted for *in silico*.

**Scheme 1.**
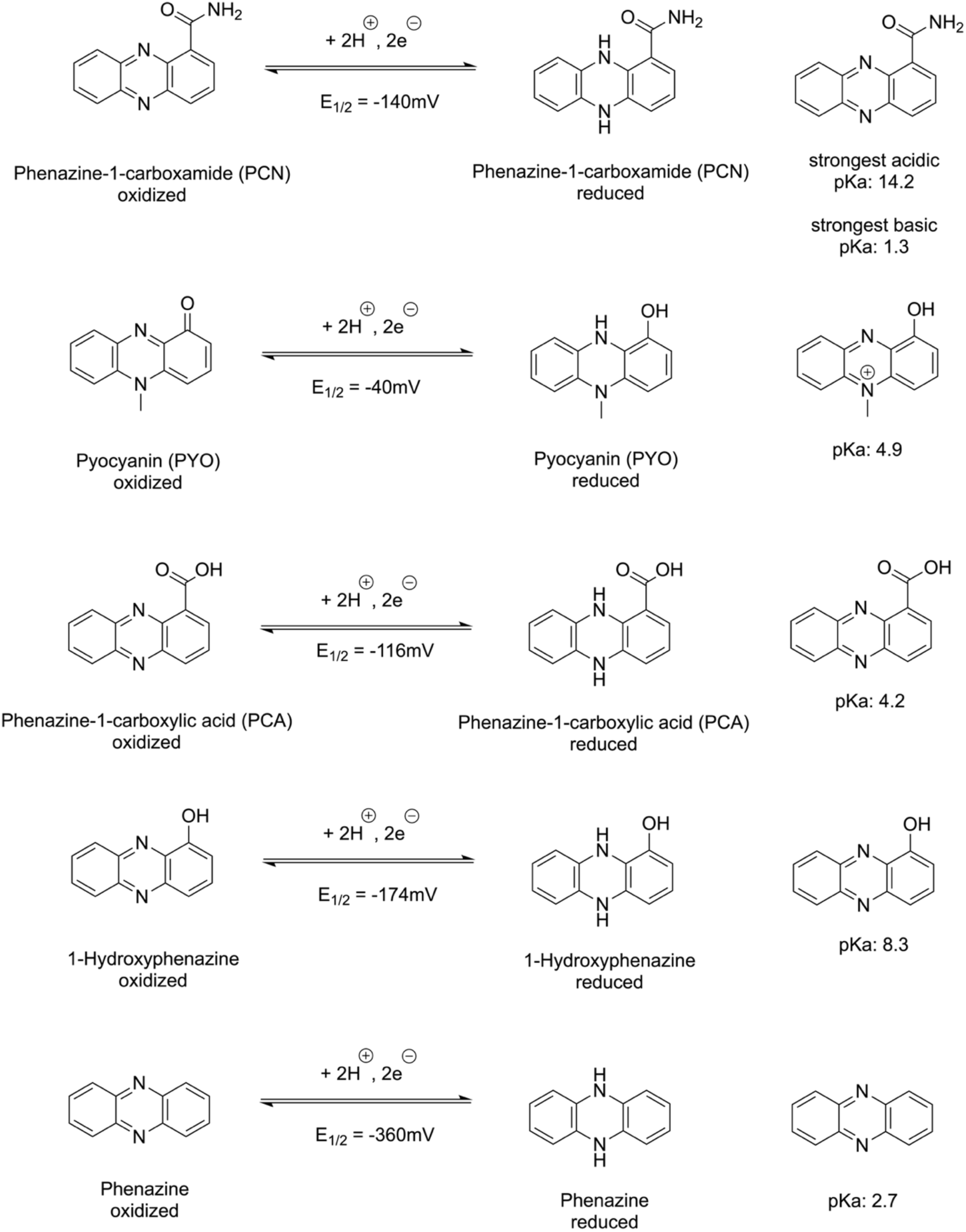
The redox equilibria and pKa values of phenazine-1 carboxamide (PCN), pyocyanin (PYO), phenazine-1 carboxylic acid (PCA), 1-hydroxyphenazine, and phenazine^15-19^. All potentials are relative to the Standard Hydrogen Electrode^8^. PCN is not predicted to change ionization state within the physiological pH range (3-10) explored in this study^15^. Experiments with 1-hydroxyphenazine and phenazine were only conducted at pH 7.2

## Results and Discussion

### Existing algorithms fail to predict redox dependence of phenazine lipophilicity

Experimental determination of LogP values is more time and resource intensive than calculation of cLogP values, so we began our study by comparing our initial experimental measurements with cLogP values from four leading prediction algorithms: miLogP, iLOGP, KOWWIN, and XLOGP3^20-28^ (see Supplement for additional algorithm details). Because the algorithms are primarily trained on neutral compounds, we limited our comparison to neutral forms of oxidized and reduced PCA, PCN, PYO, and 1-OH phenazine. But despite the relative simplicity of these phenazine structures, no individual algorithm accurately predicted the redox sensitivity of their lipophilicities (Figure 1). The fragment-based miLogP algorithm, developed using a test set of more than 12,000 mostly drug-like molecules, predicts essentially no offset between oxidized and reduced phenazines. The iLOGP algorithm, which considers a whole molecule’s solvation energy in water and in octanol, does predict offsets between oxidized and reduced phenazines. But it generally predicts the oxidized form of a phenazine to be more lipophilic, which is the opposite of the experimental trend. The most accurate algorithm, KOWWIN from the US Environmental Protection Agency, is based on atomic and molecular fragment contributions adjusted with empirical correction factors to account for more complex behavior like hydrogen bonding. The KOWWIN algorithm accurately predicts the enhanced lipophilicity of the reduced forms of PCN, PCA, and PYO, but it exaggerates the effect for PCN and predicts the wrong directionality for 1-OH phenazine, predicting that 1-OH phenazine is more lipophilic when reduced (the opposite is observed experimentally). The atom-based XLogP3 algorithm makes a similar error with 1-OH phenazine and predicts an unobserved redox-sensitive offset for phenazine, suggesting that while these two algorithms accurately capture some aspects of phenazine behavior, they are still missing some subtleties of phenazine redox chemistry. The algorithms all dramatically overestimate the lipophilicity of PYO in both redox forms. Given that these algorithms fail to predict experimental lipophilicities for neutral phenazines, and the fact that several *P. aeruginosa* phenazines are charged under physiological conditions, we continued our investigation by systematically measuring reduced and oxidized phenazine LogD values for PCN, PCA, and PYO across biologically relevant pH and ionic strength conditions.

**Figure 1:**
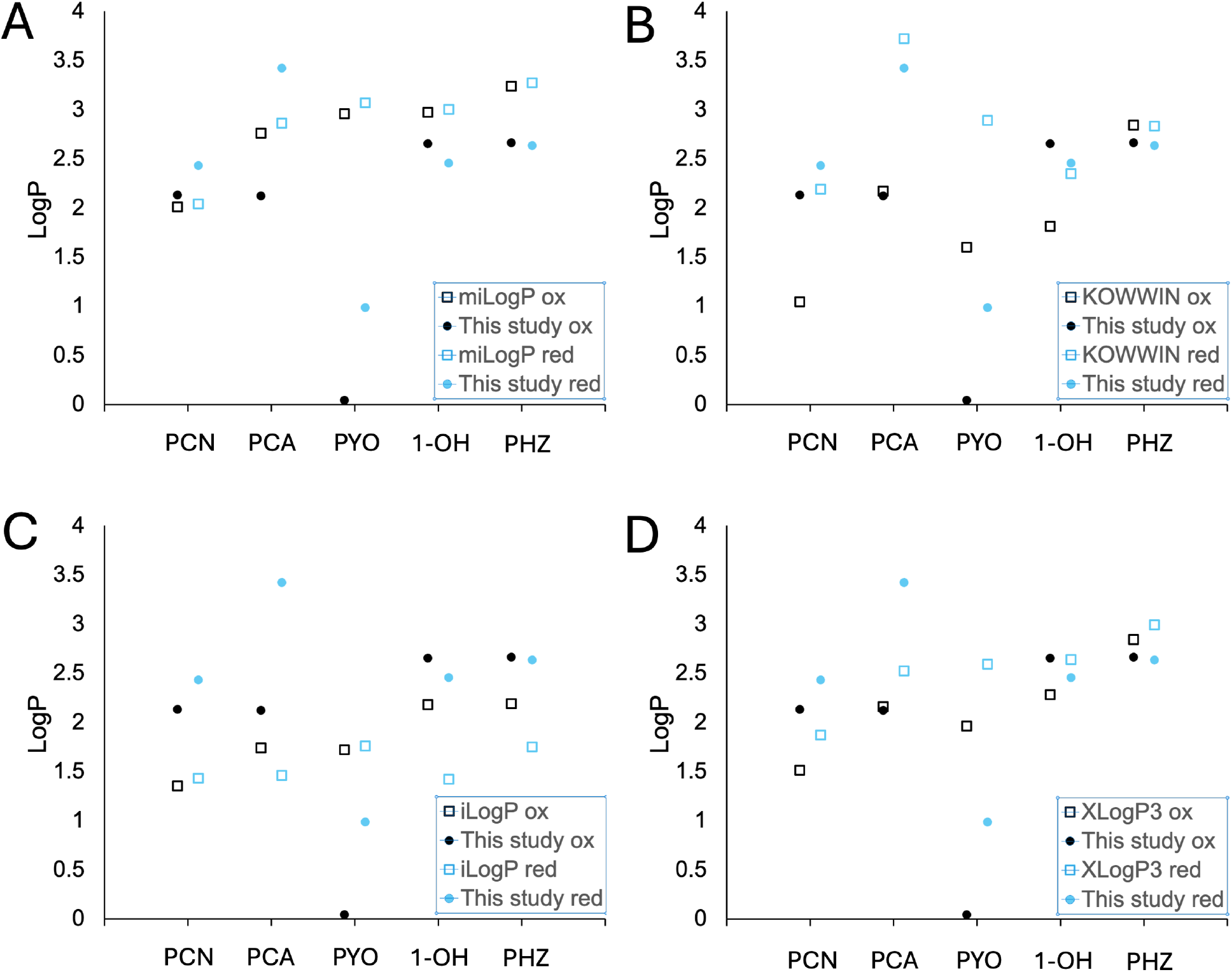
Existing cLogP prediction algorithms fail to capture redox-dependent lipophilicity changes. In all panels, closed circles denote experimental values from this study, while open squares indicate cLogP predictions. In all cases, black symbols are for the oxidized phenazine while blue symbols are for the reduced phenazine. A) miLogP predictions. B) KOWWIN predictions C) iLogP predictions. D) XLogP3 predictions. All measurements and predictions are for neutral forms of the molecules, meaning experiments were conducted at pH 7.2 except in the case of PCA (pH 3).

### Phenazine lipophilicity is sensitive to both functionalization and redox state

Across the pH range 3 to 10, reduced PCN maintains a slightly elevated lipophilicity relative to the oxidized form (Figure 2A). The difference is most pronounced at high pH. But in a given redox form, PCN exhibits only small variations in LogD between low and high pH, consistent with its predicted pKa values of 14.2 (strongest acidic) and 1.3 (strongest basic), which suggest it exists solely in its neutral form across the measured pH range^15^. In contrast, PCA contains a carboxylic acid with a pKa of 4.24 ^17^. Above this pKa, the majority of the molecules present will be ionized, while below the pKa, we expect the protonated, neutral carboxylic acid form to dominate, permitting greater solubility in the octanol phase and increasing measured lipophilicity. As expected, at low pH, PCA is significantly more lipophilic than at neutral or high pH, regardless of redox state. But in all cases, reduced PCA is more lipophilic than oxidized PCA by about an order of magnitude (Figure 2B). Remarkably, according to our measurements, the reduced, neutral form is over 5 orders of magnitude more lipophilic than the oxidized, ionized form. This large range in measured values is supported by the qualitative observation that the neutral, reduced form is highly insoluble in water, while the ionized, oxidized form is highly soluble in water^29^. At all pH values, reduced PYO is more lipophilic than oxidized PYO (Figure 2C). The difference in lipophilicities is most significant at low pH, where both forms of the molecule are expected to be charged. But a separation of an order of magnitude persists even at high pH, where both forms are neutral^16^. The influence of ionic strength was also investigated systematically, but as results conformed to expectations, the data is shown in the supplement (Figure S18).

**Figure 2.**
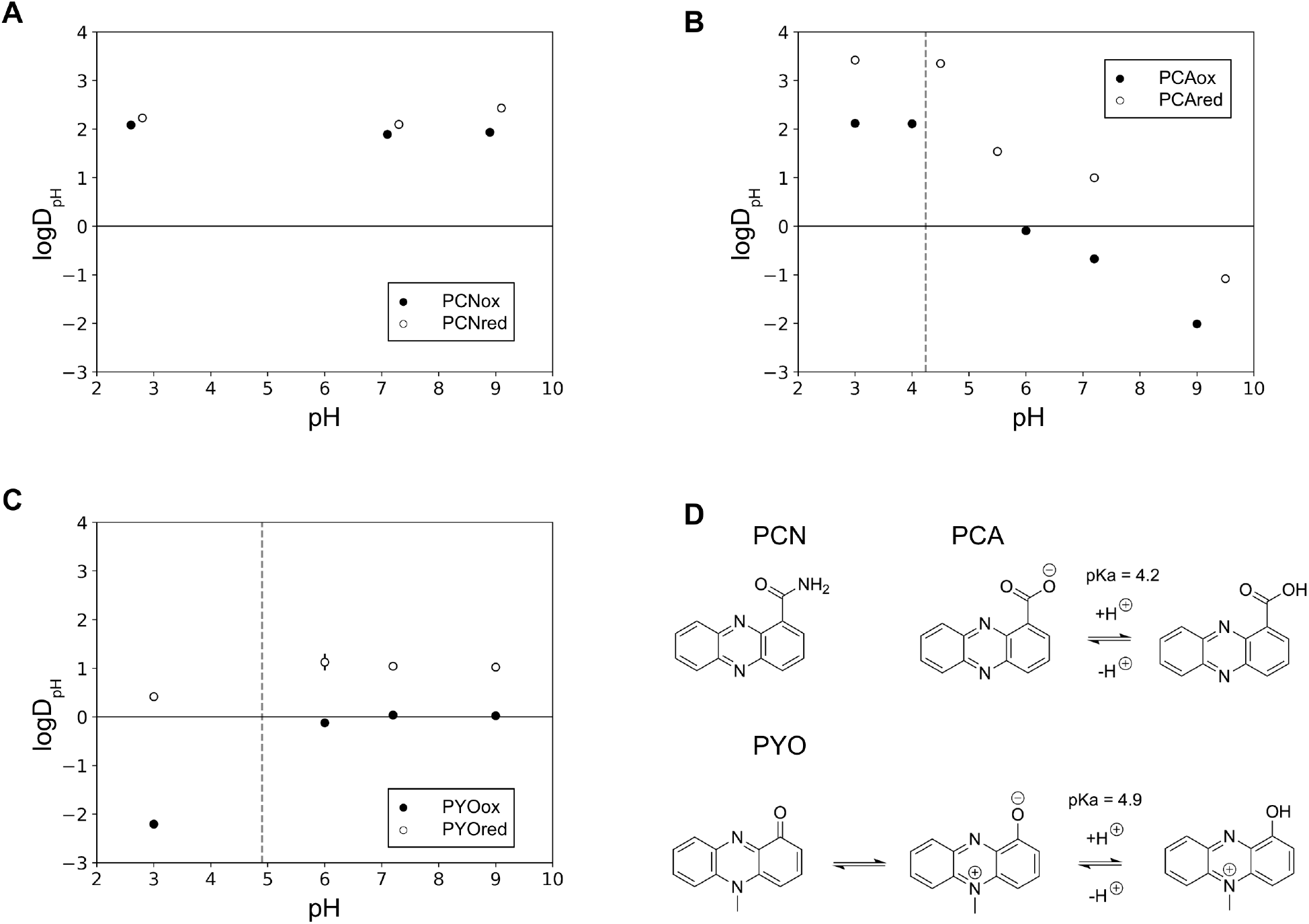
Phenazine lipophilicity is differentially sensitive to pH. Panels A, B, and C: measured LogD_pH_ values for PCN, PCA, and PYO, respectively. Within each panel, open circles represent the reduced form of the phenazine, while closed circles represent the oxidized form. Dashed lines denote pKa values of the oxidized phenazines from the literature. When greater than the size of the marker, standard deviations are depicted as vertical lines. Phenazines were equilibrated in 20 mM MOPS that was overwhelmed with small amounts of 100 mM HCl or NaOH to achieve the desired pH. Concentrations of phenazines in both the octanol and water phases were measured by HPLC. Panel D: protonation states with previously measured pKa values^16, 17^. PCN is not predicted to change ionization state within the physiological pH range (3-10) explored in this study^15^.

### Intramolecular hydrogen bonding contributes significantly to lipophilicity changes

Given the limited structural change between oxidized and reduced phenazines, we were interested in understanding the sometimes substantial redox state-dependent changes in compound lipophilicity. That redox state does not substantially affect the lipophilicity of unsubstituted phenazine (LogD = 2.66 oxidized, 2.63 reduced) led us to hypothesize that the 1-position substituents are likely responsible for the observed changes. The key difference between the oxidized and reduced form of phenazines is the addition of a hydrogen atom to the pyrazine nitrogen, converting it from a hydrogen bond acceptor to a hydrogen bond donor. This changes the ability of this moiety to interact with the substituent at the 1-position on the phenazine ring. To test this hypothesis, we probed phenazine-solvent interactions using density functional theory (see Supporting Information for more details). The strength of the *intra*molecular hydrogen bond was calculated for all structures, comparing the hydrogen bonded ‘closed’ form, *versus* the non-hydrogen bonded ‘open’ form conformation (Figure S1). These were established based on the presence of a new intramolecular hydrogen bond between the 1-position substituent and pyrazine nitrogen upon rotation of the substituent at the 1-position^30^. Energies of solvation in octanol and in water were then calculated and used to predict the effects of intramolecular hydrogen bonding on the cLogD value of a given molecular form (Figure S2 and S5)^31-33^. To compare lipophilicities across redox states, we performed natural bonding orbital (NBO) analysis of phenazines in their reduced and oxidized forms, analyzing for constructive orbital overlap, indicating favorable hydrogen bonding between the pyrazine nitrogen and 1-position substituent (Table S1)^34, 35^. We reasoned that a larger energy contribution generated from the second order perturbation theory analysis would result in a stronger NBO interaction which would deter *inter*molecular hydrogen bonding with water, thus increasing overall lipophilicity.

We began by looking at PCA, which, at physiological pH, is negatively charged and can form a strong (14.9 kcal/mol, Figure 3A) six-membered hydrogen bonded ring with the carboxylate group in its reduced form, an interaction that is not possible in the oxidized form. We speculated that the formation of this hydrogen bond disperses some of the carboxylate negative charge, resulting in less effective solvation of the molecule by water and consequently an increased lipophilicity compared to the oxidized form. The same situation is observed in PYO: no hydrogen bond is present in the oxidized form, but in the reduced form a hydrogen bond (0.71 kcal/mol, Figure 3B) can form between the pyrazine NH and the 1-position hydroxyl group. Again, this results in the reduced pyocyanin being more lipophilic than the oxidized form, as the oxidized form can be more readily solvated by water in the absence of the intramolecular hydrogen bond.

**Figure 3:**
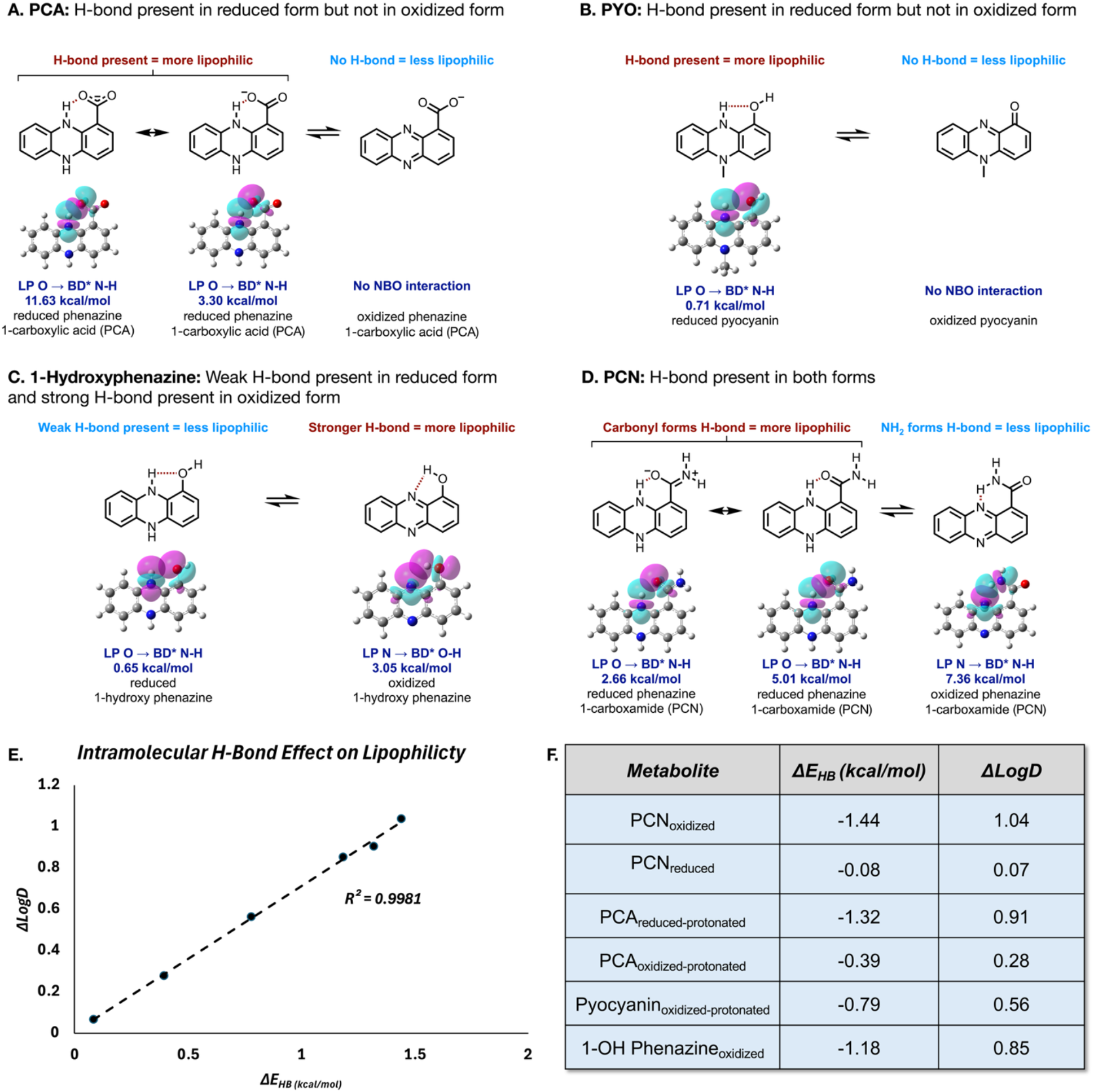
Intramolecular hydrogen bonding contributes significantly to lipophilicity changes. Natural Bonding Orbitals with their respective interaction energies generated by the intramolecular hydrogen bond calculated at the uwb97xD/cc-pvDZ-cpcm(H_2_O) level of theory for PCA (A), PYO (B), 1-hydroxyphenazine (C), and PCN (D). Predicted hydrogen bonds are indicated using dotted maroon lines. (E&F) Change in LogD values (ΔLogD) due to the formation of an intramolecular hydrogen bond plotted *vs* the change in intramolecular hydrogen bond strength (ΔE_HB_) due to changes in solvent, calculated at the uwb97xD/6-311+g(d,p)-SMD(solvent)//uwb97xD/cc-pvDZ-cpcm(solvent) level of theory. A larger difference in E_HB_, when going from water to octanol, directly correlates to a larger increase in lipophilicity when going from the open to closed form. Detailed structures and calculations can be found in the Supporting Information (Figure S6-S13).

Interestingly, 1-hydroxyphenazine exhibits the opposite change in lipophilicity between the redox states; here the oxidized form is more lipophilic than the reduced form. The hydrogen bonding situation here is more complex, as a 5-membered hydrogen bond can form in both redox states. In the oxidized form of 1-hydroxyphenazine the pyrazine nitrogen lone pair forms a stronger hydrogen bond (3.1 kcal/mol, Figure 3C) with the 1-position hydroxyl group. In the reduced form, the 1-position phenolic oxygen donates its lone pair to form a hydrogen bond with the hydrogen atom attached to the pyrazine nitrogen. This hydrogen bond is weaker (0.65 kcal/mol Figure 3C), presumably because the oxygen atom is a less effective lone pair donor compared to the nitrogen atom, due to its increased electronegativity, resulting in less electron donation into the antibonding orbital of the N-H bond^36^. The weaker hydrogen bond in the reduced form enables more effective solvation by water, meaning that it is less lipophilic than the oxidized form.

When examining PCN, we noticed a more subtle difference in lipophilicity between redox states, which can be explained by analyzing the difference in strength of the intramolecular hydrogen bonds. Due to the structure of PCN, a hydrogen bond can again form in both the reduced and oxidized forms. In the reduced form, a hydrogen bond is predicted to form between the pyrazine NH and the amide carbonyl oxygen atom. In the oxidized form the nitrogen atom of the pyrazine ring donates its lone pair to form a hydrogen bond with one of the amide NH_2_ hydrogen atoms. The strengths of these hydrogen bonds are predicted to be similar (Figure 3D), likely explaining the more similar lipophilicity between the two redox forms. The lower lipophilicity of the oxidized form is proposed to result from easier solvation of the more electronegative carbonyl oxygen atom, which is more readily presented to the solvent in this form.

To isolate and analyze the effect of the intramolecular hydrogen-bond on lipophilicity, we compared the change in LogD values (ΔLogD) due to the formation of an intramolecular hydrogen bond to the change in intramolecular hydrogen bond strength (ΔE_HB_) due to solvent changes (Figure 3F; see Supporting Information). As expected, a larger difference in the strength of the hydrogen bond in water as compared to octanol directly correlates to a larger increase in lipophilicity when going from the open to closed form. This demonstrates that the strength of the intramolecular hydrogen bond in octanol will directly impact lipophilicity. Our calculations also suggest that for a given redox state, closed forms of PCA, PCN, and PYO all exhibit increased lipophilicity compared to their ‘open’ forms.

Taken together, the data described above suggest that intramolecular hydrogen bonding contributes to enhanced lipophilicity of reduced forms of PCA>PYO>PCN, which aligns well with our experimental observations. Intramolecular hydrogen bonding in 1-OH contributes to slightly increased lipophilicity in the oxidized form. As there is no possibility for intramolecular H-bonding in either the oxidized or reduced forms of unsubstituted phenazine, these compounds exhibit essentially the same lipophilicity.

### Redox state controls biological PYO retention

As a case study to assess the biological consequences of redox-mediated changes to PYO’s lipophilicity, we experimented with late stationary phase *P. aeruginosa* PA14 cells in liquid culture. In the first experiment, cells were grown for 20 hours, then subdivided into two aliquots – one centrifuged and washed under anoxic conditions with sparged buffer, where we expect the phenazines to be and remain reduced, the other under normoxic conditions (atmospheric O_2_, 21%) with unsparged buffer, where we expect the phenazines to remain oxidized. Extracellular PYO concentrations were measured in the culture supernatant and wash supernatant by HPLC (see Supporting Information for details). After three washes with N_2_-sparged anoxic buffer, the cells continued to release a quantifiable amount of PYO (3.5 ± 0.4 µM). And after a fourth wash with normoxic buffer, the cells released 11.3 ± 0.1 µM PYO. On the other hand, the cells washed with normoxic buffer released PYO concentrations that were below the limit of quantification (sub 150 nM) by the third wash and that could not be detected (sub 25 nM) by the fourth wash. Together, these results demonstrate that the more lipophilic reduced form of PYO is strongly retained by the bacteria, while the less lipophilic oxidized form of PYO is easily washed away by aqueous buffer. The redox-mediated change to PYO’s lipophilicity explains this observed change in cellular retention, which has previously been observed to interfere with measurement of intracellular NADH/NAD^+^ ratios: measurements of NADH/NAD^+^ ratios in pyocyanin producers must be made under anoxic conditions because, in normoxic conditions, oxidized PYO skews measurements by diffusing out of the cell pellet and directly oxidizing NADH, leading to artificially low NADH/NAD^+^ ratios^37^. Moreover, hitherto puzzling observations of quenching of certain oxygen probes by bacterial cultures containing reduced PYO but not oxidized PYO can now be understood in light of our results^38^.

Unexpectedly, we observed a large difference between the supernatant PYO concentrations in the normoxic aliquots of cells (60 ± 7 µM) compared to the anoxic aliquots of the same culture (34 ± 3 µM). Calculations made by approximating the volume of the cells suggest that intracellular PYO concentrations in the anoxic condition reached millimolar levels (estimated at >3 mM), roughly 2 orders of magnitude greater than extracellular concentrations observed in the supernatant (see Supporting Information for details).

To validate this estimate and determine whether partitioning into the membrane is responsible for biological PYO retention, we conducted an additional experiment in which 1L of dense *P. aeruginosa* cells grown aerobically in LB (OD_500_ = 4.7), were removed from the shaking incubator and allowed to reduce their extracellular phenazines while standing on the benchtop (the culture changed from blue/green, indicating oxidized PYO, to colorless, indicating reduced PYO). Cells were then pelleted and lysed using a high pressure homogenizer, and the membrane fraction was isolated by ultracentrifugation. Throughout this process, the pellet, the lysed cells, and the membrane fraction remained colorless, presumably due to abiotic reactions between PYO and NADH and other residual cellular reducing agents. The membrane fraction was weighed and then extracted with MeOH. Upon addition of MeOH during the first extraction, the membrane fraction and added solvent turned bright blue, indicating oxidized PYO. After the third extraction, the ultracentrifuged membrane pellet and the overlying MeOH were colorless. Both the pooled membrane extract and the original culture supernatant were measured by HPLC to determine PYO concentrations. While the extracellular medium contained 0.03 µmol PYO per gram medium (essentially the same concentration as measured in the anoxic aliquot from the first biological retention experiment), the wet membrane fraction contained nearly two orders of magnitude more: 2.2 µmol PYO per gram. This suggests that membrane retention, which is expected to correlate directly with lipophilicity, is responsible for significant retention of reduced PYO *in vivo*.

### Conclusion

Our results indicate that even minor changes to the substituent at the 1-position (or the symmetric 6-position) can significantly alter phenazine lipophilicity by facilitating intramolecular hydrogen bonding. If a 1-position substituent can accept a hydrogen bond from a reduced pyrazine nitrogen, the lipophilicity of the reduced form compared to the oxidized form can be enhanced by over an order of magnitude. The sensitivity of a fundamental property like lipophilicity to substitutions at the 1-position is particularly intriguing given PCA or its symmetric counterpart 1,6-phenazine dicarboxylic acid are intermediates in all the known biosynthetic pathways for microbial phenazine. Changes in lipophilicity due to the local pH and redox environment, especially dramatic changes of the magnitude observed for PCA, likely dictate natural phenazines’ abilities to traverse membranes, biofilms, soils, and tissues. In general, higher lipophilicities are expected to correlate with increased retention in organic phases and decreased water solubility and aqueous transport. As is the case for quinones and quinols, changes in lipophilicity likely also influence phenazine affinities for protein binding pockets^39^, enhancing or limiting their interaction with components of the electron transport chain or with detoxifying or degrading enzymes.

The ability to effect such significant changes to a fundamental thermodynamic property through the modification of a single molecular substituent gives phenazine producers a remarkable evolutionary handle for tuning the functions of these secreted metabolites to the needs most pressing in their respective niches. Unlike many secondary metabolites, including most antibiotics, which are large or complex enough that their basic properties are difficult to change without significant structural modification, phenazine lipophilicity is highly sensitive to minor structural alterations, which can often be accomplished by a single enzyme. Perhaps this flexibility helps explain their prevalence and structural diversity across diverse niches^6, 40^, including host-microbe systems.

In the case of reduced pyocyanin, the measured enhancement in lipophilicity predicts increased biological retention, which is observed in anoxic cell pellets, despite the expected expression of an entire suite of efflux pumps capable of actively pumping phenazines out of the cell^41, 42^. There is thus good reason to believe that lipophilicity changes can play an important role even in contexts where active biological processes compete with abiotic processes like diffusion and partitioning. The extremely elevated levels of pyocyanin measured in the membrane suggest a different biological role than is typically appreciated for this metabolite, which is often treated as both toxic and primarily extracellular. One possible explanation is that in oxygen limited environments, PYO, and possibly other phenazines, play a direct role like that of quinones in the *P. aeruginosa* electron transport chain. In the archaeal methanogen *Methanosarcina mazei*, the hydrophobic methanophenazine plays a role similar to ubiquinone, accepting electrons from dehydrogenase enzymes and donating electrons to a heterodisulfide reductase enzyme^43^. In *P. aeruginosa*, phenazines are produced in aerobic cultures during early stationary phase, as oxygen becomes limited due to increased consumption by the high density of cells. Furthermore, PCN and PYO are known to accept electrons from dehydrogenase enzymes in the electron transport chain^44, 45^ and donate their electrons to quinones^45, 46^. Direct participation in redox reactions coupled to cellular survival could help explain the elevated intracellular concentrations found in this study. But even compared to known quinone concentrations in growing *Escherichia coli*^47^, our measurements are elevated by a factor of 5-10, suggesting further study is needed to illuminate the function of PYO in the membrane.

Octanol and water are imperfect substitutes for the complexities of the cell membrane and the cytosol, which are unlike any media used regularly in the lab ^48^. But the indication from simple, reductionist shake flask measurements that redox state is a fundamental control on phenazine lipophilicities led us to determine that simple modifications to the phenazine core structure provide microbial phenazine producers the opportunity to tune phenazine mobility strongly both within and beyond the cell. The understanding that this modulation stems from facilitation of intramolecular hydrogen bonding may help explain the profusion of phenazine analogs in different environmental niches. In addition, the suggestion from LogD measurements that reduced PYO might be retained by the membrane inspired our determination of the intracellular and intramembrane concentrations of the reduced metabolite. That it is retained primarily within the cell, despite being conventionally described as a “secreted” metabolite, both challenges our conceptualization of these compounds and points to an underappreciated physiological role for phenazines in the membrane. These results reinforce the ability of the octanol-water partition coefficient to transcend the complexity of bulk biological environments and lead to new understanding. While LogD measurements and cLogP predictions are generally used to predict environmental fate and inform pharmaceutical research, here they offer fresh insight into subcellular and extracellular metabolite localization, pointing us toward a new biological role for a molecule that has been studied for nearly 100 years^49^.

## Acknowledgments

We thank Dr. Nate Glasser, Dr. Nathan Dalleska, and the members of the Newman Lab for insightful suggestions that improved this study. Grants from the NIH (1R01AI189726-01A1) and ARO (Cooperative Agreement Number W911NF-22-2-0210) supported our research. The views and conclusions contained in this document are those of the authors and should not be interpreted as representing the official policies, either expressed or implied, of the Army Research Office or the U.S. Government. The U.S. Government is authorized to reproduce and distribute reprints for Government purposes notwithstanding any copyright notation herein. S.J.C. is grateful to Michael and Alice Jung for endowing the Jung Chair in Medicinal Chemistry and Drug Discovery at UCLA, which partially supported this work. O.G. acknowledges NIH NIGMS (R35GM137797) for funding and the Hoffman2 Cluster at UCLA Office of Advanced Research Computing’s Research Technology Group for providing computational resources (https://hprc.tamu.edu, https://www.hoffman2.idre.ucla.edu/). I.B.T. was supported by an EMBO Postdoctoral Fellowship (ALTF 191-2023).

## Notes

### Competing Interest Statement

The authors have declared no competing interest.

