## Supplementary Information for "Redox-dependent lipophilicity of phenazine metabolites is modulated by intramolecular hydrogen bonds and controls their biological distribution"

### Computational Details

All geometry optimizations of intermediates were achieved using the spin unrestricted uwb97xD [1]/cc-pvDZ [2] method, in water (dielectric constant: 78.35) and octanol (dielectric constant: 9.8629, EpsInf=2.0420) using the CPCM solvent model [3] as implemented in Gaussian16 [4]. Frequency calculations were also conducted at the same level of theory to obtain vibrational frequencies to determine the identity of the stationary points as intermediates (no imaginary frequencies), as well as obtaining the thermochemistry: enthalpy (ΔH) and free energy (ΔG) at the temperature of 298 K. All structural figures were generated with CYLview [5]. Distances in structural figures are shown in Å and energies are in kcal/mol. Single point energy corrections were carried out further with the following methods, using Truhlar's octanol solvent configuration. [6]

i) uwb97xD/6-311+g(d,p)-SMD($H_2O$)//uwb97xD/cc-pvDZ-cpcm($H_2O$)
ii) uwb97xD/6-311+g(d,p)-SMD(Octanol)//uwb97xD/cc-pvDZ-cpcm(Octanol)

Lipophilicity calculations (Log*D*) were done based on a previously established scheme, using the Gibbs free energies of the solvated molecules (Figure S2). [7, 8, 9] The energy of the intramolecular hydrogen-bond was calculated using the energy difference of the closed versus open form of each metabolite, values are given in kcal/mol. [10] The energy of the intramolecular hydrogen-bond was calculated only for structures that had defined closed versus open conformations, which were established based on the formation of a new intramolecular hydrogen-bond upon rotation of the substituent at the 1-position. The equilibrium constant ($K_{eq}$) at standard states and constant pressure was calculated to determine the isomeric ratio between open and closed forms. [11] Protonated forms of PCA and oxidized pyocyanin were used when analyzing the effects of an intramolecular hydrogen-bond. *NBO* orbital analysis [12] was conducted to compare intramolecular hydrogen-bond capabilities of the redox forms of each metabolite. [13] The strength of the *NBO* interaction was used to rationalize the lipophilicity of the specific redox form.

$$E_{HB} = E_{closed} - E_{open}$$

**Figure S1**. Intramolecular hydrogen-bond calculation scheme. Electronic Energies were used, with all values reported in kcal/mol.

$$\mathrm{Log}D = \frac{(\Delta G_{water} - \Delta G_{octanol})}{(2.303RT)}$$

**Figure S2**. Calculation scheme used to measure lipophilicity, using Gibbs free energy of the solvated metabolite, at room temperature. Value of *R* used was $1.987\times10^{-3}$ kcal/mol.

$$\Delta \mathrm{Log}D = \mathrm{Log}D_{closed} - \mathrm{Log}D_{open}$$

**Figure S3**. Differences in Log*D* due to the intramolecular hydrogen bond measured using the above scheme.

$$\Delta E_{HB} = E_{HB}\mathrm{octanol} - E_{HB}\mathrm{water}$$

**Figure S4**. Differences in intramolecular hydrogen bond strength due to changes in solvent.

$$K_{eq} = \frac{-(G^{o}_{closed} - G^{o}_{open})}{(RT)}$$

**Figure S5**. Calculation scheme used to measure equilibrium constant, using Gibbs free energy of the solvated metabolite, at room temperature. Value of *R* used was $1.987\times10^{-3}$ kcal/mol.

| Metabolite | $E_{HB}$ (kcal/mol) | $\Delta E_{HB}$ (kcal/mol) | $K_{eq}$ |
|---|---|---|---|
| $PCN_{oxidized}$ (water) | -1.588225 | *xx* | 2.373364 |
| $PCN_{oxidized}$ (octanol) | -3.027731 | -1.439506 | 25.862847 |
| $PCN_{reduced}$ (water) | -2.491838 | *xx* | 95.368043 |
| $PCN_{reduced}$ (octanol) | -2.574042 | -0.082204 | 111.79018 |
| $PCA_{reduced-protonated}$ (water) | -6.199789 | *xx* | 5755.5051 |
| $PCA_{reduced-protonated}$ (octanol) | -7.518813 | -1.319024 | 46376.067 |
| $PCA_{oxidized-protonated}$ (water) | -5.312491 | *xx* | 1759.2903 |
| $PCA_{oxidized-protonated}$ (octanol) | -5.706567 | -0.394076 | 3339.1868 |
| $Pyocyanin_{oxidized-protonated}$ (water) | -1.1401839 | *xx* | 5.951809 |
| $Pyocyanin_{oxidized-protonated}$ (octanol) | -1.9189225 | -0.7787386 | 21.807981 |
| 1-OH $Phenazine_{oxidized}$ (water) | -1.5386521 | *xx* | 8.920126 |
| 1-OH $Phenazine_{oxidized}$ (octanol) | -2.7227616 | -1.1841095 | 63.430893 |

**Table S1**. Intramolecular hydrogen bond energies, $\Delta E_{HB}$, and equilibrium constant ($K_{eq}$) calculated at the uwb97xD/6-311+g(d,p)-SMD(solvent)//uwb97xD/cc-pvDZ-cpcm(solvent) level of theory, according to the equations depicted above.

| Metabolite | LogD | ΔLogD |
|---|---|---|
| $PCN_{oxidized}$ (open) | -1.002651 | *xx* |
| $PCN_{oxidized}$ (closed) | 0.0344949 | 1.03714598 |

| | | |
|---|---|---|
| $PCN_{reduced}$ (open) | 0.596071 | *xx* |
| $PCN_{reduced}$ (closed) | 0.665601 | 0.06898975 |
| $PCA_{reduced\text{-}protonated}$ (open) | 0.375764 | *xx* |
| $PCA_{reduced\text{-}protonated}$ (closed) | 1.28183 | 0.90606544 |
| $PCA_{reduced}$ | -4.235971 | *xx* |
| $PCA_{oxidized\text{-}protonated}$ (open) | -0.321952 | *xx* |
| $PCA_{oxidized\text{-}protonated}$ (closed) | -0.043694 | 0.27825868 |
| $PCA_{oxidized}$ | -6.372354 | *xx* |
| $Pyocyanin_{reduced}$ | 1.74498086 | *xx* |
| $Pyocyanin_{oxidized\text{-}protonated}$ (open) | -1.1866238 | *xx* |
| $Pyocyanin_{oxidized\text{-}protonated}$ (closed) | -0.6227475 | 0.56387626 |
| $Pyocyanin_{oxidized}$ | -0.0625507 | *xx* |
| 1-OH $Phenazine_{reduced}$ | 1.11855389 | *xx* |
| 1-OH $Phenazine_{oxidized}$ (open) | -0.2000703 | *xx* |
| 1-OH $Phenazine_{oxidized}$ (closed) | 0.651722321 | 0.8517935 |

**Table S2**. LogD and ΔLogD calculated at the uwb97xD/6-311+g(d,p)-SMD(solvent)//uwb97xD/cc-pvDZ-cpcm(solvent) level of theory, according to the equations depicted above at neutral pH.

| Metabolite | NBO Interaction Energy (kcal/mol) |
|---|---|
| $PCN_{oxidized}$ | 7.36 |
| $PCN_{reduced}$ | 7.67 |
| $PCA_{reduced}$ | 15.66 |
| $PCA_{oxidized}$ | *no interaction* |

| | |
|---|---|
| $Pyocyanin_{reduced}$ | 0.71 |
| $Pyocyanin_{oxidized}$ | *no interaction* |
| 1-OH $Phenazine_{reduced}$ | 0.65 |
| 1-OH $Phenazine_{oxidized}$ | 3.05 |

**Table S3**. Intramolecular hydrogen bonding strength calculated using *NBO* analysis at the uwb97xD/cc-pvDZ-cpcm($H_2O$) level of theory.

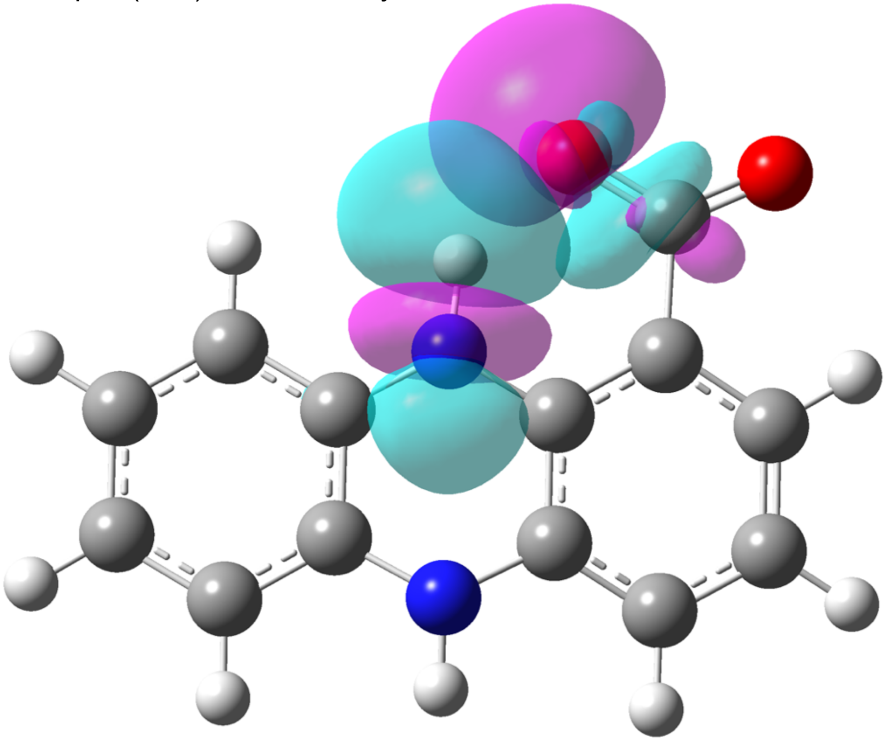

**Figure S6**. Intramolecular hydrogen bonding within $PCA_{reduced}$ shown using *NBO* orbitals, generated from the oxygen lone pair, calculated at the uwb97xD/cc-pvDZ-cpcm($H_2O$) level of theory.

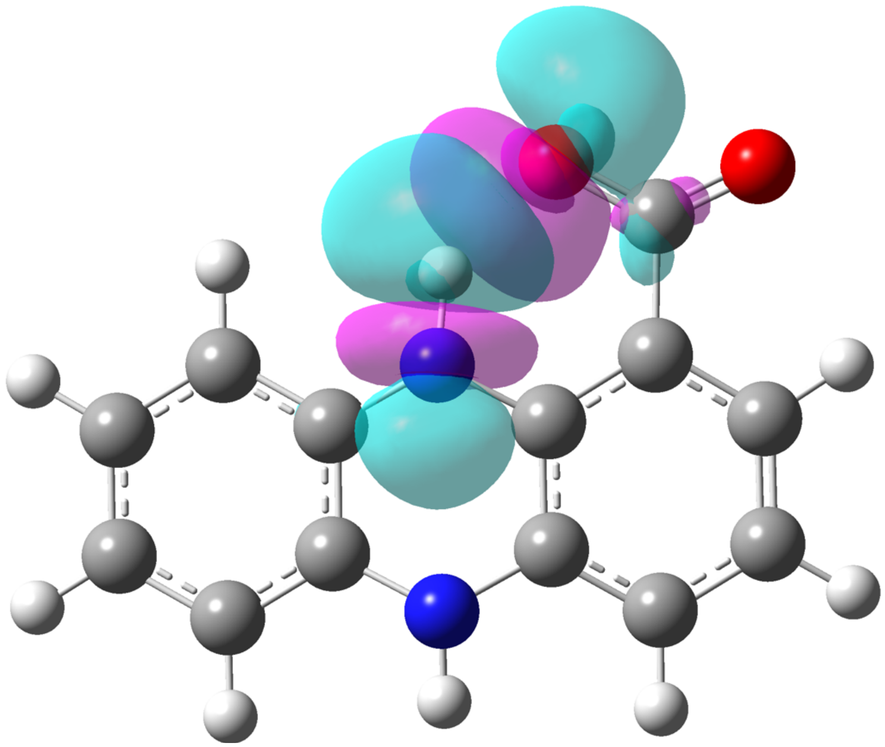

**Figure S7**. Intramolecular hydrogen bonding within $PCA_{reduced}$ shown using *NBO* orbitals, generated from the carbonyl π-bond, calculated at the uwb97xD/cc-pvDZ-cpcm($H_2O$) level of theory.

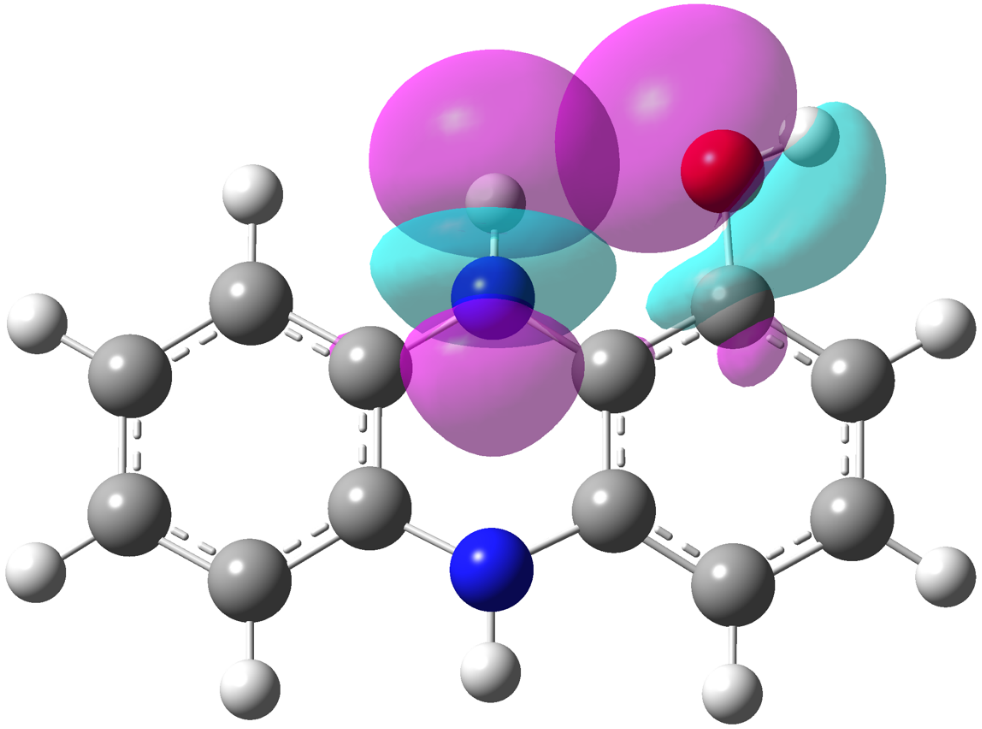

**Figure S8**. Intramolecular hydrogen bonding within OH-$Phenazine_{reduced}$ shown using *NBO* orbitals, calculated at the uwb97xD/cc-pvDZ-cpcm($H_2O$) level of theory.

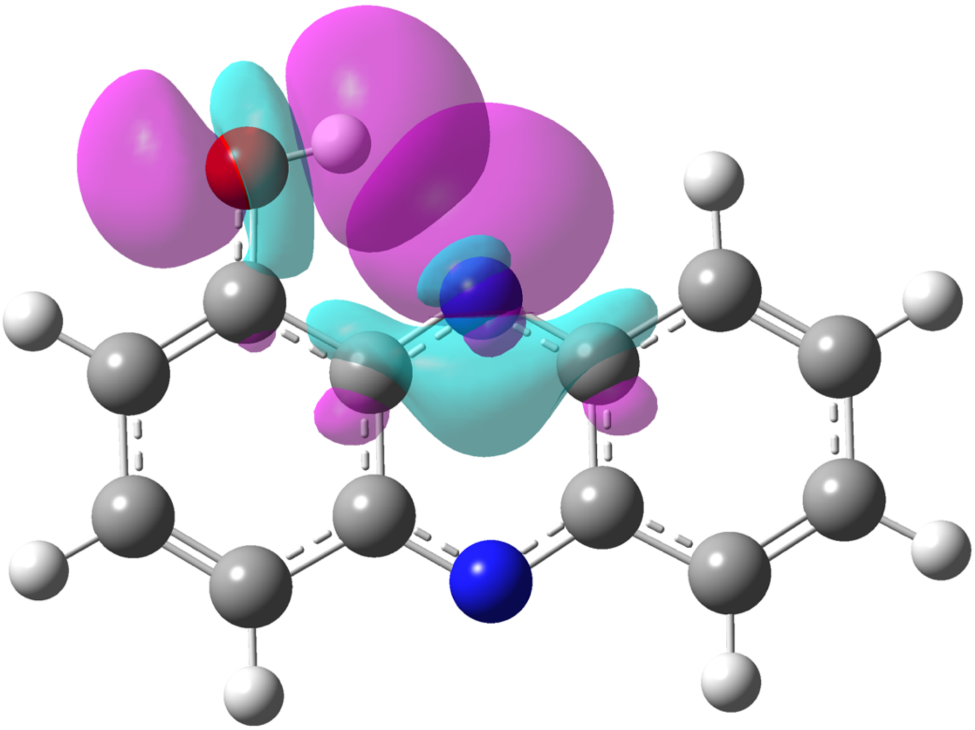

**Figure S9**. Intramolecular hydrogen bonding within OH-Phenazine$_{oxidized}$ shown using *NBO* orbitals, calculated at the uwb97xD/cc-pvDZ-cpcm($H_2O$) level of theory.

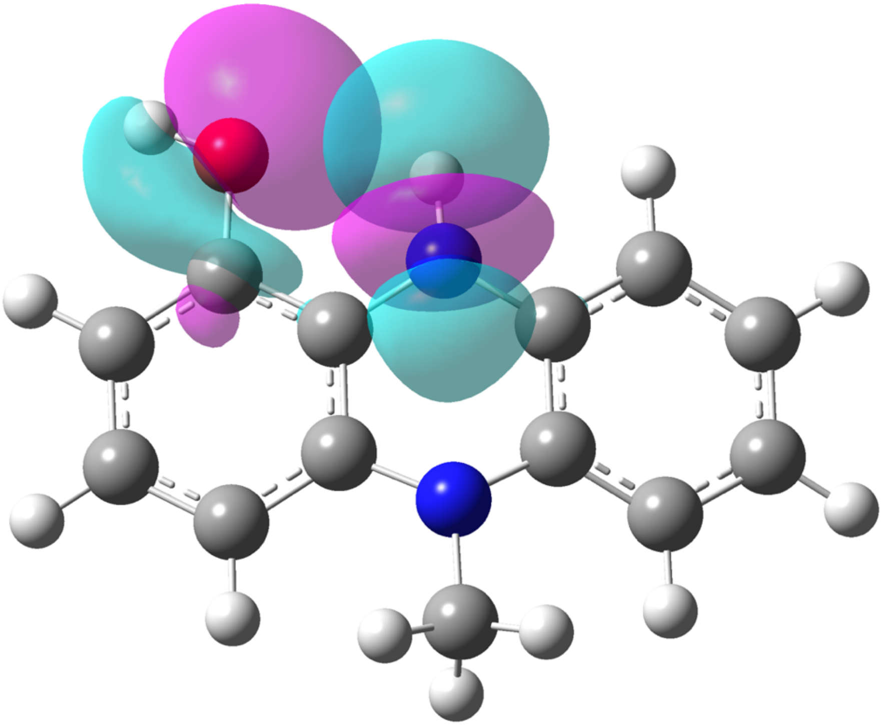

**Figure S10**. Intramolecular hydrogen bonding within Pyocyanin$_{reduced}$ shown using *NBO* orbitals, calculated at the uwb97xD/cc-pvDZ-cpcm($H_2O$) level of theory.

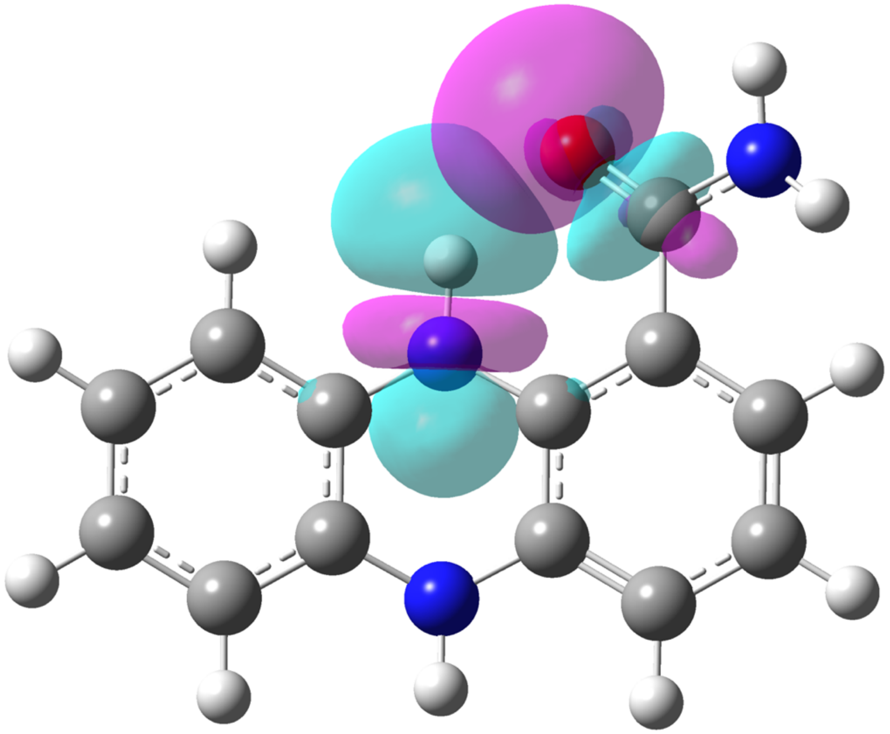

**Figure S11**. Intramolecular hydrogen bonding within $PCN_{reduced}$ shown using *NBO* orbitals, generated from the oxygen lone pair, calculated at the uwb97xD/cc-pvDZ-cpcm($H_2O$) level of theory.

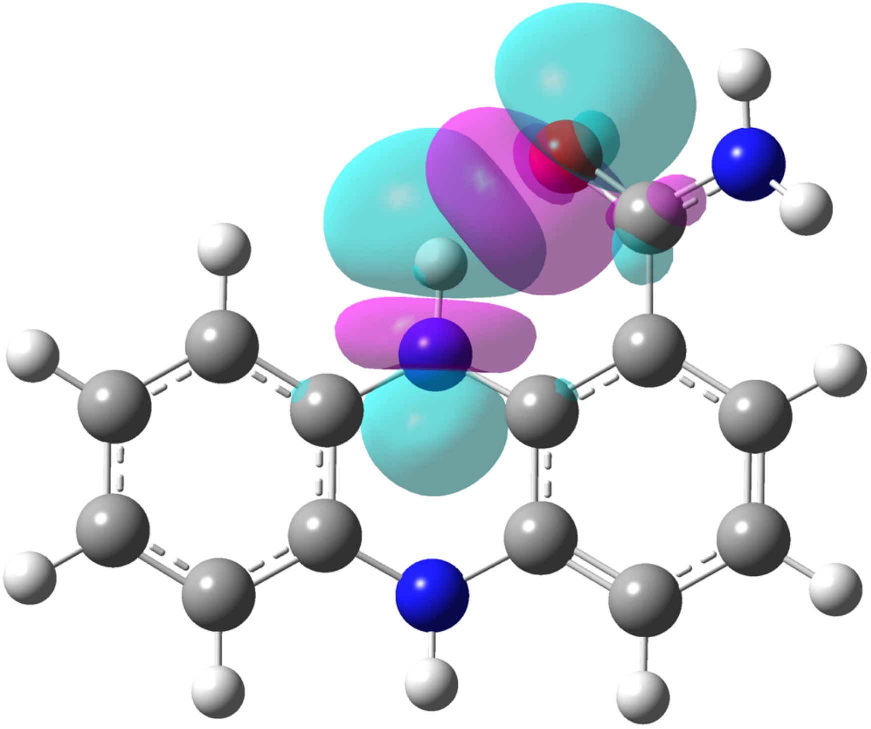

**Figure S12**. Intramolecular hydrogen bonding within $PCN_{reduced}$ shown using *NBO* orbitals, generated from the carbonyl π-bond, calculated at the uwb97xD/cc-pvDZ-cpcm($H_2O$) level of theory.

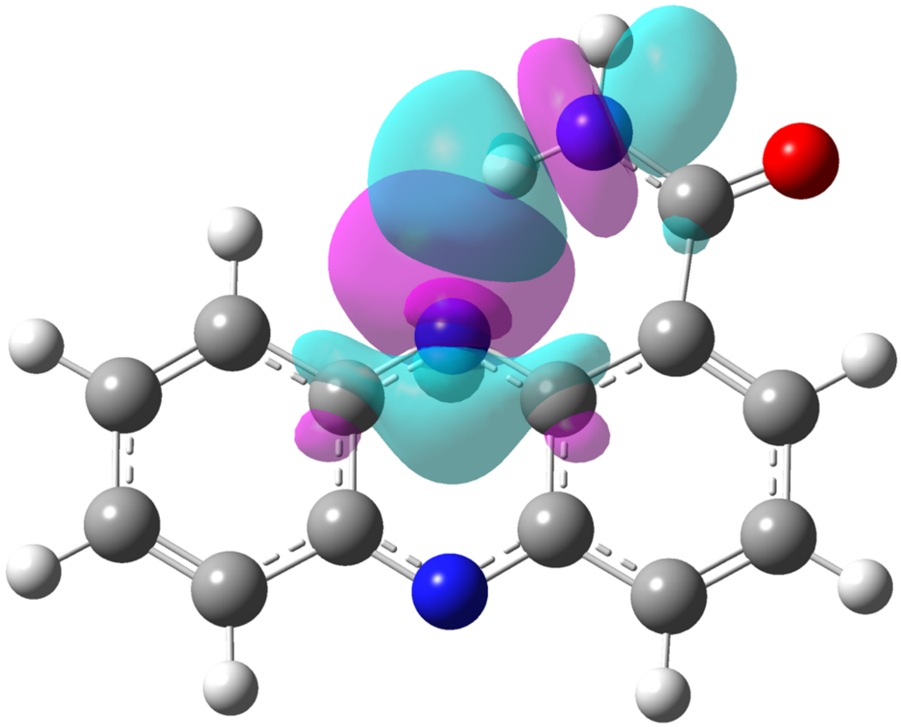

**Figure S13**. Intramolecular hydrogen bonding within $PCN_{oxidized}$ shown using *NBO* orbitals, calculated at the uwb97xD/cc-pvDZ-cpcm($H_2O$) level of theory.

**Table S4.** Cartesian coordinates (xyz format) and energies of all the structures involved in each reaction mechanism studied calculated at the uwb97xD//cc-pvDZ-cpcm(solvent) level of theory.

**PCN_reduced_open**
E(scf) = -741.344857226 a.u.

| | | | | | | | |
|---|---|---|---|---|---|---|---|
| C | 4.265822 | -0.133093 | 0.340205 | C | -2.947864 | 1.060149 | 0.254109 |
| C | 3.389019 | 0.924961 | 0.073929 | C | -2.096846 | -0.039801 | 0.034268 |
| C | 2.426911 | -1.688854 | 0.170160 | C | -0.726498 | 0.172012 | -0.184989 |
| C | 3.786399 | -1.436961 | 0.389139 | H | -0.688131 | 3.583336 | -0.010593 |
| H | 5.322723 | 0.075121 | 0.512800 | H | -3.121051 | 3.201360 | 0.388692 |
| H | 3.755852 | 1.953264 | 0.042483 | H | -4.003768 | 0.861611 | 0.436758 |
| H | 2.040146 | -2.709517 | 0.212753 | N | 0.195316 | -0.860491 | -0.394234 |
| H | 4.460565 | -2.267972 | 0.600920 | N | 1.128684 | 1.690304 | -0.484629 |
| C | 2.040658 | 0.676385 | -0.164317 | H | -0.142493 | -1.793648 | -0.182225 |
| C | 1.556077 | -0.642874 | -0.116658 | H | 1.466877 | 2.636903 | -0.359613 |
| C | -0.231976 | 1.498480 | -0.207046 | C | -2.741660 | -1.393823 | 0.080776 |
| C | -1.090898 | 2.568235 | 0.004977 | O | -3.703161 | -1.622773 | 0.801994 |
| C | -2.456396 | 2.351576 | 0.230176 | N | -2.189927 | -2.365130 | -0.712545 |

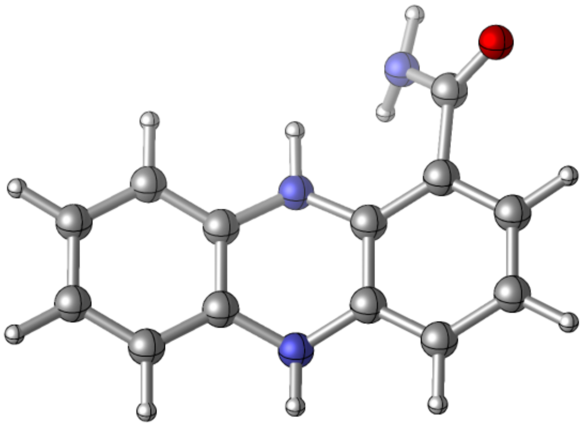

Zero-point correction= 0.222875 (Hartree/Particle)
Thermal correction to Energy= 0.235843
Thermal correction to Enthalpy= 0.236787
Thermal correction to Gibbs Free Energy= 0.183369
Sum of electronic and zero-point Energies= -741.121982
Sum of electronic and thermal Energies= -741.109014
Sum of electronic and thermal Enthalpies= -741.108070
Sum of electronic and thermal Free Energies= -741.161489

**PCN_reduced_closed**
E(scf) = -741.350279331 a.u.

| | | | | | | | |
|---|---|---|---|---|---|---|---|
| C | 4.271787 | -0.145413 | 0.354472 | C | -2.965827 | 1.027005 | 0.131204 |
| C | 3.391967 | 0.929385 | 0.173050 | C | -2.081468 | -0.065234 | -0.027110 |
| C | 2.441989 | -1.688515 | 0.027962 | C | -0.710059 | 0.186012 | -0.215425 |
| C | 3.798778 | -1.450056 | 0.280834 | H | -0.732275 | 3.590127 | 0.087019 |
| H | 5.325847 | 0.051139 | 0.555935 | H | -3.182778 | 3.158784 | 0.284093 |
| H | 3.755125 | 1.957840 | 0.232459 | H | -4.039620 | 0.852101 | 0.201404 |
| H | 2.058485 | -2.709739 | -0.023069 | N | 0.214556 | -0.813612 | -0.468596 |
| H | 4.476039 | -2.293488 | 0.422821 | N | 1.137292 | 1.730039 | -0.354673 |
| C | 2.048551 | 0.696658 | -0.102001 | H | -0.180227 | -1.753239 | -0.474568 |
| C | 1.567578 | -0.625031 | -0.171959 | H | 1.470185 | 2.669043 | -0.173273 |
| C | -0.231908 | 1.520595 | -0.154592 | C | -2.578933 | -1.472480 | -0.005937 |
| C | -1.115625 | 2.568270 | 0.047005 | O | -1.901711 | -2.427305 | -0.406402 |
| C | -2.492721 | 2.322897 | 0.166460 | N | -3.835571 | -1.673839 | 0.463875 |

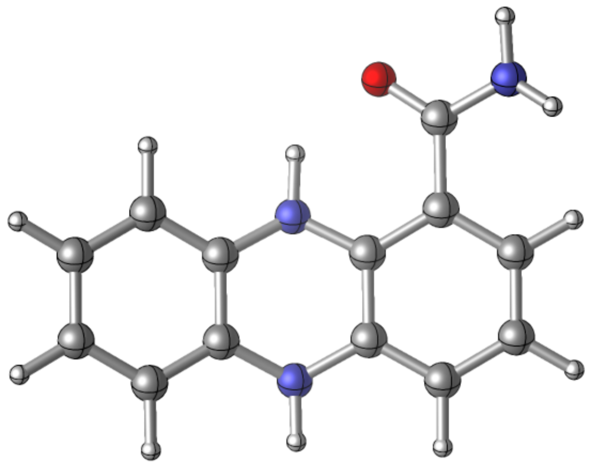

Zero-point correction= 0.222637 (Hartree/Particle)
Thermal correction to Energy= 0.235668
Thermal correction to Enthalpy= 0.236612
Thermal correction to Gibbs Free Energy= 0.183037
Sum of electronic and zero-point Energies= -741.127643
Sum of electronic and thermal Energies= -741.114611
Sum of electronic and thermal Enthalpies= -741.113667
Sum of electronic and thermal Free Energies= -741.167242

**PCN_reduced_open_octanol**
E(scf) = -741.343252018 a.u.

| | | | | | | | |
|---|---|---|---|---|---|---|---|
| C | 4.266249 | -0.132813 | 0.339122 | C | -2.947672 | 1.060104 | 0.255271 |
| C | 3.389359 | 0.924834 | 0.072322 | C | -2.096826 | -0.039698 | 0.035043 |
| C | 2.427298 | -1.688224 | 0.171316 | C | -0.726489 | 0.172047 | -0.184274 |
| C | 3.786764 | -1.436441 | 0.389526 | H | -0.688351 | 3.583098 | -0.011319 |
| H | 5.323217 | 0.075440 | 0.511061 | H | -3.120785 | 3.201125 | 0.390088 |
| H | 3.756466 | 1.953048 | 0.039799 | H | -4.003182 | 0.860831 | 0.439111 |
| H | 2.040517 | -2.708852 | 0.215237 | N | 0.195387 | -0.860931 | -0.392746 |
| H | 4.460926 | -2.267257 | 0.601926 | N | 1.128755 | 1.690227 | -0.485153 |
| C | 2.040914 | 0.676366 | -0.164856 | H | -0.142689 | -1.792873 | -0.175788 |
| C | 1.556279 | -0.642675 | -0.116010 | H | 1.466690 | 2.636588 | -0.358573 |
| C | -0.232087 | 1.498275 | -0.207066 | C | -2.742847 | -1.393074 | 0.080304 |
| C | -1.091009 | 2.567905 | 0.004758 | O | -3.710140 | -1.620595 | 0.792477 |
| C | -2.456224 | 2.351409 | 0.230973 | N | -2.183770 | -2.368368 | -0.706543 |

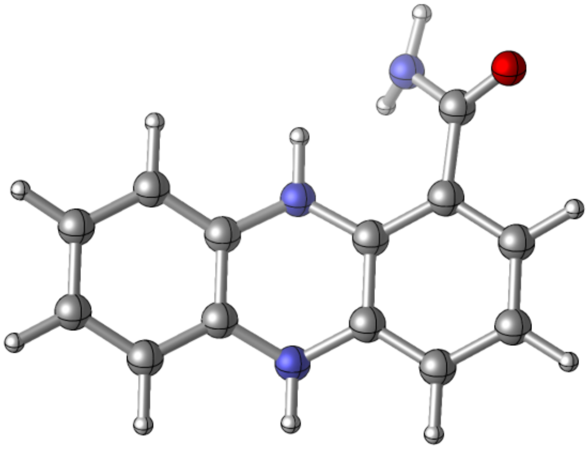

Zero-point correction= 0.222925 (Hartree/Particle)
Thermal correction to Energy= 0.235885
Thermal correction to Enthalpy= 0.236829
Thermal correction to Gibbs Free Energy= 0.183435
Sum of electronic and zero-point Energies= -741.120327
Sum of electronic and thermal Energies= -741.107367
Sum of electronic and thermal Enthalpies= -741.106423
Sum of electronic and thermal Free Energies= -741.159817

**PCN_reduced_closed_octanol**
E(scf) = -741.348848925 a.u.

| | | | | | | | |
|---|---|---|---|---|---|---|---|
| C | 4.274051 | -0.144341 | 0.348602 | C | -2.967497 | 1.026370 | 0.125983 |
| C | 3.393058 | 0.930094 | 0.171457 | C | -2.082095 | -0.065798 | -0.028023 |
| C | 2.444131 | -1.687691 | 0.025565 | C | -0.710129 | 0.185518 | -0.211966 |
| C | 3.801461 | -1.448927 | 0.274436 | H | -0.734993 | 3.589978 | 0.086706 |
| H | 5.328641 | 0.052440 | 0.546852 | H | -3.185902 | 3.158143 | 0.275785 |
| H | 3.756130 | 1.958619 | 0.230817 | H | -4.041483 | 0.850700 | 0.191334 |
| H | 2.060902 | -2.708954 | -0.025912 | N | 0.214580 | -0.813973 | -0.460438 |
| H | 4.479601 | -2.292202 | 0.412830 | N | 1.136565 | 1.730677 | -0.346972 |
| C | 2.048979 | 0.697134 | -0.099261 | H | -0.180011 | -1.753662 | -0.470509 |
| C | 1.568394 | -0.624565 | -0.169557 | H | 1.469130 | 2.669571 | -0.165572 |
| C | -0.232784 | 1.520685 | -0.150992 | C | -2.578317 | -1.472963 | -0.005634 |
| C | -1.117649 | 2.567820 | 0.046771 | O | -1.898674 | -2.429172 | -0.396416 |
| C | -2.495249 | 2.322250 | 0.161805 | N | -3.840234 | -1.673359 | 0.454903 |

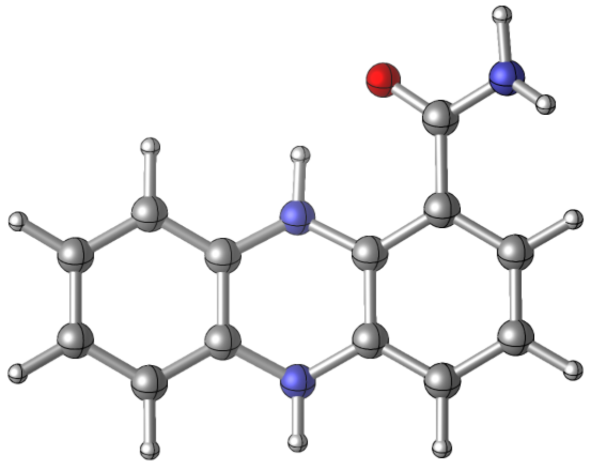

Zero-point correction= 0.222673 (Hartree/Particle)
Thermal correction to Energy= 0.235698
Thermal correction to Enthalpy= 0.236643
Thermal correction to Gibbs Free Energy= 0.183084
Sum of electronic and zero-point Energies= -741.126176
Sum of electronic and thermal Energies= -741.113150
Sum of electronic and thermal Enthalpies= -741.112206
Sum of electronic and thermal Free Energies= -741.165764

**PCN_oxidized_open**
E(scf) = -740.117781549 a.u.

| | | | | | | | |
|---|---|---|---|---|---|---|---|
| C | 4.268597 | -0.149452 | -0.057849 | C | -2.902959 | 1.040718 | 0.022377 |
| C | 3.418177 | 0.917723 | -0.016329 | C | -2.065135 | -0.042198 | 0.035074 |
| C | 2.419715 | -1.728407 | -0.081988 | C | -0.640245 | 0.163139 | 0.028671 |
| C | 3.763577 | -1.488011 | -0.092264 | H | -0.636280 | 3.615611 | 0.053943 |
| H | 5.347645 | 0.011511 | -0.066259 | H | -3.098568 | 3.209574 | 0.034698 |
| H | 3.780517 | 1.946314 | 0.008074 | H | -3.983520 | 0.886673 | 0.022691 |
| H | 2.011973 | -2.739829 | -0.106099 | N | 0.181388 | -0.890543 | -0.019334 |
| H | 4.467787 | -2.320652 | -0.126566 | N | 1.180302 | 1.763172 | 0.022955 |
| C | 2.002308 | 0.707369 | -0.005385 | C | -2.625609 | -1.439817 | 0.130561 |
| C | 1.495221 | -0.636637 | -0.034648 | O | -2.359024 | -2.185444 | 1.062767 |
| C | -0.134069 | 1.508387 | 0.035333 | N | -3.481903 | -1.767981 | -0.863254 |
| C | -1.051504 | 2.607128 | 0.044139 | H | -3.895894 | -2.692300 | -0.867740 |
| C | -2.395302 | 2.375964 | 0.032432 | H | -3.596606 | -1.181321 | -1.678176 |

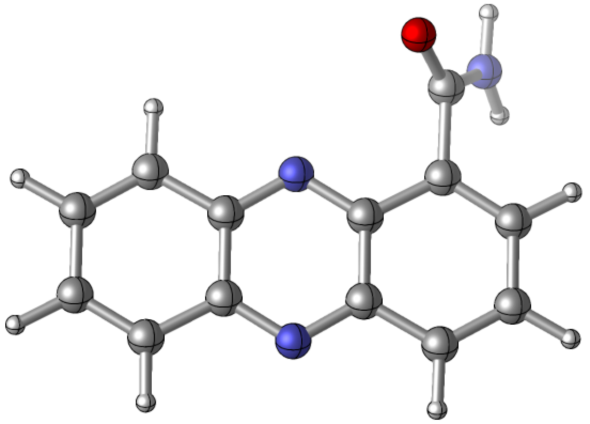

Zero-point correction= 0.198600 (Hartree/Particle)
Thermal correction to Energy= 0.211223
Thermal correction to Enthalpy= 0.212167
Thermal correction to Gibbs Free Energy= 0.159034
Sum of electronic and zero-point Energies= -739.919182
Sum of electronic and thermal Energies= -739.906558
Sum of electronic and thermal Enthalpies= -739.905614
Sum of electronic and thermal Free Energies= -739.958748

**PCN_oxidized_closed**
E(scf) = -740.126441754 a.u.

| | | | | | | | |
|---|---|---|---|---|---|---|---|
| C | 4.274329 | -0.080236 | -0.000040 | C | -2.913119 | 1.024198 | 0.000075 |
| C | 3.405623 | 0.972809 | -0.000105 | C | -2.086065 | -0.069045 | 0.000015 |
| C | 2.451906 | -1.692672 | 0.000151 | C | -0.655906 | 0.148648 | 0.000018 |
| C | 3.791522 | -1.427083 | 0.000094 | H | -0.663274 | 3.606340 | -0.000059 |
| H | 5.350509 | 0.098160 | -0.000088 | H | -3.129852 | 3.189046 | 0.000082 |
| H | 3.748822 | 2.008072 | -0.000203 | H | -3.987589 | 0.839869 | 0.000126 |
| H | 2.066064 | -2.712973 | 0.000247 | N | 0.200789 | -0.880145 | 0.000102 |
| H | 4.510003 | -2.248099 | 0.000147 | N | 1.150390 | 1.773560 | -0.000080 |
| C | 1.994351 | 0.735985 | -0.000043 | C | -2.777180 | -1.420415 | -0.000001 |
| C | 1.512233 | -0.614895 | 0.000073 | O | -4.005661 | -1.484062 | 0.000089 |
| C | -0.159736 | 1.501576 | -0.000032 | N | -1.990017 | -2.509700 | -0.000277 |
| C | -1.078581 | 2.597968 | -0.000020 | H | -2.432034 | -3.419644 | -0.000376 |
| C | -2.420738 | 2.360587 | 0.000053 | H | -0.977340 | -2.398828 | -0.000227 |

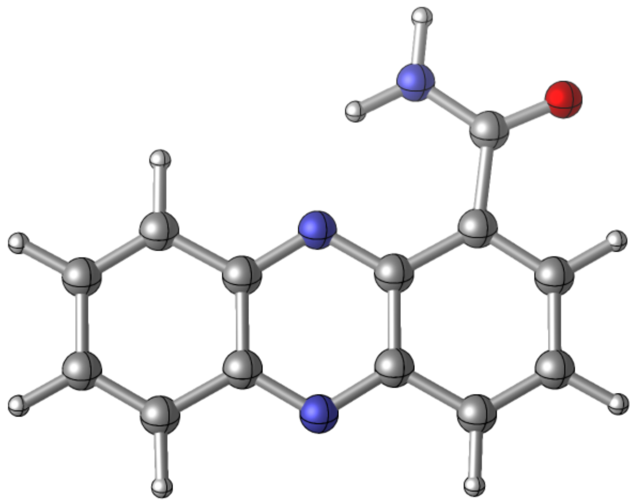

Zero-point correction= 0.199481 (Hartree/Particle)
Thermal correction to Energy= 0.211626
Thermal correction to Enthalpy= 0.212570
Thermal correction to Gibbs Free Energy= 0.160749
Sum of electronic and zero-point Energies= -739.926961
Sum of electronic and thermal Energies= -739.914816
Sum of electronic and thermal Enthalpies= -739.913872
Sum of electronic and thermal Free Energies= -739.965693

**PCN_oxidized_open_octanol**
E(scf) = -740.116075096 a.u.

| | | | | | | | |
|---|---|---|---|---|---|---|---|
| C | 4.267756 | -0.150208 | 0.057228 | C | -2.902790 | 1.041062 | -0.021913 |
| C | 3.417806 | 0.917255 | 0.015963 | C | -2.064909 | -0.041713 | -0.034938 |
| C | 2.418525 | -1.728594 | 0.081733 | C | -0.640150 | 0.163769 | -0.028284 |
| C | 3.762360 | -1.488591 | 0.091631 | H | -0.635517 | 3.615748 | -0.053518 |
| H | 5.346877 | 0.010451 | 0.065289 | H | -3.098166 | 3.210021 | -0.034164 |
| H | 3.779943 | 1.945870 | -0.008593 | H | -3.983261 | 0.886324 | -0.022942 |
| H | 2.009676 | -2.739514 | 0.105364 | N | 0.180803 | -0.890228 | 0.020080 |
| H | 4.466300 | -2.321497 | 0.125401 | N | 1.180502 | 1.763572 | -0.022629 |
| C | 2.001887 | 0.707407 | 0.005304 | C | -2.624526 | -1.439793 | -0.131907 |
| C | 1.494464 | -0.636481 | 0.034743 | O | -2.361952 | -2.182750 | -1.065576 |
| C | -0.133809 | 1.508827 | -0.034985 | N | -3.482142 | -1.768677 | 0.863265 |
| C | -1.051239 | 2.607508 | -0.043547 | H | -3.876435 | -2.701520 | 0.875113 |
| C | -2.395024 | 2.376248 | -0.031721 | H | -3.570053 | -1.196723 | 1.691812 |

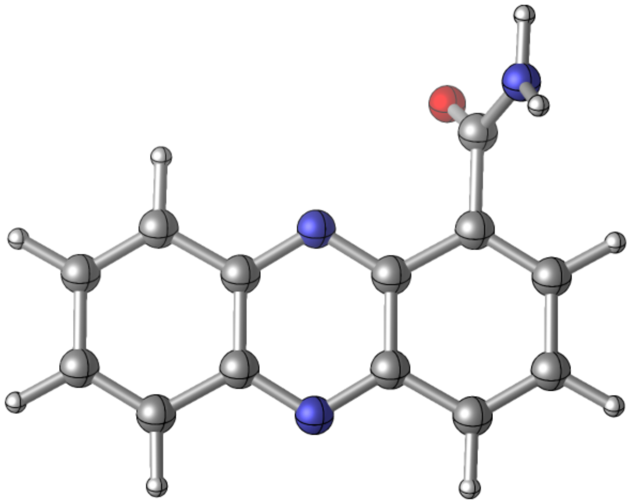

Zero-point correction= 0.198606 (Hartree/Particle)
Thermal correction to Energy= 0.211228
Thermal correction to Enthalpy= 0.212172
Thermal correction to Gibbs Free Energy= 0.159018
Sum of electronic and zero-point Energies= -739.917469
Sum of electronic and thermal Energies= -739.904847
Sum of electronic and thermal Enthalpies= -739.903903
Sum of electronic and thermal Free Energies= -739.957057

**PCN_oxidized_closed_octanol**
E(scf) = -740.125300697 a.u.

| | | | | | | | |
|---|---|---|---|---|---|---|---|
| C | 4.274595 | -0.080572 | -0.000088 | C | -2.912600 | 1.024532 | 0.000054 |
| C | 3.405751 | 0.972320 | -0.000033 | C | -2.085925 | -0.068845 | 0.000093 |
| C | 2.452154 | -1.692772 | -0.000106 | C | -0.655772 | 0.148320 | 0.000065 |
| C | 3.791772 | -1.427300 | -0.000125 | H | -0.661399 | 3.605951 | 0.000073 |
| H | 5.350784 | 0.097963 | -0.000102 | H | -3.128678 | 3.189446 | 0.000020 |
| H | 3.748259 | 2.007798 | -0.000003 | H | -3.987013 | 0.839586 | 0.000025 |
| H | 2.066248 | -2.713055 | -0.000134 | N | 0.200828 | -0.880606 | -0.000010 |
| H | 4.510135 | -2.248469 | -0.000170 | N | 1.150841 | 1.773244 | 0.000035 |
| C | 1.994521 | 0.735469 | -0.000012 | C | -2.778760 | -1.419709 | 0.000007 |
| C | 1.512307 | -0.615154 | -0.000041 | O | -4.006391 | -1.482046 | -0.000794 |
| C | -0.159106 | 1.501308 | 0.000065 | N | -1.991868 | -2.510076 | 0.000841 |
| C | -1.077623 | 2.597972 | 0.000076 | H | -2.436049 | -3.418845 | -0.000017 |
| C | -2.419725 | 2.360793 | 0.000057 | H | -0.979303 | -2.400113 | 0.000510 |

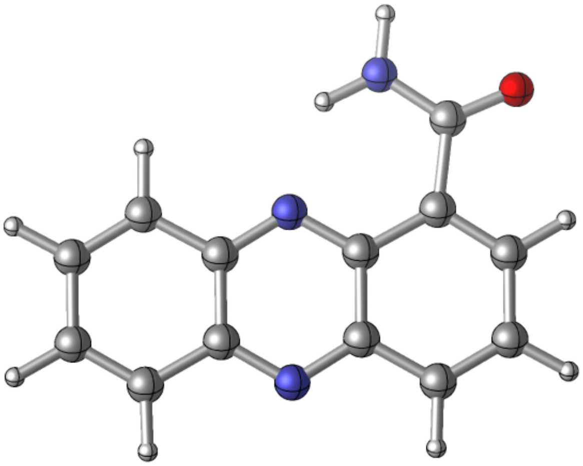

Zero-point correction= 0.199512 (Hartree/Particle)
Thermal correction to Energy= 0.211659
Thermal correction to Enthalpy= 0.212603
Thermal correction to Gibbs Free Energy= 0.160772
Sum of electronic and zero-point Energies= -739.925789
Sum of electronic and thermal Energies= -739.913642
Sum of electronic and thermal Enthalpies= -739.912698
Sum of electronic and thermal Free Energies= -739.964528

**PCA_reduced_open**
E(scf) = -761.201341105 a.u.

| | | | | | | | |
|---|---|---|---|---|---|---|---|
| C | -4.259925 | -0.088303 | 0.255285 | C | 3.011883 | 0.981990 | 0.192962 |
| C | -3.356999 | 0.973941 | 0.126341 | C | 2.108781 | -0.085982 | 0.015653 |
| C | -2.454687 | -1.657321 | -0.078580 | C | 0.744306 | 0.172370 | -0.179179 |
| C | -3.811877 | -1.399637 | 0.150225 | H | 0.805651 | 3.566147 | 0.185254 |
| H | -5.314097 | 0.122902 | 0.440183 | H | 3.244749 | 3.107639 | 0.421218 |
| H | -3.702333 | 2.006789 | 0.207980 | H | 4.071840 | 0.751575 | 0.298147 |
| H | -2.090671 | -2.684367 | -0.151800 | N | -0.196474 | -0.806090 | -0.494327 |
| H | -4.508306 | -2.233158 | 0.250193 | N | -1.077419 | 1.746331 | -0.332748 |
| C | -2.012237 | 0.723921 | -0.128104 | C | 2.689585 | -1.460518 | 0.020798 |
| C | -1.559125 | -0.604380 | -0.225785 | O | 3.781918 | -1.730642 | -0.415763 |
| C | 0.282712 | 1.509525 | -0.109398 | O | 1.942298 | -2.446330 | 0.570087 |
| C | 1.179447 | 2.541682 | 0.128672 | H | 1.223262 | -2.067673 | 1.098089 |
| C | 2.551178 | 2.281831 | 0.261223 | H | 0.113571 | -1.757666 | -0.635296 |

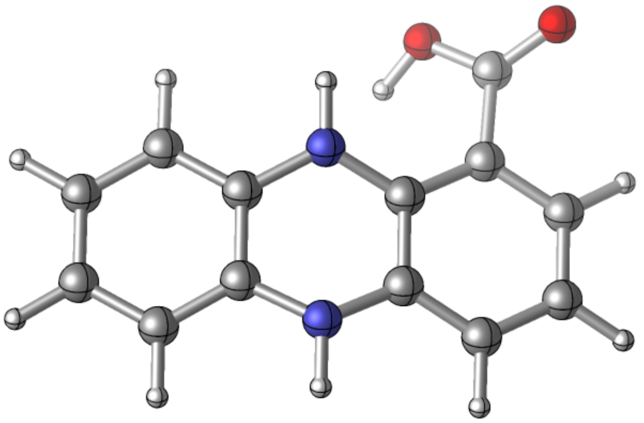

Zero-point correction= 0.209784 (Hartree/Particle)
Thermal correction to Energy= 0.222886
Thermal correction to Enthalpy= 0.223830
Thermal correction to Gibbs Free Energy= 0.169523
Sum of electronic and zero-point Energies= -760.991557
Sum of electronic and thermal Energies= -760.978455
Sum of electronic and thermal Enthalpies= -760.977511
Sum of electronic and thermal Free Energies= -761.031818

**PCA_reduced_closed**
E(scf) = -761.215996897 a.u.

| | | | | | | | |
|---|---|---|---|---|---|---|---|
| C | 4.298767 | -0.126822 | -0.257468 | C | -2.976924 | 1.017332 | -0.174335 |
| C | 3.406658 | 0.939664 | -0.090769 | C | -2.086331 | -0.070028 | -0.007978 |
| C | 2.465527 | -1.684020 | -0.044680 | C | -0.708871 | 0.169411 | 0.155239 |
| C | 3.830515 | -1.434971 | -0.233802 | H | -0.742647 | 3.576494 | -0.081894 |
| H | 5.359467 | 0.079086 | -0.408113 | H | -3.186421 | 3.148688 | -0.332177 |
| H | 3.766727 | 1.970597 | -0.111561 | H | -4.040248 | 0.812877 | -0.284817 |
| H | 2.085666 | -2.707797 | -0.031892 | N | 0.212111 | -0.838047 | 0.360457 |
| H | 4.517411 | -2.272187 | -0.364867 | N | 1.129272 | 1.722215 | 0.344869 |
| C | 2.052725 | 0.696330 | 0.117305 | C | -2.596686 | -1.453629 | -0.007818 |
| C | 1.578066 | -0.628787 | 0.137250 | O | -1.917281 | -2.461508 | 0.131264 |
| C | -0.235963 | 1.508374 | 0.130144 | O | -3.921116 | -1.536058 | -0.183638 |
| C | -1.124180 | 2.553374 | -0.058754 | H | -4.146620 | -2.479924 | -0.168227 |
| C | -2.502192 | 2.310148 | -0.200657 | | | | |

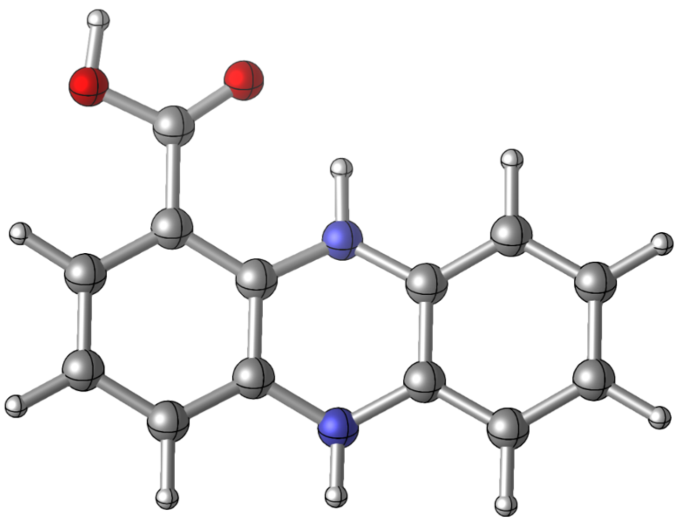

Zero-point correction= 0.210624 (Hartree/Particle)
Thermal correction to Energy= 0.223276
Thermal correction to Enthalpy= 0.224220
Thermal correction to Gibbs Free Energy= 0.171229
Sum of electronic and zero-point Energies= -761.005373
Sum of electronic and thermal Energies= -760.992721
Sum of electronic and thermal Enthalpies= -760.991777
Sum of electronic and thermal Free Energies= -761.044768

**PCA_oxidized_open**
E(scf) = -759.980243658 a.u.

| | | | | | | | |
|---|---|---|---|---|---|---|---|
| C | -4.289817 | -0.130663 | 0.032770 | C | 2.890471 | 1.028428 | 0.010995 |
| C | -3.432029 | 0.929998 | -0.019327 | C | 2.055414 | -0.060240 | -0.006310 |
| C | -2.451764 | -1.721637 | 0.105607 | C | 0.625712 | 0.140980 | -0.013633 |
| C | -3.793921 | -1.471652 | 0.097513 | H | 0.624540 | 3.595786 | -0.077690 |
| H | -5.367807 | 0.037166 | 0.026967 | H | 3.089663 | 3.194336 | -0.018449 |
| H | -3.786675 | 1.960467 | -0.065868 | H | 3.968632 | 0.871511 | 0.027402 |
| H | -2.051261 | -2.735102 | 0.152446 | N | -0.207960 | -0.901439 | 0.047524 |
| H | -4.504268 | -2.298687 | 0.140068 | N | -1.186134 | 1.755522 | -0.045986 |
| C | -2.017925 | 0.708516 | -0.011540 | C | 2.645772 | -1.431828 | -0.080194 |
| C | -1.519557 | -0.637049 | 0.046404 | O | 2.159816 | -2.383511 | -0.644587 |
| C | 0.125842 | 1.489812 | -0.041310 | O | 3.839543 | -1.490059 | 0.533282 |
| C | 1.042480 | 2.588707 | -0.050934 | H | 4.189200 | -2.383375 | 0.381730 |
| C | 2.386282 | 2.361274 | -0.017863 | | | | |

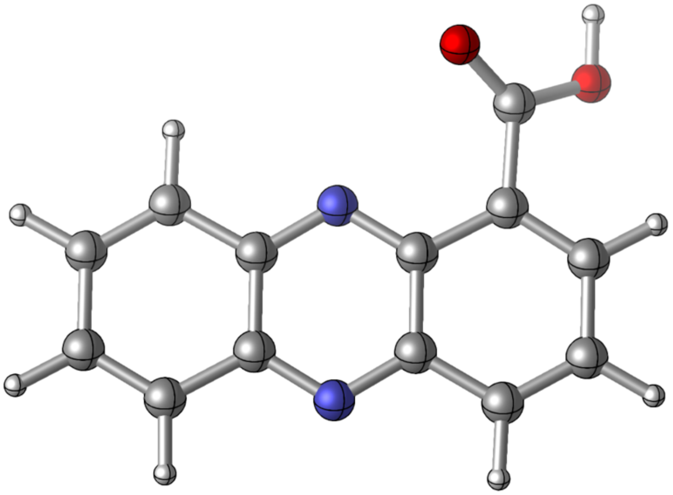

Zero-point correction= 0.186892 (Hartree/Particle)
Thermal correction to Energy= 0.198931
Thermal correction to Enthalpy= 0.199875
Thermal correction to Gibbs Free Energy= 0.147772
Sum of electronic and zero-point Energies= -759.793352
Sum of electronic and thermal Energies= -759.781313
Sum of electronic and thermal Enthalpies= -759.780369
Sum of electronic and thermal Free Energies= -759.832472

**PCA_oxidized_closed**
E(scf) = -759.992882352 a.u.

| | | | | | | | |
|---|---|---|---|---|---|---|---|
| C | -4.246355 | -0.041451 | 0.000047 | C | 2.953185 | 0.977558 | -0.000027 |
| C | -3.368867 | 1.003913 | 0.000027 | C | 2.096482 | -0.092601 | -0.000021 |
| C | -2.439176 | -1.672719 | 0.000055 | C | 0.677969 | 0.155792 | 0.000000 |
| C | -3.775870 | -1.392602 | 0.000062 | H | 0.749435 | 3.602620 | -0.000004 |
| H | -5.320797 | 0.146200 | 0.000052 | H | 3.203105 | 3.137897 | -0.000018 |
| H | -3.702591 | 2.042056 | 0.000016 | H | 4.025052 | 0.778381 | -0.000043 |
| H | -2.063739 | -2.696645 | 0.000064 | N | -0.181579 | -0.870138 | 0.000022 |
| H | -4.502095 | -2.206630 | 0.000079 | N | -1.109233 | 1.788058 | 0.000005 |
| C | -1.959629 | 0.755414 | 0.000020 | C | 2.693398 | -1.477027 | -0.000061 |
| C | -1.492803 | -0.603224 | 0.000033 | O | 3.893851 | -1.652525 | -0.000135 |
| C | 0.199130 | 1.508722 | -0.000001 | O | 1.852795 | -2.503169 | 0.000002 |
| C | 1.141292 | 2.584980 | -0.000007 | H | 0.918551 | -2.153510 | 0.000049 |
| C | 2.480509 | 2.321535 | -0.000015 | | | | |

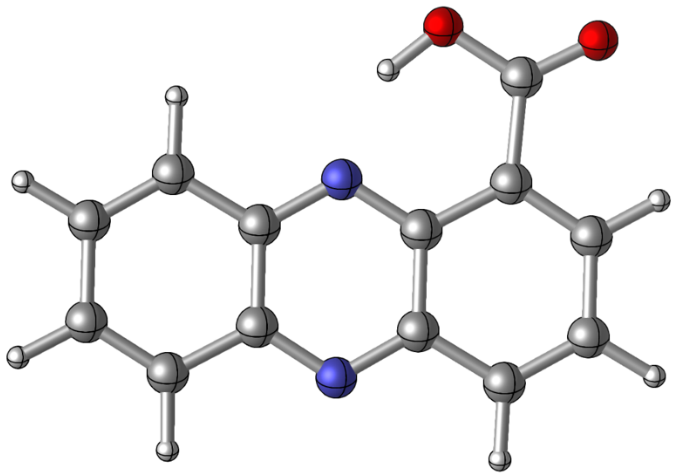

Zero-point correction= 0.187337 (Hartree/Particle)
Thermal correction to Energy= 0.198982
Thermal correction to Enthalpy= 0.199926
Thermal correction to Gibbs Free Energy= 0.149183
Sum of electronic and zero-point Energies= -759.805546
Sum of electronic and thermal Energies= -759.793900
Sum of electronic and thermal Enthalpies= -759.792956
Sum of electronic and thermal Free Energies= -759.843699

**PCA_reduced_open_octanol**
E(scf) = -761.199691694 a.u.

| | | | | | | | |
|---|---|---|---|---|---|---|---|
| C | -4.261289 | -0.087140 | 0.251274 | C | 3.013527 | 0.981461 | 0.189518 |
| C | -3.357376 | 0.974508 | 0.125126 | C | 2.109865 | -0.086167 | 0.015226 |
| C | -2.456185 | -1.656489 | -0.079256 | C | 0.745131 | 0.172405 | -0.176806 |
| C | -3.813766 | -1.398443 | 0.146331 | H | 0.808192 | 3.566227 | 0.183726 |
| H | -5.315783 | 0.124526 | 0.433564 | H | 3.247466 | 3.107219 | 0.415032 |
| H | -3.702675 | 2.007456 | 0.206175 | H | 4.073619 | 0.750328 | 0.291481 |
| H | -2.092720 | -2.683764 | -0.152714 | N | -0.196446 | -0.807032 | -0.487416 |
| H | -4.510897 | -2.231637 | 0.243708 | N | -1.076782 | 1.746509 | -0.327862 |
| C | -2.012194 | 0.724266 | -0.126082 | C | 2.689993 | -1.461532 | 0.020020 |
| C | -1.559450 | -0.604228 | -0.223327 | O | 3.780680 | -1.733342 | -0.417137 |
| C | 0.283862 | 1.509455 | -0.107513 | O | 1.939803 | -2.445492 | 0.571470 |
| C | 1.181434 | 2.541471 | 0.127701 | H | 1.224197 | -2.061727 | 1.099993 |
| C | 2.553265 | 2.281455 | 0.257647 | | | | |

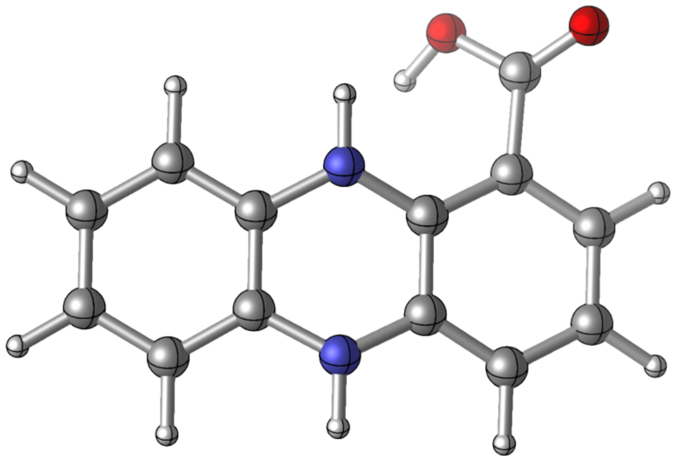

Zero-point correction= 0.209795 (Hartree/Particle)
Thermal correction to Energy= 0.222919
Thermal correction to Enthalpy= 0.223863
Thermal correction to Gibbs Free Energy= 0.169449
Sum of electronic and zero-point Energies= -760.989896
Sum of electronic and thermal Energies= -760.976773
Sum of electronic and thermal Enthalpies= -760.975828
Sum of electronic and thermal Free Energies= -761.030243

**PCA_reduced_closed_octanol**
E(scf) = -761.214907541 a.u.

| | | | | | | | |
|---|---|---|---|---|---|---|---|
| C | 4.299660 | -0.126425 | -0.254270 | C | -2.977831 | 1.017053 | -0.172076 |
| C | 3.407050 | 0.939837 | -0.089576 | C | -2.086642 | -0.070063 | -0.007772 |
| C | 2.466329 | -1.683501 | -0.043958 | C | -0.708979 | 0.169353 | 0.153258 |
| C | 3.831505 | -1.434433 | -0.230742 | H | -0.744205 | 3.576476 | -0.081127 |
| H | 5.360588 | 0.079503 | -0.403062 | H | -3.188194 | 3.148308 | -0.328328 |
| H | 3.767215 | 1.970791 | -0.109978 | H | -4.041186 | 0.812023 | -0.281006 |
| H | 2.086511 | -2.707277 | -0.031342 | N | 0.212091 | -0.838054 | 0.355429 |
| H | 4.518726 | -2.271607 | -0.360112 | N | 1.128948 | 1.722568 | 0.341134 |
| C | 2.052901 | 0.696525 | 0.115969 | C | -2.596086 | -1.453682 | -0.007716 |
| C | 1.578289 | -0.628509 | 0.135651 | O | -1.916580 | -2.461616 | 0.128687 |
| C | -0.236539 | 1.508512 | 0.128449 | O | -3.921425 | -1.537092 | -0.180568 |
| C | -1.125355 | 2.553150 | -0.058305 | H | -4.143879 | -2.481522 | -0.165145 |
| C | -2.503573 | 2.309861 | -0.198385 | | | | |

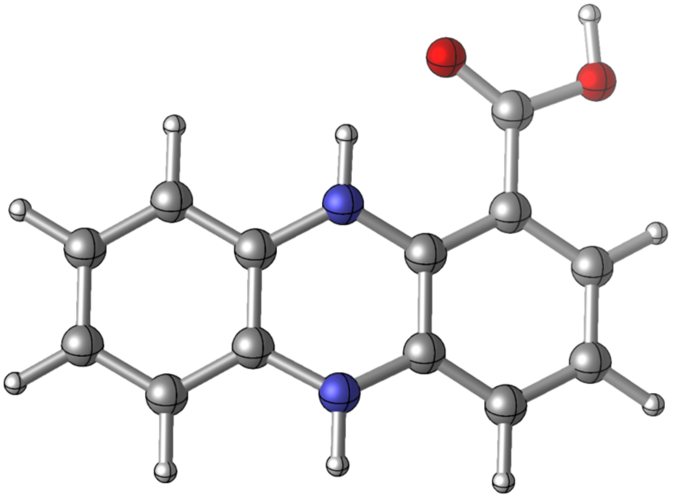

Zero-point correction= 0.210677 (Hartree/Particle)
Thermal correction to Energy= 0.223327
Thermal correction to Enthalpy= 0.224271
Thermal correction to Gibbs Free Energy= 0.171287
Sum of electronic and zero-point Energies= -761.004230
Sum of electronic and thermal Energies= -760.991580
Sum of electronic and thermal Enthalpies= -760.990636
Sum of electronic and thermal Free Energies= -761.043621

**PCA_oxidized_open_octanol**
E(scf) = -759.979014688 a.u.

| | | | | | | | |
|---|---|---|---|---|---|---|---|
| C | -4.289349 | -0.131025 | 0.032079 | C | 2.890165 | 1.028462 | 0.011327 |
| C | -3.431899 | 0.929830 | -0.019962 | C | 2.055153 | -0.060132 | -0.005938 |
| C | -2.451119 | -1.721641 | 0.105826 | C | 0.625483 | 0.141233 | -0.013077 |
| C | -3.793225 | -1.471912 | 0.097211 | H | 0.623646 | 3.595662 | -0.077573 |
| H | -5.367408 | 0.036631 | 0.025858 | H | 3.089245 | 3.194447 | -0.017934 |
| H | -3.786246 | 1.960365 | -0.066861 | H | 3.968262 | 0.871086 | 0.027926 |
| H | -2.049545 | -2.734635 | 0.152682 | N | -0.207844 | -0.901247 | 0.048414 |
| H | -4.503406 | -2.299141 | 0.139537 | N | -1.186366 | 1.755883 | -0.046069 |
| C | -2.017753 | 0.708672 | -0.011717 | C | 2.644806 | -1.432050 | -0.080187 |
| C | -1.519207 | -0.636785 | 0.046715 | O | 2.157326 | -2.384676 | -0.640315 |
| C | 0.125518 | 1.489969 | -0.041126 | O | 3.842326 | -1.489228 | 0.528434 |
| C | 1.042137 | 2.588818 | -0.050699 | H | 4.188007 | -2.383925 | 0.377014 |
| C | 2.385905 | 2.361275 | -0.017454 | | | | |

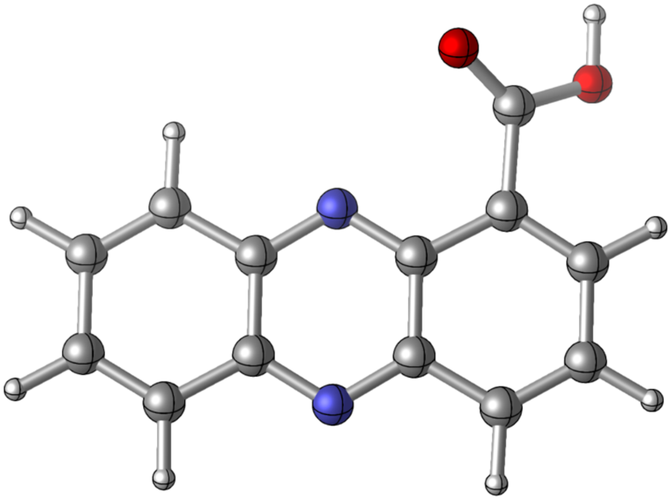

Zero-point correction= 0.186912 (Hartree/Particle)
Thermal correction to Energy= 0.198950
Thermal correction to Enthalpy= 0.199894
Thermal correction to Gibbs Free Energy= 0.147789
Sum of electronic and zero-point Energies= -759.792102
Sum of electronic and thermal Energies= -759.780065
Sum of electronic and thermal Enthalpies= -759.779120
Sum of electronic and thermal Free Energies= -759.831226

**PCA_oxidized_closed_octanol**
E(scf) = -759.991601693 a.u.

| | | | | | | | |
|---|---|---|---|---|---|---|---|
| C | -4.247046 | -0.042524 | 0.000073 | C | 2.952011 | 0.978030 | -0.000105 |
| C | -3.369700 | 1.002951 | 0.000093 | C | 2.095996 | -0.092548 | -0.000047 |
| C | -2.439530 | -1.673216 | -0.000045 | C | 0.677332 | 0.155435 | -0.000025 |
| C | -3.776283 | -1.393520 | 0.000002 | H | 0.746783 | 3.602397 | -0.000030 |
| H | -5.321555 | 0.145029 | 0.000104 | H | 3.201170 | 3.138567 | -0.000170 |
| H | -3.703084 | 2.041213 | 0.000135 | H | 4.023779 | 0.777977 | -0.000120 |
| H | -2.063568 | -2.696979 | -0.000112 | N | -0.182012 | -0.870638 | -0.000053 |
| H | -4.502233 | -2.207855 | -0.000009 | N | -1.110386 | 1.787701 | 0.000034 |
| C | -1.960425 | 0.754738 | 0.000041 | C | 2.695976 | -1.475705 | 0.000055 |
| C | -1.493239 | -0.603555 | -0.000019 | O | 3.895975 | -1.647784 | 0.000171 |
| C | 0.197864 | 1.508465 | -0.000006 | O | 1.858179 | -2.504207 | -0.000003 |
| C | 1.139605 | 2.585118 | -0.000050 | H | 0.924140 | -2.157386 | -0.000080 |
| C | 2.478791 | 2.321916 | -0.000122 | | | | |

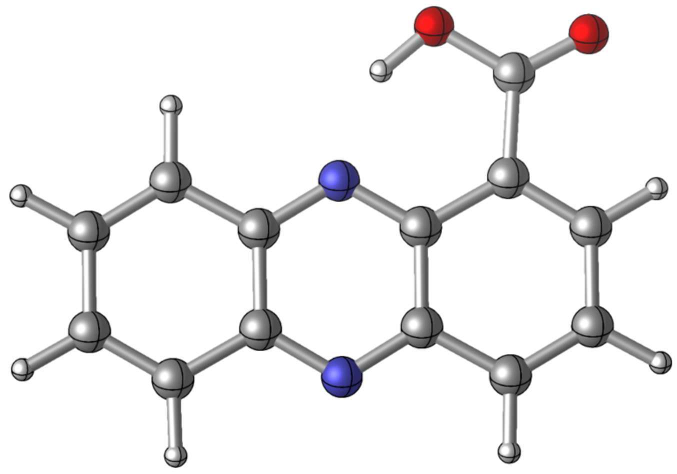

Zero-point correction= 0.187403 (Hartree/Particle)
Thermal correction to Energy= 0.199054
Thermal correction to Enthalpy= 0.199998
Thermal correction to Gibbs Free Energy= 0.149223
Sum of electronic and zero-point Energies= -759.804199
Sum of electronic and thermal Energies= -759.792548
Sum of electronic and thermal Enthalpies= -759.791604
Sum of electronic and thermal Free Energies= -759.842378

**PCA_reduced_negative_closed**
E(scf) = -760.737927513 a.u.

| | | | | | | | |
|---|---|---|---|---|---|---|---|
| C | 4.228089 | -0.080592 | -0.331218 | C | -2.994831 | 0.941148 | -0.238223 |
| C | 3.328840 | 0.974091 | -0.120281 | C | -2.107443 | -0.127220 | -0.016046 |
| C | 2.420506 | -1.658916 | -0.045872 | C | -0.752626 | 0.163867 | 0.206591 |
| C | 3.773949 | -1.393116 | -0.292929 | H | -0.826322 | 3.564790 | -0.105151 |
| H | 5.278901 | 0.138334 | -0.527533 | H | -3.251654 | 3.072727 | -0.448414 |
| H | 3.675549 | 2.009876 | -0.153210 | H | -4.044491 | 0.691034 | -0.392725 |
| H | 2.053636 | -2.687509 | -0.022382 | N | 0.180439 | -0.836966 | 0.483792 |
| H | 4.463810 | -2.222445 | -0.458446 | N | 1.065035 | 1.731446 | 0.437145 |
| C | 1.989619 | 0.715024 | 0.150943 | C | -2.649268 | -1.561069 | -0.033145 |
| C | 1.523711 | -0.617938 | 0.185853 | O | -1.809279 | -2.502678 | 0.099870 |
| C | -0.298591 | 1.501650 | 0.173327 | O | -3.880254 | -1.698651 | -0.186800 |
| C | -1.191459 | 2.535166 | -0.075127 | H | -0.248397 | -1.766557 | 0.344832 |
| C | -2.552249 | 2.253938 | -0.269527 | H | 1.387433 | 2.672824 | 0.245840 |

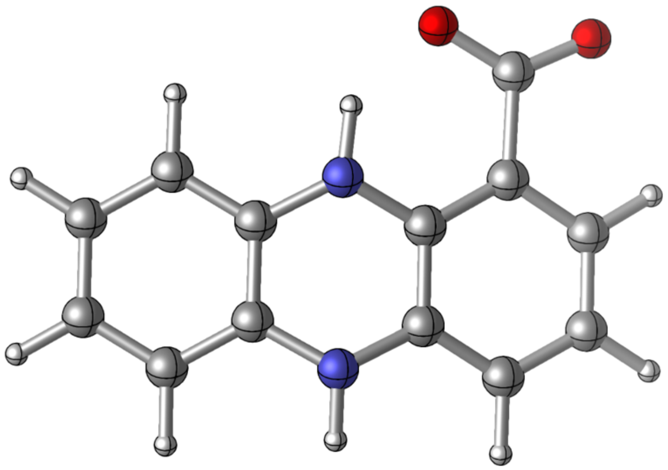

Zero-point correction= 0.197392 (Hartree/Particle)
Thermal correction to Energy= 0.209658
Thermal correction to Enthalpy= 0.210603
Thermal correction to Gibbs Free Energy= 0.158373
Sum of electronic and zero-point Energies= -760.540536
Sum of electronic and thermal Energies= -760.528269
Sum of electronic and thermal Enthalpies= -760.527325
Sum of electronic and thermal Free Energies= -760.579554

**PCA_oxidized_negative**
E(scf) = -759.498515862 a.u.

| | | | | | | | |
|---|---|---|---|---|---|---|---|
| C | -4.246396 | -0.096328 | 0.027940 | C | 2.457872 | 2.311626 | -0.008023 |
| C | -3.375913 | 0.954335 | -0.033302 | C | 2.941461 | 0.969543 | 0.045500 |
| C | -2.424514 | -1.701425 | 0.127911 | C | 2.103719 | -0.115551 | 0.013985 |
| C | -3.765041 | -1.440946 | 0.111767 | C | 0.680039 | 0.125624 | -0.004984 |
| H | -5.322587 | 0.083645 | 0.014849 | H | 0.719007 | 3.583491 | -0.101789 |
| H | -3.720600 | 1.987766 | -0.094157 | H | 3.174383 | 3.135556 | -0.007058 |
| H | -2.033432 | -2.718202 | 0.188676 | H | 4.017685 | 0.795292 | 0.101943 |
| H | -4.483192 | -2.260970 | 0.161973 | N | -0.170309 | -0.906253 | 0.064589 |
| C | -1.963614 | 0.720720 | -0.016302 | N | -1.121647 | 1.760867 | -0.058788 |
| C | -1.479599 | -0.627519 | 0.058618 | C | 2.706251 | -1.530944 | -0.046480 |
| C | 0.188939 | 1.482782 | -0.045709 | O | 2.303707 | -2.253309 | -0.983421 |
| C | 1.118846 | 2.569418 | -0.060416 | O | 3.576308 | -1.786302 | 0.817412 |

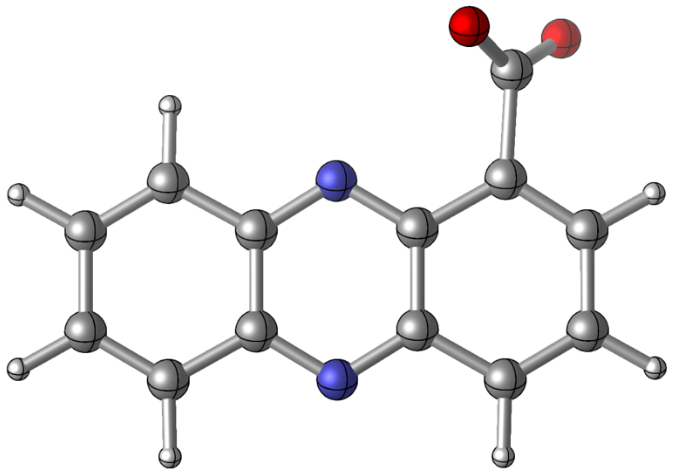

Zero-point correction= 0.173632 (Hartree/Particle)
Thermal correction to Energy= 0.185528
Thermal correction to Enthalpy= 0.186472
Thermal correction to Gibbs Free Energy= 0.134397
Sum of electronic and zero-point Energies= -759.324884
Sum of electronic and thermal Energies= -759.312988
Sum of electronic and thermal Enthalpies= -759.312044
Sum of electronic and thermal Free Energies= -759.364119

**PCA_reduced_negative_octanol**
E(scf) = -760.729175798 a.u.

| | | | | | | | |
|---|---|---|---|---|---|---|---|
| C | 4.228125 | -0.079914 | -0.329975 | C | -2.554966 | 2.252764 | -0.267270 |
| C | 3.327825 | 0.974602 | -0.121144 | C | -2.995915 | 0.939194 | -0.235921 |
| C | 2.420123 | -1.657814 | -0.044340 | C | -2.107379 | -0.127666 | -0.014871 |
| C | 3.773795 | -1.392073 | -0.290168 | C | -0.753112 | 0.165284 | 0.206512 |
| H | 5.279128 | 0.139023 | -0.525581 | H | -0.830168 | 3.565765 | -0.104134 |
| H | 3.674346 | 2.010582 | -0.154725 | H | -3.255697 | 3.070748 | -0.445227 |
| H | 2.052825 | -2.686170 | -0.020528 | H | -4.045012 | 0.685909 | -0.389617 |
| H | 4.463772 | -2.221759 | -0.453937 | N | 0.179882 | -0.836399 | 0.480121 |
| C | 1.988544 | 0.716104 | 0.149162 | N | 1.064552 | 1.733309 | 0.434600 |
| C | 1.521852 | -0.617306 | 0.185083 | C | -2.646592 | -1.564173 | -0.033284 |
| C | -0.300376 | 1.502805 | 0.172836 | O | -1.802336 | -2.501786 | 0.101190 |
| C | -1.194305 | 2.535608 | -0.074153 | O | -3.876152 | -1.702520 | -0.189234 |

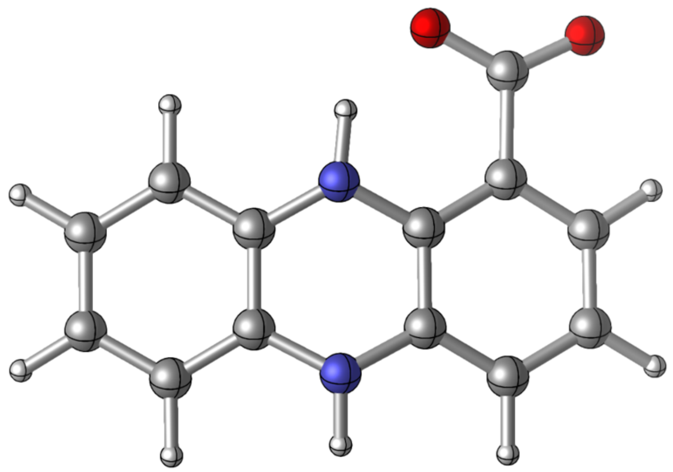

Zero-point correction= 0.197300 (Hartree/Particle)
Thermal correction to Energy= 0.209571
Thermal correction to Enthalpy= 0.210515
Thermal correction to Gibbs Free Energy= 0.158285
Sum of electronic and zero-point Energies= -760.531876
Sum of electronic and thermal Energies= -760.519605
Sum of electronic and thermal Enthalpies= -760.518661
Sum of electronic and thermal Free Energies= -760.570891

**PCA_oxidized_negative_octanol**
E(scf) = -759.488860169 a.u.

| | | | | | | | |
|---|---|---|---|---|---|---|---|
| C | -4.247441 | -0.097964 | 0.028371 | C | 2.455547 | 2.311039 | -0.008391 |
| C | -3.377257 | 0.953063 | -0.033924 | C | 2.939584 | 0.969387 | 0.047030 |
| C | -2.424698 | -1.701368 | 0.129655 | C | 2.103121 | -0.116653 | 0.015456 |
| C | -3.765526 | -1.442137 | 0.113259 | C | 0.679145 | 0.124857 | -0.004679 |
| H | -5.323804 | 0.081805 | 0.015193 | H | 0.716043 | 3.583088 | -0.104890 |
| H | -3.721876 | 1.986528 | -0.095683 | H | 3.171969 | 3.135293 | -0.007593 |
| H | -2.031307 | -2.717206 | 0.190861 | H | 4.015283 | 0.793337 | 0.106301 |
| H | -4.483015 | -2.262831 | 0.163947 | N | -0.171378 | -0.906166 | 0.066322 |
| C | -1.964858 | 0.720416 | -0.016872 | N | -1.123344 | 1.761128 | -0.060244 |
| C | -1.480439 | -0.626969 | 0.059495 | C | 2.710981 | -1.532125 | -0.046678 |
| C | 0.187063 | 1.482584 | -0.046890 | O | 2.292839 | -2.259368 | -0.971763 |
| C | 1.116632 | 2.569336 | -0.062262 | O | 3.598240 | -1.773825 | 0.802750 |

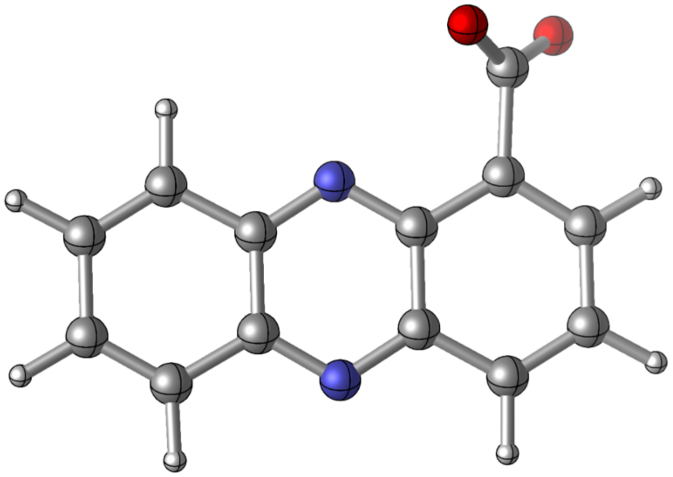

Zero-point correction= 0.173589 (Hartree/Particle)
Thermal correction to Energy= 0.185485
Thermal correction to Enthalpy= 0.186429
Thermal correction to Gibbs Free Energy= 0.134403
Sum of electronic and zero-point Energies= -759.315271
Sum of electronic and thermal Energies= -759.303376
Sum of electronic and thermal Enthalpies= -759.302431
Sum of electronic and thermal Free Energies= -759.354457

**Pyocyanin_reduced_closed**
E(scf) = -687.172803951 a.u.

| | | | | | | | |
|---|---|---|---|---|---|---|---|
| C | 3.769293 | 0.059908 | 0.475917 | C | -2.372023 | -1.064893 | 0.026112 |
| C | 2.716227 | 0.959781 | 0.260445 | C | -1.116365 | -0.511874 | -0.227138 |
| C | 2.282851 | -1.780057 | 0.010883 | H | -1.847965 | 2.764622 | 0.384444 |
| C | 3.555878 | -1.305627 | 0.349036 | H | -4.079224 | 1.774243 | 0.770505 |
| H | 4.752989 | 0.444405 | 0.749596 | H | -4.431181 | -0.683846 | 0.553032 |
| H | 2.895657 | 2.027514 | 0.382930 | N | -0.036495 | -1.321916 | -0.618450 |
| H | 2.097562 | -2.852656 | -0.080861 | N | 0.371841 | 1.355165 | -0.410096 |
| H | 4.368683 | -2.013680 | 0.517639 | O | -2.458876 | -2.418936 | -0.106403 |
| C | 1.454301 | 0.500371 | -0.114814 | C | 0.595776 | 2.782472 | -0.459862 |
| C | 1.240241 | -0.891871 | -0.229427 | H | 0.679480 | 3.248050 | 0.539960 |
| C | -0.917564 | 0.872805 | -0.113498 | H | 1.518685 | 2.985715 | -1.018515 |
| C | -1.986974 | 1.691275 | 0.265376 | H | -0.230804 | 3.260748 | -1.000674 |
| C | -3.245660 | 1.128153 | 0.491517 | H | -3.354180 | -2.707347 | 0.108697 |
| C | -3.448422 | -0.242876 | 0.373391 | H | -0.211473 | -2.314414 | -0.503344 |

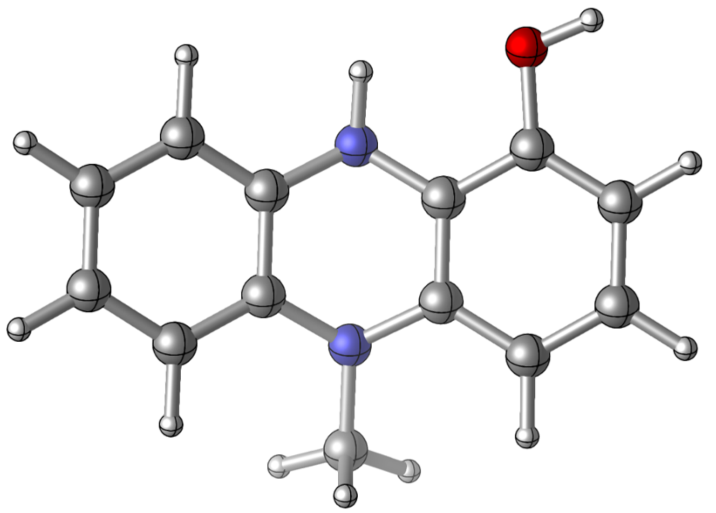

Zero-point correction= 0.228000 (Hartree/Particle)
Thermal correction to Energy= 0.240437
Thermal correction to Enthalpy= 0.241382
Thermal correction to Gibbs Free Energy= 0.189938
Sum of electronic and zero-point Energies= -686.944804
Sum of electronic and thermal Energies= -686.932367
Sum of electronic and thermal Enthalpies= -686.931422
Sum of electronic and thermal Free Energies= -686.982866

**Pyocyanin_reduced_closed_octanol**
E(scf) = -687.171721153 a.u.

| | | | | | | | |
|---|---|---|---|---|---|---|---|
| C | 3.769989 | 0.060489 | 0.474777 | C | -2.372054 | -1.064754 | 0.025963 |
| C | 2.716408 | 0.959956 | 0.260703 | C | -1.116258 | -0.512090 | -0.226103 |
| C | 2.283840 | -1.779467 | 0.010081 | H | -1.848770 | 2.764174 | 0.384585 |
| C | 3.556946 | -1.304864 | 0.347248 | H | -4.080510 | 1.774084 | 0.767895 |
| H | 4.753781 | 0.445118 | 0.747820 | H | -4.431934 | -0.683853 | 0.550474 |
| H | 2.895434 | 2.027687 | 0.383813 | N | -0.036296 | -1.322003 | -0.615839 |
| H | 2.098967 | -2.852099 | -0.082209 | N | 0.371550 | 1.355021 | -0.407706 |
| H | 4.370182 | -2.012686 | 0.514583 | O | -2.458907 | -2.419334 | -0.105932 |
| C | 1.454471 | 0.500396 | -0.113679 | C | 0.595191 | 2.781786 | -0.460615 |
| C | 1.240645 | -0.891734 | -0.228592 | H | 0.679120 | 3.249921 | 0.538149 |
| C | -0.917853 | 0.872604 | -0.112434 | H | 1.518003 | 2.983833 | -1.019987 |
| C | -1.987728 | 1.690870 | 0.265174 | H | -0.231625 | 3.258725 | -1.002387 |
| C | -3.246631 | 1.128008 | 0.489929 | H | -3.355069 | -2.706277 | 0.106454 |
| C | -3.448911 | -0.242964 | 0.371776 | H | -0.211429 | -2.314495 | -0.502288 |

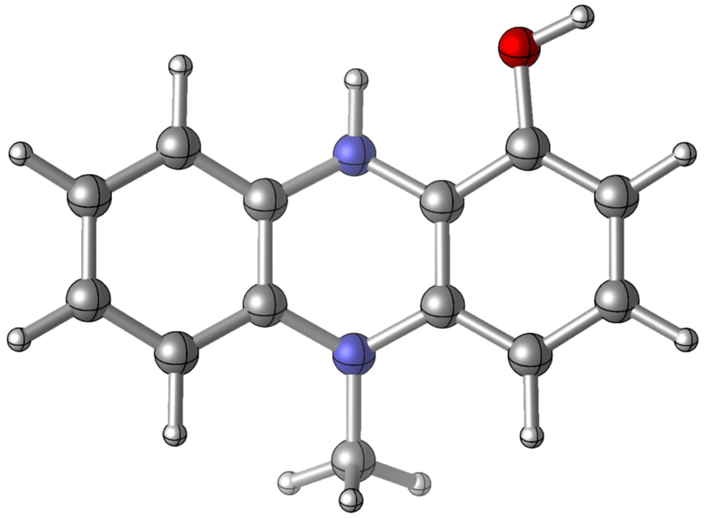

Zero-point correction= 0.228019 (Hartree/Particle)
Thermal correction to Energy= 0.240462
Thermal correction to Enthalpy= 0.241406
Thermal correction to Gibbs Free Energy= 0.189954
Sum of electronic and zero-point Energies= -686.943702
Sum of electronic and thermal Energies= -686.931260
Sum of electronic and thermal Enthalpies= -686.930315
Sum of electronic and thermal Free Energies= -686.981767

**Pyocyanin_oxidized**
E(scf) = -685.939462987 a.u.

| | | | | | | | |
|---|---|---|---|---|---|---|---|
| C | 3.816581 | 0.076282 | 0.085863 | C | -3.553944 | -0.347071 | 0.059604 |
| C | 2.767988 | 0.977394 | 0.056406 | C | -2.441816 | -1.262214 | -0.027344 |
| C | 2.302339 | -1.782974 | 0.001714 | C | -1.070620 | -0.619298 | -0.026090 |
| C | 3.596371 | -1.312816 | 0.051996 | H | -2.010356 | 2.708822 | 0.164917 |
| H | 4.836009 | 0.460586 | 0.144345 | H | -4.216751 | 1.662708 | 0.194897 |
| H | 2.989692 | 2.040612 | 0.109384 | H | -4.558815 | -0.771019 | 0.074068 |
| H | 2.075029 | -2.849528 | -0.015046 | N | -0.051314 | -1.421899 | -0.055431 |
| H | 4.439006 | -2.003783 | 0.074434 | N | 0.341938 | 1.347828 | -0.061319 |
| C | 1.438238 | 0.509814 | -0.012440 | O | -2.551125 | -2.486980 | -0.088933 |
| C | 1.210753 | -0.889384 | -0.023554 | C | 0.494138 | 2.793403 | -0.167434 |
| C | -0.938409 | 0.839731 | 0.012720 | H | 0.329261 | 3.276051 | 0.806861 |
| C | -2.066474 | 1.625696 | 0.099520 | H | 1.488875 | 3.045335 | -0.538599 |
| C | -3.348049 | 1.003158 | 0.122644 | H | -0.239888 | 3.174233 | -0.888173 |

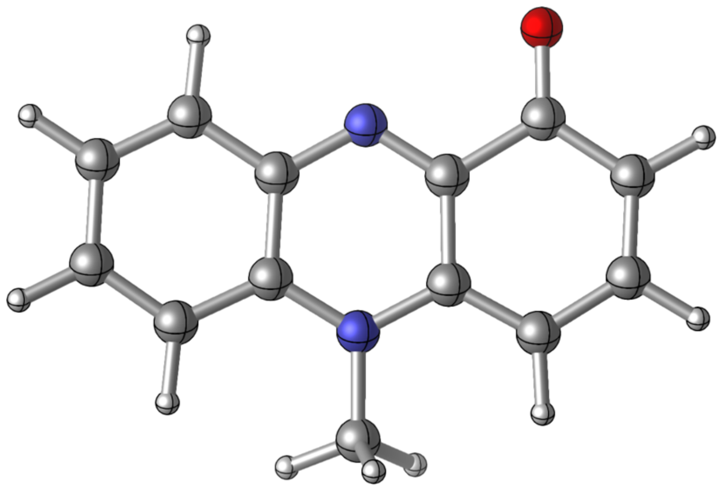

Zero-point correction= 0.204471 (Hartree/Particle)
Thermal correction to Energy= 0.216363
Thermal correction to Enthalpy= 0.217307
Thermal correction to Gibbs Free Energy= 0.166121
Sum of electronic and zero-point Energies= -685.734992
Sum of electronic and thermal Energies= -685.723100
Sum of electronic and thermal Enthalpies= -685.722156
Sum of electronic and thermal Free Energies= -685.773342

**Pyocyanin_oxidized_octanol**
E(scf) = -685.937499895 a.u.

| | | | | | | | |
|---|---|---|---|---|---|---|---|
| C | 3.817491 | 0.074970 | 0.088843 | C | -3.556081 | -0.345610 | 0.062926 |
| C | 2.768537 | 0.976521 | 0.057548 | C | -2.442826 | -1.261943 | -0.029331 |
| C | 2.301710 | -1.782749 | 0.003427 | C | -1.070148 | -0.619451 | -0.027345 |
| C | 3.596644 | -1.313236 | 0.055302 | H | -2.009334 | 2.707978 | 0.172718 |
| H | 4.836978 | 0.459084 | 0.148760 | H | -4.217211 | 1.663211 | 0.204939 |
| H | 2.990010 | 2.039899 | 0.111023 | H | -4.560350 | -0.770686 | 0.077997 |
| H | 2.073224 | -2.849041 | -0.013171 | N | -0.052087 | -1.422053 | -0.057329 |
| H | 4.438610 | -2.005007 | 0.079285 | N | 0.342794 | 1.348216 | -0.064378 |
| C | 1.439242 | 0.509402 | -0.013168 | O | -2.553055 | -2.484462 | -0.095274 |
| C | 1.211213 | -0.889254 | -0.023846 | C | 0.496274 | 2.792371 | -0.175068 |
| C | -0.938392 | 0.840242 | 0.013582 | H | 0.343459 | 3.279379 | 0.799349 |
| C | -2.065535 | 1.624893 | 0.104689 | H | 1.487518 | 3.041161 | -0.558500 |
| C | -3.349514 | 1.002723 | 0.129136 | H | -0.245109 | 3.173310 | -0.888442 |

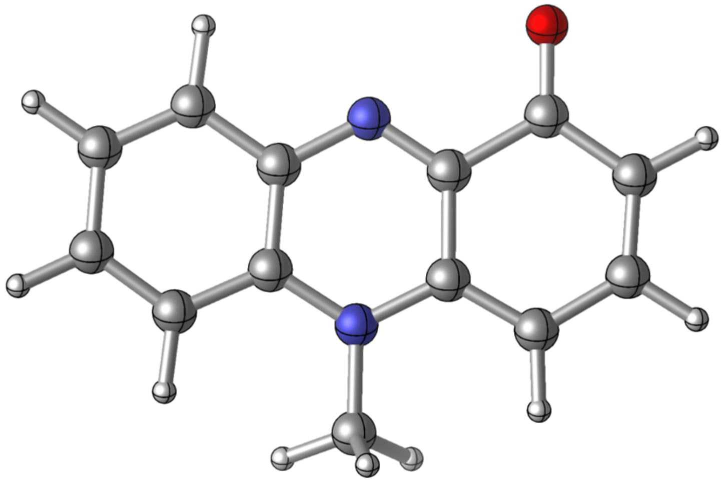

Zero-point correction= 0.204450 (Hartree/Particle)
Thermal correction to Energy= 0.216357
Thermal correction to Enthalpy= 0.217301
Thermal correction to Gibbs Free Energy= 0.166097
Sum of electronic and zero-point Energies= -685.733050
Sum of electronic and thermal Energies= -685.721143
Sum of electronic and thermal Enthalpies= -685.720199
Sum of electronic and thermal Free Energies= -685.771403

**Pyocyanin_oxidized_protonated_closed**
E(scf) = -686.399216694 a.u.

| | | | | | | | |
|---|---|---|---|---|---|---|---|
| C | 3.795705 | 0.081443 | 0.078293 | C | -3.492922 | -0.334325 | 0.039920 |
| C | 2.765322 | 0.988548 | 0.058985 | C | -2.388366 | -1.148490 | -0.008160 |
| C | 2.290994 | -1.796676 | -0.017522 | C | -1.070370 | -0.553340 | -0.018980 |
| C | 3.569320 | -1.320492 | 0.030173 | H | -2.016607 | 2.763922 | 0.121293 |
| H | 4.819069 | 0.453020 | 0.138981 | H | -4.200045 | 1.695550 | 0.140181 |
| H | 2.991872 | 2.049216 | 0.121360 | H | -4.490839 | -0.769689 | 0.053983 |
| H | 2.062616 | -2.861875 | -0.041950 | N | -0.048953 | -1.389964 | -0.047784 |
| H | 4.417098 | -2.004659 | 0.041735 | N | 0.350334 | 1.364088 | -0.046081 |
| C | 1.428158 | 0.523352 | -0.007435 | O | -2.483903 | -2.473173 | -0.037001 |
| C | 1.190980 | -0.890441 | -0.025981 | C | 0.549850 | 2.818425 | -0.124026 |
| C | -0.923403 | 0.875137 | -0.000319 | H | 0.558954 | 3.246773 | 0.886221 |
| C | -2.076009 | 1.681369 | 0.063076 | H | 1.488988 | 3.031486 | -0.636583 |
| C | -3.311883 | 1.064777 | 0.083234 | H | -0.257799 | 3.256015 | -0.715100 |

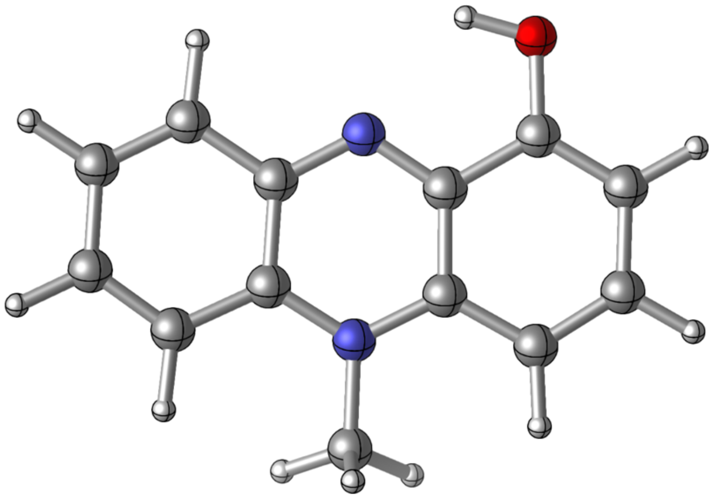

Zero-point correction= 0.218585 (Hartree/Particle)
Thermal correction to Energy= 0.230417
Thermal correction to Enthalpy= 0.231361
Thermal correction to Gibbs Free Energy= 0.180725
Sum of electronic and zero-point Energies= -686.180632
Sum of electronic and thermal Energies= -686.168800
Sum of electronic and thermal Enthalpies= -686.167855
Sum of electronic and thermal Free Energies= -686.218491

**Pyocyanin_oxidized_protonated_open**
E(scf) = -686.393711483 a.u.

| | | | | | | | |
|---|---|---|---|---|---|---|---|
| C | 3.812853 | 0.047861 | 0.081289 | C | -3.483875 | -0.279570 | 0.043777 |
| C | 2.790024 | 0.963508 | 0.060682 | C | -2.399999 | -1.122517 | -0.009372 |
| C | 2.292140 | -1.815848 | -0.017345 | C | -1.063937 | -0.563881 | -0.021262 |
| C | 3.574776 | -1.352090 | 0.032304 | H | -1.949260 | 2.781303 | 0.125715 |
| H | 4.839206 | 0.411031 | 0.143814 | H | -4.158880 | 1.762179 | 0.150622 |
| H | 3.026432 | 2.021947 | 0.124416 | H | -4.495863 | -0.685159 | 0.060281 |
| H | 2.052835 | -2.878639 | -0.042488 | N | -0.041277 | -1.402675 | -0.050948 |
| H | 4.416599 | -2.043656 | 0.044828 | N | 0.375115 | 1.351916 | -0.047935 |
| C | 1.449179 | 0.508352 | -0.007652 | O | -2.466789 | -2.452523 | -0.044116 |
| C | 1.197223 | -0.901349 | -0.026731 | C | 0.578990 | 2.805473 | -0.130888 |
| C | -0.900440 | 0.866881 | -0.001226 | H | 0.548724 | 3.243887 | 0.874596 |
| C | -2.035183 | 1.700795 | 0.066074 | H | 1.537803 | 3.013702 | -0.606739 |
| C | -3.282321 | 1.116096 | 0.089539 | H | -0.203478 | 3.238454 | -0.758547 |

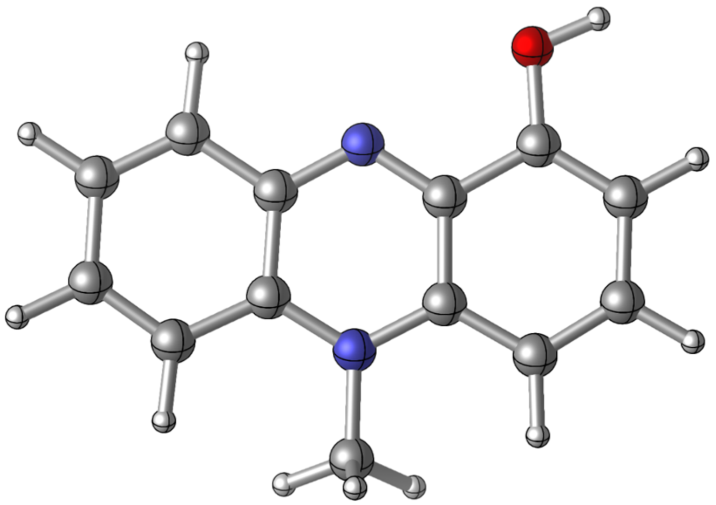

Zero-point correction= 0.218397 (Hartree/Particle)
Thermal correction to Energy= 0.230302
Thermal correction to Enthalpy= 0.231246
Thermal correction to Gibbs Free Energy= 0.180592
Sum of electronic and zero-point Energies= -686.175314
Sum of electronic and thermal Energies= -686.163410
Sum of electronic and thermal Enthalpies= -686.162466
Sum of electronic and thermal Free Energies= -686.213120

**Pyocyanin_oxidized_protonated_closed_octanol**
E(scf) = -686.393003486 a.u.

| | | | | | | | |
|---|---|---|---|---|---|---|---|
| C | 3.796316 | 0.080601 | 0.078611 | C | -3.493461 | -0.334190 | 0.040075 |
| C | 2.765948 | 0.988144 | 0.059396 | C | -2.388705 | -1.148734 | -0.008282 |
| C | 2.291025 | -1.796733 | -0.017612 | C | -1.070486 | -0.553293 | -0.019040 |
| C | 3.569676 | -1.321055 | 0.030206 | H | -2.017213 | 2.764042 | 0.122065 |
| H | 4.819823 | 0.451918 | 0.139487 | H | -4.200381 | 1.695094 | 0.140829 |
| H | 2.993141 | 2.048733 | 0.122149 | H | -4.491202 | -0.769906 | 0.054157 |
| H | 2.062380 | -2.861899 | -0.042198 | N | -0.048983 | -1.389478 | -0.047997 |
| H | 4.417205 | -2.005551 | 0.041695 | N | 0.350649 | 1.364405 | -0.046076 |
| C | 1.428819 | 0.523436 | -0.007249 | O | -2.485212 | -2.472410 | -0.037305 |
| C | 1.191357 | -0.890229 | -0.026022 | C | 0.550178 | 2.818512 | -0.124978 |
| C | -0.923660 | 0.875696 | -0.000081 | H | 0.559228 | 3.248464 | 0.884760 |
| C | -2.075885 | 1.681405 | 0.063496 | H | 1.489476 | 3.031438 | -0.637605 |
| C | -3.312211 | 1.064256 | 0.083649 | H | -0.257541 | 3.255919 | -0.716359 |

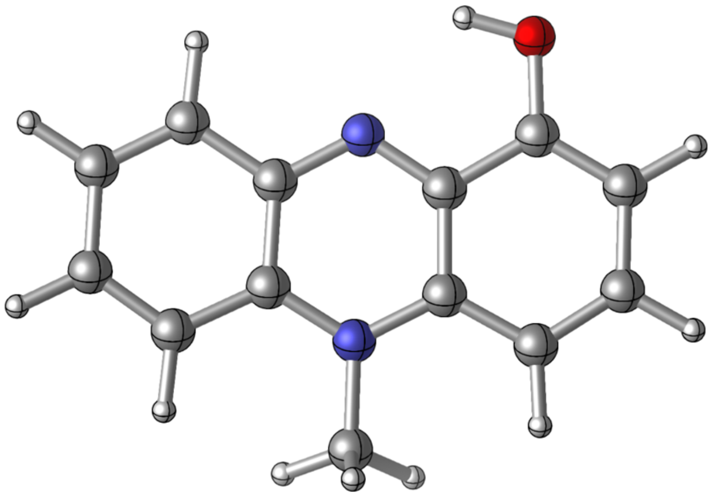

Zero-point correction= 0.218567 (Hartree/Particle)
Thermal correction to Energy= 0.230412
Thermal correction to Enthalpy= 0.231356
Thermal correction to Gibbs Free Energy= 0.180667
Sum of electronic and zero-point Energies= -686.174437
Sum of electronic and thermal Energies= -686.162592
Sum of electronic and thermal Enthalpies= -686.161648
Sum of electronic and thermal Free Energies= -686.212337

**Pyocyanin_oxidized_protonated_open_octanol**
E(scf) = -686.387409115 a.u.

| | | | | | | | |
|---|---|---|---|---|---|---|---|
| C | 3.813018 | 0.047650 | 0.081493 | C | -3.484166 | -0.280434 | 0.043878 |
| C | 2.790221 | 0.963764 | 0.060966 | C | -2.399753 | -1.123247 | -0.009445 |
| C | 2.292109 | -1.815624 | -0.017418 | C | -1.063843 | -0.563717 | -0.021281 |
| C | 3.574917 | -1.352087 | 0.032318 | H | -1.950855 | 2.781020 | 0.126351 |
| H | 4.839480 | 0.410651 | 0.144132 | H | -4.159856 | 1.760755 | 0.151068 |
| H | 3.027253 | 2.022114 | 0.124990 | H | -4.496170 | -0.686111 | 0.060362 |
| H | 2.052163 | -2.878270 | -0.042682 | N | -0.041010 | -1.402035 | -0.051080 |
| H | 4.416647 | -2.043763 | 0.044796 | N | 0.374890 | 1.352451 | -0.047880 |
| C | 1.449530 | 0.508940 | -0.007474 | O | -2.465147 | -2.452554 | -0.044389 |
| C | 1.197510 | -0.900836 | -0.026755 | C | 0.578046 | 2.805883 | -0.131587 |
| C | -0.900946 | 0.867143 | -0.001044 | H | 0.545271 | 3.246013 | 0.873217 |
| C | -2.035849 | 1.700394 | 0.066392 | H | 1.537864 | 3.014591 | -0.605437 |
| C | -3.283050 | 1.114931 | 0.089827 | H | -0.203411 | 3.238012 | -0.761370 |

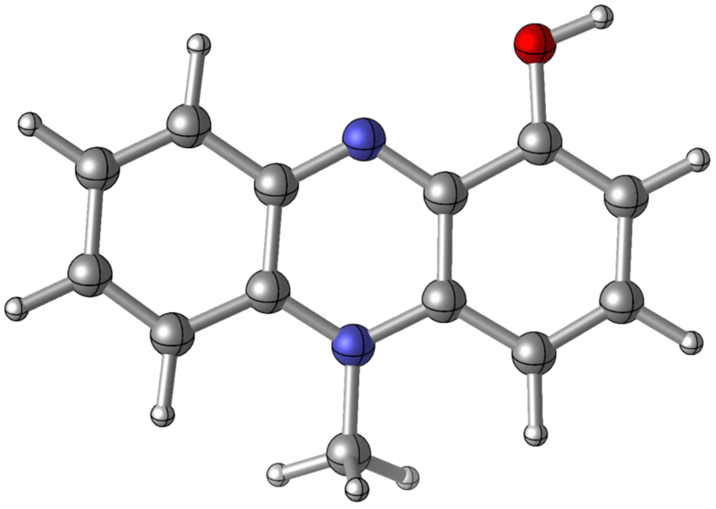

Zero-point correction= 0.218367 (Hartree/Particle)
Thermal correction to Energy= 0.230286
Thermal correction to Enthalpy= 0.231230
Thermal correction to Gibbs Free Energy= 0.180519
Sum of electronic and zero-point Energies= -686.169042
Sum of electronic and thermal Energies= -686.157123
Sum of electronic and thermal Enthalpies= -686.156179
Sum of electronic and thermal Free Energies= -686.206890

**OH-phenazine_reduced_closed**
E(scf) = -647.878437189 a.u.

| | | | |
|---|---|---|---|
| C | -3.835848 | -0.473704 | -0.323529 |
| C | -2.732907 | -1.304385 | -0.090556 |
| C | -2.415767 | 1.462222 | -0.064706 |
| C | -3.677183 | 0.906866 | -0.311290 |
| H | -4.812860 | -0.918262 | -0.518679 |
| H | -2.847157 | -2.390734 | -0.107917 |
| H | -2.280429 | 2.546183 | -0.060923 |
| H | -4.527849 | 1.564368 | -0.496849 |
| C | -1.481475 | -0.758493 | 0.177972 |
| C | -1.320486 | 0.642795 | 0.191924 |
| C | 0.914744 | -1.042687 | 0.184973 |
| C | 2.014320 | -1.860927 | -0.080611 |
| C | 3.264118 | -1.283799 | -0.312231 |
| C | 3.422001 | 0.099789 | -0.300612 |
| C | 2.314014 | 0.916031 | -0.051649 |
| C | 1.063024 | 0.350245 | 0.202036 |
| H | 1.888303 | -2.944977 | -0.098020 |
| H | 4.125889 | -1.923276 | -0.508342 |
| H | 4.397173 | 0.555021 | -0.485711 |
| N | -0.051261 | 1.155408 | 0.503810 |
| N | -0.360366 | -1.555355 | 0.468914 |
| O | 2.362705 | 2.277764 | -0.026095 |
| H | 0.084690 | 2.138293 | 0.295666 |
| H | -0.476967 | -2.545127 | 0.285199 |
| H | 3.257621 | 2.572303 | -0.235054 |

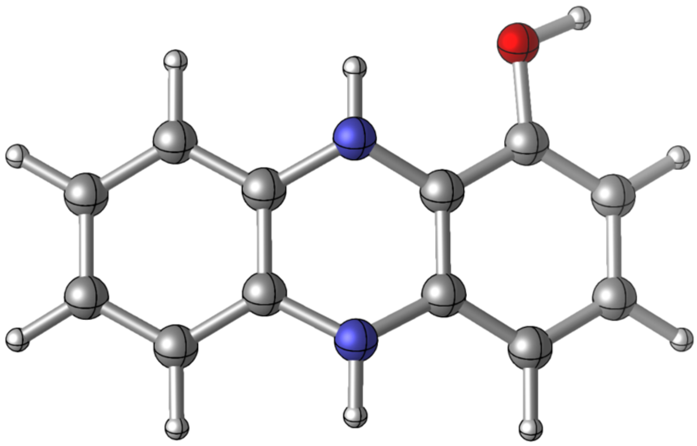

Zero-point correction= 0.199668 (Hartree/Particle)
Thermal correction to Energy= 0.210780
Thermal correction to Enthalpy= 0.211724
Thermal correction to Gibbs Free Energy= 0.162997
Sum of electronic and zero-point Energies= -647.678770
Sum of electronic and thermal Energies= -647.667658
Sum of electronic and thermal Enthalpies= -647.666713
Sum of electronic and thermal Free Energies= -647.715440

**OH-phenazine_reduced_closed_octanol**
E(scf) = -647.877216847 a.u.

| | | | | | | | |
|---|---|---|---|---|---|---|---|
| C | -3.836716 | -0.473875 | -0.321460 | C | 3.422701 | 0.099901 | -0.298279 |
| C | -2.733325 | -1.304293 | -0.090188 | C | 2.314201 | 0.915838 | -0.051344 |
| C | -2.416575 | 1.461826 | -0.063908 | C | 1.063013 | 0.350361 | 0.200741 |
| C | -3.678193 | 0.906511 | -0.308939 | H | 1.889517 | -2.944611 | -0.097174 |
| H | -4.813936 | -0.918471 | -0.515293 | H | 4.127390 | -1.923055 | -0.504194 |
| H | -2.847661 | -2.390665 | -0.107465 | H | 4.398249 | 0.555060 | -0.481854 |
| H | -2.281512 | 2.545824 | -0.059748 | N | -0.051364 | 1.155344 | 0.500348 |
| H | -4.529202 | 1.563947 | -0.492957 | N | -0.360147 | -1.555382 | 0.465588 |
| C | -1.481715 | -0.758487 | 0.176647 | O | 2.362807 | 2.278025 | -0.026756 |
| C | -1.320805 | 0.642705 | 0.190750 | H | 0.084845 | 2.138593 | 0.294407 |
| C | 0.915067 | -1.042568 | 0.183683 | H | -0.476610 | -2.544893 | 0.282073 |
| C | 2.015169 | -1.860497 | -0.080016 | H | 3.258509 | 2.571410 | -0.232765 |
| C | 3.265267 | -1.283600 | -0.309776 | | | | |

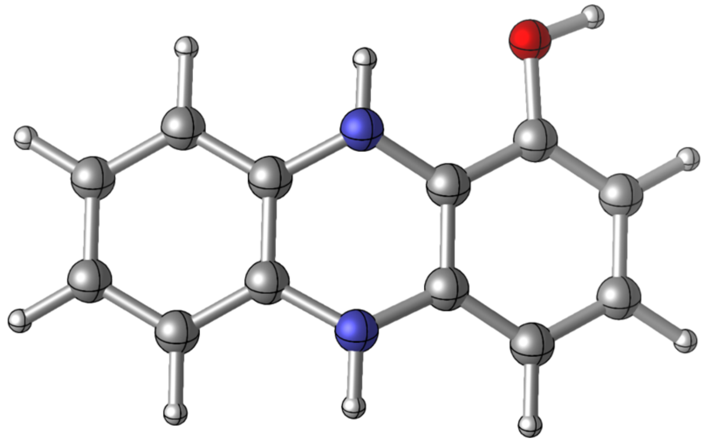

Zero-point correction= 0.199664 (Hartree/Particle)
Thermal correction to Energy= 0.210788
Thermal correction to Enthalpy= 0.211732
Thermal correction to Gibbs Free Energy= 0.162980
Sum of electronic and zero-point Energies= -647.677552
Sum of electronic and thermal Energies= -647.666429
Sum of electronic and thermal Enthalpies= -647.665484
Sum of electronic and thermal Free Energies= -647.714237

**OH-phenazine_oxidized_closed**
E(scf) = -646.663074024 a.u.

| | | | | | | | |
|---|---|---|---|---|---|---|---|
| C | 3.815118 | -0.525115 | 0.000043 | C | -3.272275 | -1.265953 | -0.000041 |
| C | 2.725707 | -1.348930 | 0.000029 | C | -3.426319 | 0.153139 | -0.000039 |
| C | 2.416814 | 1.462709 | 0.000031 | C | -2.319775 | 0.959691 | -0.000025 |
| C | 3.659748 | 0.896394 | 0.000044 | C | -1.007332 | 0.362463 | -0.000011 |
| H | 4.820217 | -0.949481 | 0.000054 | H | -1.932539 | -2.955674 | -0.000028 |
| H | 2.822682 | -2.435374 | 0.000028 | H | -4.174867 | -1.879652 | -0.000052 |
| H | 2.273882 | 2.544115 | 0.000031 | H | -4.420633 | 0.600290 | -0.000050 |
| H | 4.548724 | 1.528743 | 0.000055 | N | 0.035546 | 1.191033 | 0.000003 |
| C | 1.405106 | -0.797445 | 0.000015 | N | 0.349833 | -1.624652 | 0.000001 |
| C | 1.253618 | 0.632425 | 0.000016 | O | -2.399072 | 2.300025 | -0.000023 |
| C | -0.866359 | -1.063936 | -0.000012 | H | -1.476937 | 2.618587 | -0.000011 |
| C | -2.044987 | -1.871513 | -0.000027 | | | | |

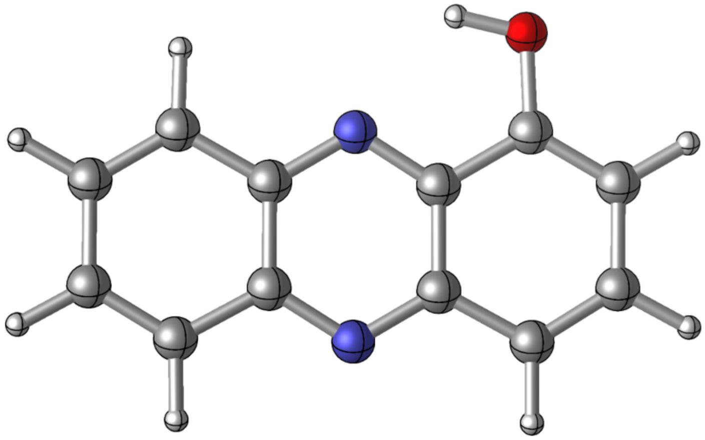

Zero-point correction= 0.176682 (Hartree/Particle)
Thermal correction to Energy= 0.186739
Thermal correction to Enthalpy= 0.187683
Thermal correction to Gibbs Free Energy= 0.140997
Sum of electronic and zero-point Energies= -646.486392
Sum of electronic and thermal Energies= -646.476335
Sum of electronic and thermal Enthalpies= -646.475391
Sum of electronic and thermal Free Energies= -646.522077

**OH-phenazine_oxidized_closed_octanol**
E(scf) = -646.662458149 a.u.

| | | | | | | | |
|---|---|---|---|---|---|---|---|
| C | 3.814895 | -0.525183 | 0.000043 | C | -3.271888 | -1.266000 | -0.000040 |
| C | 2.725411 | -1.348802 | 0.000029 | C | -3.426141 | 0.153058 | -0.000039 |
| C | 2.416695 | 1.462612 | 0.000031 | C | -2.319787 | 0.959764 | -0.000025 |
| C | 3.659578 | 0.896263 | 0.000044 | C | -1.007339 | 0.362510 | -0.000011 |
| H | 4.819960 | -0.949704 | 0.000054 | H | -1.931605 | -2.955615 | -0.000028 |
| H | 2.821551 | -2.435292 | 0.000028 | H | -4.174429 | -1.879806 | -0.000052 |
| H | 2.273560 | 2.543986 | 0.000031 | H | -4.420294 | 0.600489 | -0.000050 |
| H | 4.548538 | 1.528697 | 0.000055 | N | 0.035386 | 1.191240 | 0.000003 |
| C | 1.404869 | -0.797226 | 0.000015 | N | 0.349853 | -1.624607 | 0.000001 |
| C | 1.253438 | 0.632500 | 0.000016 | O | -2.398820 | 2.299936 | -0.000023 |
| C | -0.866180 | -1.063949 | -0.000012 | H | -1.476422 | 2.617508 | -0.000012 |
| C | -2.044713 | -1.871578 | -0.000027 | | | | |

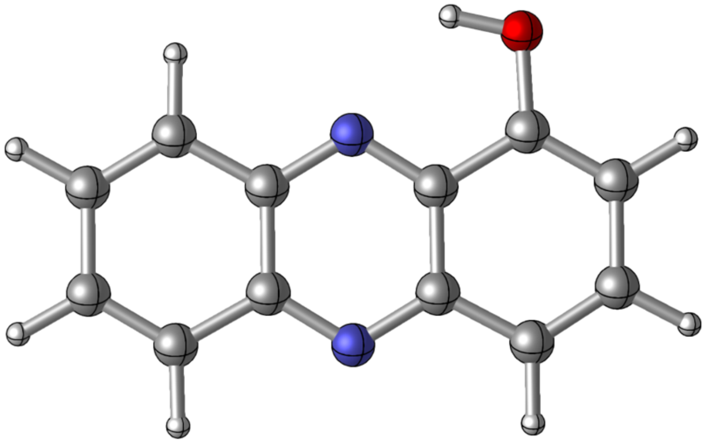

Zero-point correction= 0.176716 (Hartree/Particle)
Thermal correction to Energy= 0.186769
Thermal correction to Enthalpy= 0.187713
Thermal correction to Gibbs Free Energy= 0.141034
Sum of electronic and zero-point Energies= -646.485742
Sum of electronic and thermal Energies= -646.475689
Sum of electronic and thermal Enthalpies= -646.474745
Sum of electronic and thermal Free Energies= -646.521424

**OH-phenazine_oxidized_open**
E(scf) = -646.656242773 a.u.

| | | | | | | | |
|---|---|---|---|---|---|---|---|
| C | 3.838920 | -0.493013 | 0.000043 | C | -3.237008 | -1.320985 | -0.000040 |
| C | 2.755228 | -1.324309 | 0.000029 | C | -3.413361 | 0.093900 | -0.000039 |
| C | 2.425899 | 1.483666 | 0.000031 | C | -2.330632 | 0.935055 | -0.000025 |
| C | 3.673294 | 0.927766 | 0.000044 | C | -0.996213 | 0.379488 | -0.000011 |
| H | 4.847109 | -0.910280 | 0.000054 | H | -1.849078 | -2.970321 | -0.000027 |
| H | 2.859538 | -2.410205 | 0.000028 | H | -4.126830 | -1.952649 | -0.000052 |
| H | 2.273461 | 2.563874 | 0.000032 | H | -4.422946 | 0.509996 | -0.000050 |
| H | 4.557737 | 1.566571 | 0.000056 | N | 0.049726 | 1.207291 | 0.000003 |
| C | 1.431485 | -0.779973 | 0.000015 | N | 0.379206 | -1.608702 | 0.000001 |
| C | 1.265920 | 0.645898 | 0.000016 | O | -2.409479 | 2.279624 | -0.000023 |
| C | -0.838662 | -1.051134 | -0.000012 | H | -3.340569 | 2.539090 | -0.000033 |
| C | -1.995723 | -1.890223 | -0.000027 | | | | |

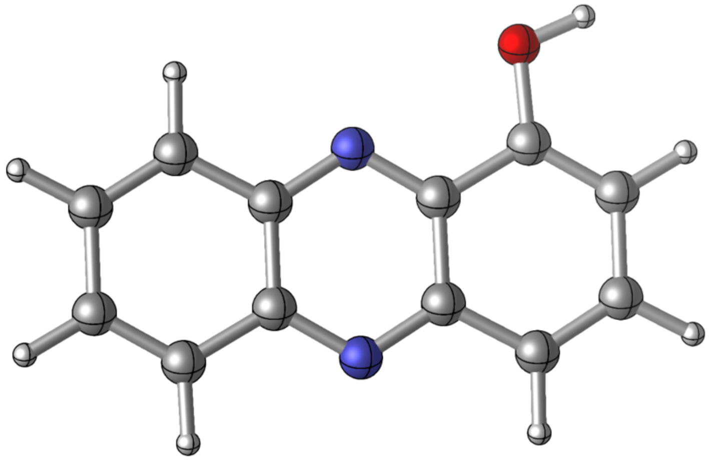

Zero-point correction= 0.176434 (Hartree/Particle)
Thermal correction to Energy= 0.186622
Thermal correction to Enthalpy= 0.187566
Thermal correction to Gibbs Free Energy= 0.140611
Sum of electronic and zero-point Energies= -646.479809
Sum of electronic and thermal Energies= -646.469621
Sum of electronic and thermal Enthalpies= -646.468677
Sum of electronic and thermal Free Energies= -646.515631

**OH-phenazine_oxidized_open_octanol**
E(scf) = -646.655107629 a.u.

| | | | | | | | |
|---|---|---|---|---|---|---|---|
| C | 3.838518 | -0.493037 | 0.000043 | C | -3.412912 | 0.093802 | -0.000039 |
| C | 2.754896 | -1.324305 | 0.000029 | C | -2.330305 | 0.935047 | -0.000025 |
| C | 2.425620 | 1.483655 | 0.000031 | C | -0.995851 | 0.379590 | -0.000011 |
| C | 3.672920 | 0.927718 | 0.000044 | H | -1.848074 | -2.970112 | -0.000027 |
| H | 4.846716 | -0.910384 | 0.000054 | H | -4.126434 | -1.952964 | -0.000052 |
| H | 2.858502 | -2.410239 | 0.000028 | H | -4.422642 | 0.509816 | -0.000050 |
| H | 2.272477 | 2.563727 | 0.000032 | N | 0.049741 | 1.207560 | 0.000003 |
| H | 4.557368 | 1.566578 | 0.000056 | N | 0.379168 | -1.608788 | 0.000001 |
| C | 1.431202 | -0.779924 | 0.000015 | O | -2.409384 | 2.279628 | -0.000023 |
| C | 1.265674 | 0.645895 | 0.000016 | H | -3.340448 | 2.538221 | -0.000033 |
| C | -0.838402 | -1.050944 | -0.000012 | | | | |
| C | -1.995481 | -1.890166 | -0.000027 | | | | |
| C | -3.236670 | -1.321178 | -0.000040 | | | | |

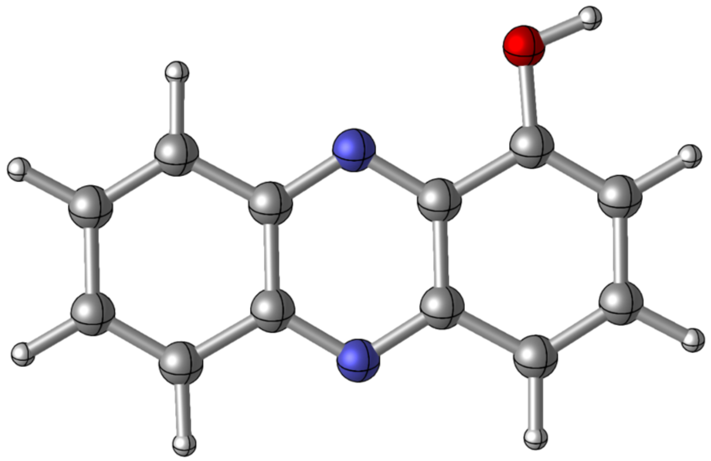

Zero-point correction= 0.176438 (Hartree/Particle)
Thermal correction to Energy= 0.186629
Thermal correction to Enthalpy= 0.187573
Thermal correction to Gibbs Free Energy= 0.140613
Sum of electronic and zero-point Energies= -646.478670
Sum of electronic and thermal Energies= -646.468479
Sum of electronic and thermal Enthalpies= -646.467535
Sum of electronic and thermal Free Energies= -646.514495

**LogP prediction algorithms**

We used four freely available, peer reviewed predictive algorithms relying on different methods of calculation to predict LogP values for neutral forms of the phenazines used in this study. We restricted our search to neutral forms of the molecules because several of these tools do not include charged molecules in their domains and thus give the same prediction for the protonated and deprotonated forms of, for example, PCA. iLOGP and XLogP3 were accessed through the SwissADME online tool[14]. The KOWWIN algorithm was accessed through the EPI (Estimation Program Interface) Suite[15].The molecular inspirations miLogP algorithm was accessed as a free tool on the molinspiration website, which has unfortunately been discontinued as of January 2026[16].

**cLogP Algorithm descriptions**

miLogP

miLogP is a free LogP prediction algorithm produced by Molinspiration[16]. It bases predictions on the contributions of known molecular fragments that it identifies within a molecule's structure. The empirical fragment contributions to lipophilicity are based on a training set of over 12,000 compounds, most of them drug-like, with measured LogP values. They include the influence of hydrogen bonding and charge interactions.

XLogP3

XLogP3 is based on the work of Wang et al., who developed an additive method based not on molecular fragments but on the individual atoms that constitute a molecule[17]. It includes contributions from element type, the identity of adjacent atoms, conjugation to a ring or extended pi system, and hybridization state. Summed atomic contributions are adjusted using correction factors to account for more complicated interactions attributable to halogens, intramolecular hydrogen bonds, etc. XLogP3 is the latest release of the software that, whenever possible, improves the accuracy of the original algorithm's calculations by using a structurally similar reference compound as a starting point for the additive, atom-based calculation[18].

KOWWIN:

Based on the work of Meylan and Howard[19-21], KOWWIN sums contributions from atoms and/or fragments identified within the molecule of interest. The 150 atom/fragment contributions recognized by the model were initially determined based on a training set of 2473 experimental LogP structure/value pairs for compounds with relatively simple structures. The model was then validated on an additional set of over 10,500 molecules of greater structural complexity. Correction factors account for interactions like steric hindrance and hydrogen bonding that might otherwise be missed by the atom/fragment approach.

iLOGP

iLOGP, short for "implicit log P", is a physics-based prediction method correlating experimental LogP measurements with computed solvation energies of analytes in octanol and in water[22]. It approximates contributions to solvation free energies in both solvents using two terms: 1) the electrostatic contribution, assessed using a generalized Born (GB) model to approximate solutions to the Poisson-Boltzman equation; and 2) the nonpolar contribution, approximated by the computed surface accessible free surface area (SASA). The resulting GB/SA model was then regressed against experimental values for thousands of molecules in a test set. The resulting model requires no corrective terms because it considers the geometry and polarity of the whole molecule, a fact which differentiates it wholly from atomistic or fragmental approaches like those listed above.

For each algorithm, predictions were based on the following SMILES sequences:
PCA: O=C(O)C1=CC=CC2=C1N=C3C(C=CC=C3)=N2 (oxidized) and
O=C(O)C1=CC=CC2=C1NC3=C(N2)C=CC=C3 (reduced);
PCN: O=C(N)C1=CC=CC2=C1N=C3C(C=CC=C3)=N2 (oxidized) and
O=C(N)C1=CC=CC2=C1NC3=C(N2)C=CC=C3 (reduced);
PYO: O=C1C=CC=C(C1=N2)N(C)C3=C2C=CC=C3 (oxidized) and
OC1=C2C(N(C)C3=CC=CC=C3N2)=CC=C1 (reduced);
1-OH-phenazine: OC1=CC=CC2=C1N=C3C(C=CC=C3)=N2 (oxidized) and
OC1=CC=CC2=C1NC3=C(N2)C=CC=C3 (reduced);
phenazine: C12=NC(C=CC=C3)=C3N=C1C=CC=C2 (oxidized) and
C1(NC(C=CC=C2)=C2N3)=C3C=CC=C1 (reduced).

### Experimental Methods
No unexpected or unusually high safety hazards were encountered.

#### Chemicals
Unless otherwise noted, all chemicals were purchased from Sigma. All solvents were HPLC grade. Formic acid and Acetonitrile were purchased from Fisher. All water used was nanopore grade (resistivity > 18 MΩ). PCN was purchased from ChemScene. PCA was purchased from Princeton Biomolecules. These molecules were used without further purification.

PYO was synthesized according to an established method with slight modifications[23]. Briefly, an aqueous solution of sodium bicarbonate and phenazine methosulfate was exposed to broad spectrum light in a light box for 16 hours at room temperature, causing decomposition to PYO. PYO was then purified in three extraction steps: first with DCM, then with 0.1 M HCl, and finally with DCM after neutralizing the acid with 10 M NaOH. DCM was removed by rotovap, and PYO was dissolved in a small volume of 80:20 DCM:MeOH solution. PYO was then precipitated by the addition of 200mL hexanes, and solids were vacuum filtered and washed with additional hexanes until flowthrough was clear. The identity and purity of PYO was confirmed against a lab standard by HPLC, and the powder was stored in the dark at -20 $^{o}$C for future experiments.
To ensure no significant decomposition, phenazine purity was assessed by HPLC at the start of each experiment.

#### Phenazine solutions
Phenazine stocks were all prepared in 20 mM MOPS at pH 7.2. For those that did not dissolve readily in water, we first dissolved a known amount of the solid in a known mass of methanol, then diluted by a factor of at least 25 using 20 mM MOPS. Small amounts of carrier solvent are not expected to influence partitioning behavior[24].

Because of observed side reactions between octanol and phenazines, especially when exposed to light, oxygen, a continuous reducing potential, or long-term storage, we did not saturate any phenazine stock solutions with octanol prior to partition experiments. However, we pre-saturated all octanol with 20 mM MOPS at pH 7.2. The amount of octanol that dissolves in water ($7.5 \times 10^{-5}$ mole fraction) is very small relative to the appreciable amount of water that dissolves in octanol (0.275 mole fraction)[25-27]. Thus, we assumed that volume changes due to small amounts of octanol dissolving in the aqueous phase during partition experiments would be negligible. Furthermore, our HPLC method is insensitive to volume changes in the equilibration experiment because it relies on the detection of analyte in both the octanol and the water phases.

#### Electrochemical Phenazine Reduction
All reduced phenazine stocks were prepared in well-stirred glass reaction vessels using solutions containing 0.1 mM to 1 mM phenazine in 20 mM MOPS at pH 7.2. Reducing potentials were chosen to be at least 250mV lower than the midpoint potential of the phenazine to be reduced. We then used a potentiostat to apply the chosen potential via a graphite rod working electrode and a Ag/AgCl (3M KCl) reference electrode. In all cases, we used a platinum counter electrode submerged in oxidized phenazine solution and connected to the working electrode vessel either by a glass frit (for PCA, PCN, and PYO), or by a salt bridge made by solidifying a mixture of 4% agar saturated with 3 M KCl in a serological pipette tip (for phenazine and 1-OH phenazine). Reduction was considered complete when the current measured at the reference electrode dropped below 1% of its initial value.

#### Equilibration Conditions

Exploring our desired condition space of octanol-water partitioning required the development of a high throughput equilibration and measurement method that could make use of small amounts of expensive or difficult-to-acquire analytes. We settled on a modified shake-flask method using sealable 2mL polypropylene eppendorf-style centrifuge tubes as reaction vessels and HPLC as an analytical method[28].

To minimize the influence of oxygen, we set up, equilibrated, and sampled all experiments in a Coy anaerobic chamber with an atmosphere of 95% $N_2$ and 5% $H_2$ . To perform an experiment, we used a syringe to dispense a small volume (typically 300µL, but volumes down to 150 µL in some cases) of octanol (pre-saturated with 20 mM MOPS at pH 7.2) into an eppendorf-style tube. Then, we used a pipette to dispense an aqueous phenazine solution (300µL) into the same tube. We then closed the tube and vortexed the mixture for 30s before placing it in a temperature-controlled shaker at 250 RPM for at least 16 hours, which was determined to be sufficient to reach equilibrium (Figure S14).

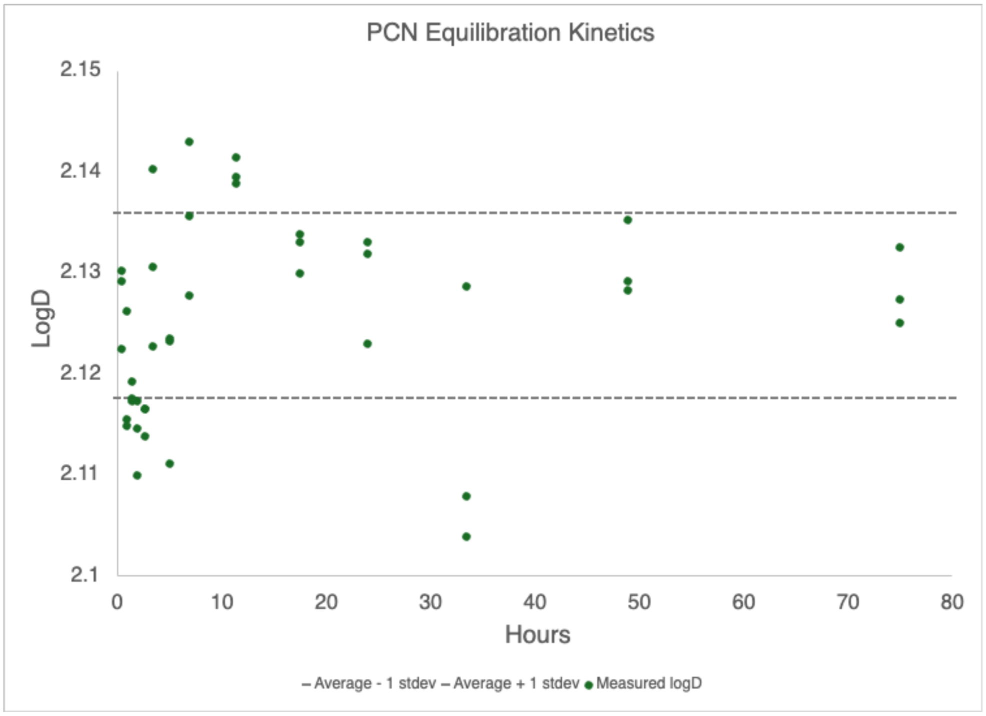

**Figure S14**. Control experiments to establish sufficient equilibration times with our experimental setup were carried out with oxidized PCN. Plotted points indicate individual replicates taken at each time point. Dashed lines indicate the average value from the last five time points plus or minus the standard deviation of those measurements. Based on these experiments, we determined that sampling after 16 hours was sufficient for PCA, PCN, and PYO. Out of an abundance of caution, we lengthened equilibration time to 24 hours for phenazine and 1-OH phenazine, which we worked with less frequently.

**Sampling Procedure**

To sample partition experiments after equilibration, we removed tubes from the heating block and centrifuged them for 90 s at 14,100xg using a microcentrifuge. We then collected about 100 µL of each of the fully separated octanol and water phases using separate 1 mL insulin syringes. Because the water phase sits below the octanol phase and small amounts of carryover can skew the measured partition coefficient significantly, we took great care to ensure no octanol contaminated the water samples. Before sampling the water phase, we pulled a small amount of air into the syringe. We then inserted the needle through the octanol phase, and, once the needle tip entered the water phase, ejected the air before drawing any of the water phase into the syringe. After withdrawing the needle and before dispensing the collected water phase into the HPLC vial for measurement, a small volume of the water phase was discarded to ensure no octanol remained at the tip of the syringe. All samples were measured in triplicate.

Because the reduced and oxidized forms of phenazines are known to absorb light differently, HPLC samples were removed from the Coy chamber and left uncapped under aluminum foil (to exclude light) on the benchtop for up to 2 hours prior to measurement to allow re-oxidation of reduced phenazine samples[29]. Samples were then capped and stored at 10$^{o}$C until measurement. Samples were measured within 3 days of equilibration to avoid complications from reaction with octanol that were occasionally observed, especially in the case of PCA.

**Liquid Chromatography Measurements**
All HPLC separations were performed on a Waters Alliance HPLC connected to a Waters 2998 PDA detector set to detect wavelengths from 200-800nm at 20Hz with 1.2nm resolution. To minimize errors in equilibration experiments, especially those associated with dispensing small volumes of octanol, we chose to measure analyte in both octanol and water phases. To avoid on-column interference from octanol in the samples, we used a phenyl column rather than a C18 column as well as a large proportion of methanol in the mobile phase, which helped to solubilize both phenazines and octanol. Even so, the retention times for the same analyte often differed between octanol and water fractions.

For equilibration experiments, we used Methods A, B, and C. A and B were both isocratic 1.0 mL/min flows of 70:30 MeOH: $H_2O$ with a phenyl column (XBridge BEH 5 µM particle size 4.6 x 250 mm), both solvents containing either 0.1% *v/v* formic acid (final pH ≤ 3, Method A) or 0.1% *v/v* $NH_4OH$ (final pH ≥ 9, Method C). Method C was an acidic gradient method using the same phenyl column used in Methods A and B, using solvents X ($H_2O$ + 0.1% *v/v* formic acid) and Y (70:30 MeOH: $H_2O$ + 0.1% *v/v* formic acid) flowing at 1mL/min according to the following gradient: 0-5 minutes: linear gradient from 95% X, 5% Y to 40% X, 60% Y; 5-8 minutes: linear gradient to 5% X, 95% Y; 8-10 minutes: linear gradient to 95% X, 5% Y.

For measurements of PYO in cell culture supernatant for the cell pellet retention assay, we used a C18 column (XBridge BEH 2.5µm particle size, 3 x 100mm) and the following basic gradient method (final pH ≥ 9, Method D), using solvent A ($H_2O$ + 2% *v/v* MeOH + 0.1% *v/v* $NH_4OH$) and Solvent B (acetonitrile + 2% *v/v* MeOH + 0.1% *v/v* $NH_4OH$) at 0.5 mL/min: 0-6 minutes: linear gradient from 98%A, 2%B to 80%A, 20%B; 6-8 minutes: linear gradient to 5%A, 95%B; 8-9 minutes hold at 5%A, 95%B; 9-9.1 minutes: linear gradient to 98%A, 2%B; 9.1-16 minutes: hold at 98%A, 2%B. A standard curve was prepared using the same method and was used to assess PYO concentrations down to 150nM. Prior to introducing them to the column, pellet supernatant samples were filtered using Costar SpinX centrifuge tube filters with 0.22 µm cellulose acetate filters.

| Phenazine | Sample Type | Method A (acidic, isocratic, phenyl column) | Method B (basic, isocratic, phenyl column) | Method C (acidic, gradient, phenyl column) | Method D (basic, gradient, C18 column) | Detection Wavelengths (nm) | Retention Time (min) |
|---|---|---|---|---|---|---|---|
| PCA | abiotic partitioning | X | | | | 364 | 8.4 |
| PCN | abiotic partitioning | X | | | | 287, 364 | 4.3 in oct. 4.5, 5.5[a] in water |
| PYO | abiotic partitioning | | X[b] | | | 313 | 3.6 |
| PYO | abiotic partitioning | | | X | | 387 | 6.6 |
| PYO | cell pellet and membrane retention | | | | X | 313 | 6.3 |
| Colchicine | abiotic partitioning control | X | | | | 352 | 5.0 |

**Table S5.** Samples and corresponding HPLC methods. a: multiple peaks attributed to PCN and its protonated rotamer in aqueous fraction (see the following methods section and Figures S2 and S3 for more information) b: Subsequent PYO samples were measured in basic conditions after recognizing an improvement in peak shapes compared to the acidic method.

Waters Empower 3 software was used to run the HPLC methods and to process the data. The built-in Apex Track Algorithm was used to extract chromatograms at wavelengths characteristic of each phenazine (Table 1). When strong partitioning resulted in extremely dilute water fractions, equilibration volumes and HPLC injection volumes were adjusted to ensure signals remained within the linear range of the detector. Peak areas in octanol and water, corrected for injection volume, were used to calculate distribution coefficients directly according to Equation 2. For each sample, the same wavelength was monitored in both octanol and water fractions.

**Treatment of PCN rotamer peak observed via HPLC**

In most of our samples, only one peak attributable to the analyte was visible. However, aqueous fractions of PCN samples often exhibited two peaks with different retention times but identical absorbance spectra characteristic of PCN. To determine whether both peaks were attributable to PCN, we used the same chromatography settings and PDA detector then routed the outflow through a Waters Acquity QDa single quadrupole mass spectrometer. Mass scans were collected simultaneously from 100 to 900 Da in the positive mode and 100 to 600 Da in the negative mode.

In addition, a selected ion recording channel was collected in the positive mode for 211.09, 224.08, and 225.07 Da, the masses expected for pyocyanin, phenazine-1-carboxamide, and phenazine-1-carboxylic acid, respectively. The probe temperature was 600°C and the capillary voltage was 0.8 kV. In all cases, one of the two peaks had a mass corresponding to PCN (~224.08 Da) while the other possessed a mass about one atomic mass unit greater than PCN (~224.94 Da). We speculate that the additional peaks are likely the result of a protonated rotamer preferentially stabilized in the water phase but not in the octanol phase (Figures S2 and S3). In these samples, we assumed that the 224.9 Da peak possessed the same absorption coefficient as the ~224.08 Da peak observed in the water and in the octanol phases. We thus calculated the distribution coefficient using the sum of the peak areas in the water fraction as the aqueous peak area (Equation 2).

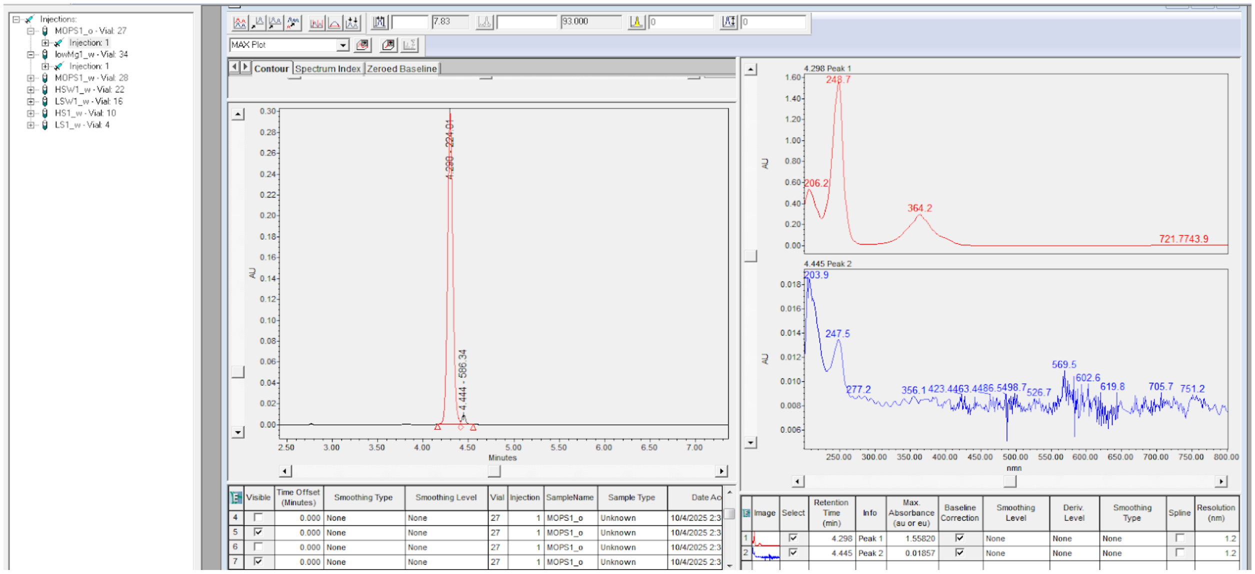

**Figure S15.** An example of a HPLC chromatogram extracted at 248nm to measure PCN in the octanol fraction of an equilibration experiment. The peak at 4.298 minutes has the absorbance spectrum (top right panel) and $[M+1]^+$ ion mass characteristic of protonated PCN (224.01Da). The small peak at 4.444 minutes has neither the correct mass nor the correct absorption spectrum for PCN. It was deemed an impurity, possibly related to the commercial synthesis of the PCN, and was not included in LogD calculations.

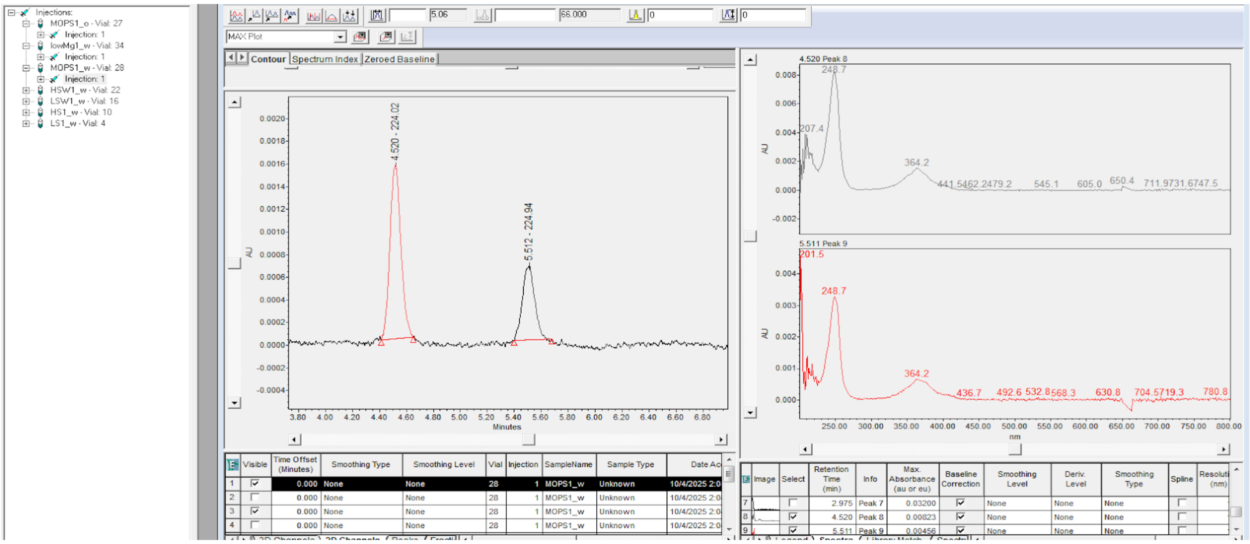

**Figure S16.** The HPLC chromatogram extracted at 248nm to measure PCN in the aqueous fraction of the same equilibration experiment sample as the panel above (Figure S2). The peak at 4.520 minutes has the characteristic absorbance spectrum and mass of the protonated PCN $[M+1]^+$ ion (224.02 Da) (top right panel), while the peak at 5.512 minutes has an essentially identical absorbance spectrum but a mass of 224.94 Da (bottom right panel). In this study, we attributed this peak to a rotamer of PCN stabilized in water but not in octanol, which would help justify its similar absorbance spectrum while accounting for its different retention time and mass. We assumed the absorption coefficient was not significantly different from PCN, and thus added the two measured peak areas in the water fraction together when calculating the distribution coefficient LogD according to Figure S17.

$$LogD = Log_{10} \sum_{i}^{n} \frac{[peak\ area\ octanol]_i}{[peak\ area\ water]_i} * \frac{injection\ volume\ water}{injection\ volume\ octanol}$$

**Figure S17**. Equation used to calculate octanol-water distribution coefficients for an analyte with n chromatographic peaks. For all samples, the same wavelength was monitored in octanol and water fractions. In rare cases when multiple peaks were present, mass and absorption spectra were used together to select and sum only peaks attributable to the intended analyte in its different ionization/rotamer forms and to exclude any peaks from impurities.

**Ionic Strength Samples**

To assess the effects of ionic identity and strength on phenazine partitioning, we used single-salt solutions at 0.1 M and 1 M ionic strengths. To check for constructive or destructive influences of multiple salts in an environmentally relevant mixture, we used a solution of artificial seawater (minus carbonate and trace metals, to avoid interference) consisting of the following salts and concentrations: $MgCl_2$, 46.6 mM; $CaCl_2$, 1.36 mM; NaCl, 456.88 mM; KCl, 7.00 mM; $Na_2SO_4$, 10.00 mM; $K_2HPO_4$, 1.00 mM; $NH_4Cl$, 2.00 mM. The ionic strength of the artificial seawater solution was 0.7 M, and we also used a 10x dilution of the same salt mixture (0.07 M ionic strength) to check for concentration effects.

Prior to equilibration with octanol, phenazine stock solutions were amended with volumes (up to 1/3rd of the final aqueous sample volume) of concentrated salt solutions prepared in water. pH changes were monitored with pH paper. Only the phosphate samples demonstrated a significant deviation from the buffered pH of 7.2 to a final pH of 8.5, a change which, based on the distance from the relevant pKa values, should not significantly affect the percentage of protonated and deprotonated species for any of the phenazines studied.

Samples were then equilibrated at 25 °C with shaking, sampled by syringe, and measured by HPLC as described above.

**pH Samples**

Phenazine stock solutions were prepared as above, buffered with 20 mM MOPS. To adjust the pH, small volumes (up to 1/30$^{th}$ of the final aqueous sample volume) of 1 M HCl or NaOH were added to overwhelm the buffer, and pH was measured to the nearest 0.5 pH units in the anaerobic chamber using pH paper. When the pH was lowered to 3, PCA samples precipitated in the aqueous phase, but crystals dissolved fully when the octanol phase was introduced.

**PYO Retention Samples**

**Experiment 1: cell pellets washed under normoxic or anoxic conditions**

To assess the biological implications of the measured redox sensitivity of PYO's LogD, we conducted an experiment with late stationary phase *P. aeruginosa* cultures. *P. aeruginosa* PA14 cells were grown to late stationary phase (20 hours) in 5 mL of LB at 37 °C with vigorous shaking to ensure aeration. For each of three biological replicates, two 1 mL aliquots were taken from the same culture tube and washed either aerobically or anaerobically. While the aerobic aliquot was washed three times with PBS on the benchtop, the anaerobic aliquot was allowed to rest in an anaerobic chamber for 45 minutes, allowing the cells to reduce the PYO in the supernatant (a process that is visually apparent as the blue, oxidized PYO transforms to its colorless reduced form). Then the anaerobic aliquot was washed three times in the anaerobic chamber with $N_2$-sparged PBS. One wash consisted of spinning the cells down at 10,000xg for 1 minute, removing the supernatant, and resuspending the cell pellet in fresh PBS. After three washes, both the aerobic and the anaerobic cells were washed once more with aerobic PBS on the benchtop. In all conditions, supernatant was collected from the original culture, from the third wash, and from the fourth, aerobic wash. Supernatant samples were then analyzed by HPLC using method C listed in HPLC Conditions. Concentrations were calculated using a calibration curve constructed using pure PYO (Figure S4).

**Experiment 2: membrane PYO retention measurement**

A Pseudomonas aeruginosa culture (1 L) was grown under aerobic conditions in LB medium at 37 °C with shaking at 250 RPM to stationary phase ($OD_{500}$ = 4.7). Cells were allowed to stand on the benchtop for 5 minutes before centrifugation, during which time the color of the culture changed from blue (attributable to oxidized pyocyanin) to colorless (suggesting pyocyanin was reduced). Cells were harvested by centrifugation at 8000 RPM for 10 minutes. Culture supernatant was collected, mixed 1:1 with methanol, and stored at −20 °C until measurement. Cell pellets were resuspended in PBS (pH 7.0) supplemented with DNase I and lysed using an Avestin Emulsiflex C3 homogenizer (three passes at 15,000 PSI). The resulting lysate was subjected to ultracentrifugation at 38,000 RPM for 50 minutes at 4 °C to separate soluble and membrane fractions. Phenazines were extracted from the membrane fraction by adding 20 mL of methanol followed by vigorous vortexing and ultracentrifugation at 38,000 RPM for 10 minutes. During the first extraction, the color of the pellet changed from colorless (tinged with red, likely because of cytochrome hemes) to blue (attributable to oxidized pyocyanin). This extraction procedure was repeated three times, at which point both the membrane pellet and the methanol were colorless. The three resulting methanol fractions were pooled and evaporated on a rotovap to a volume of 4.7mL prior to LC–MS analysis. During the drying process, solids appeared in the solvent. These presumed protein aggregates were separated via centrifugation at 5300 RPM for 10 minutes, and the resulting supernatant was diluted 1:10 with fresh methanol prior to measurement. Medium and membrane fraction extract were both measured by HPLC method D described above, and concentrations were calculated based on the same standard curve as in PYO retention experiment 1 (Figure S4).

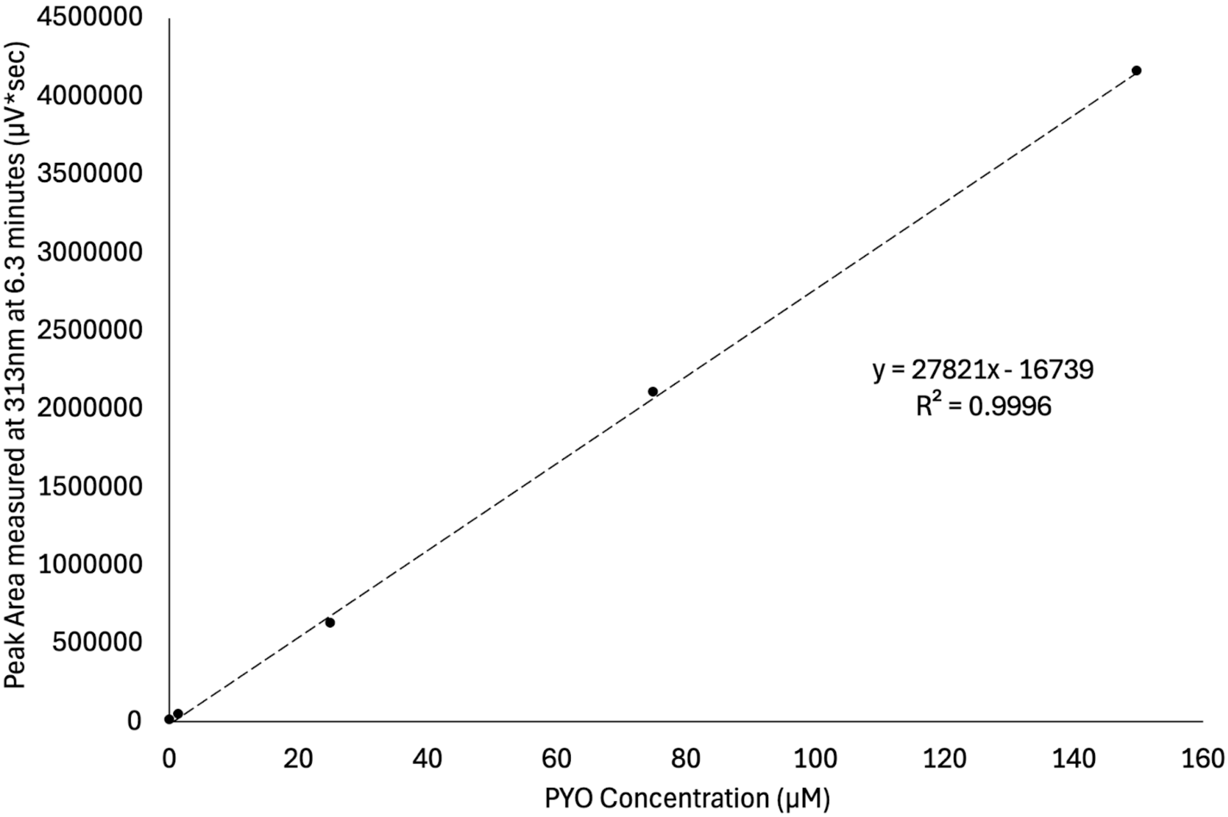

**Figure S17.** Standard curve used to calculate pyocyanin concentrations in supernatant samples from cellular retention assay and membrane retention assay. Standard solutions were prepared at concentrations of 150, 75, 25, 1.5, and 0.15 µM.

**Ionic context exerts only minor control on phenazine lipophilicity**

For most organic solutes, increasing ionic strength in the aqueous phase induces a "salting out" effect, increasing the observed lipophilicity. For the purposes of this discussion, we consider a "normal" salting out effect to be an increase in lipophilicity with an increase in ionic strength, while a "reverse" salting out effect is a decrease in lipophilicity with increasing ionic strength. Variations with a 10-fold change in ionic strength were generally small for PCA, PCN, and PYO. For all the salts tested, PCN exhibited a normal salting out effect (Figure S18 A). While the differences in the lipophilicities are small enough that we hesitate to overinterpret them, PYO and PCA showed slightly different behavior when treated with different salts, only sometimes adhering to normal salting out expectations (Figure S18 B, C). Additionally, when treated with NaCl or KCl, the change in lipophilicity with increasing ionic strength was normal for the reduced species but reversed for the oxidized species. Particularly in the case of potassium phosphate, higher ionic strength actually made both oxidized and reduced PCA appreciably less lipophilic. This in spite of the fact that the phosphate salt overwhelmed the pH 7.2 MOPS buffer, resulting in an experimental pH of 8.5, which should have increased the proportion of deprotonated, charged PCA. Although not strong chelators, PCA and PYO are both known to bind to certain metals. Chelation of trace impurities in the salts may have contributed to the deviations we noted, but the effects were mostly small. Due to uncontrolled redox interactions and subsequent precipitation, we were unable to precisely characterize the effects of $Cu^{2+}$, $Fe^{2+}$, or $Fe^{3+}$ in this study.

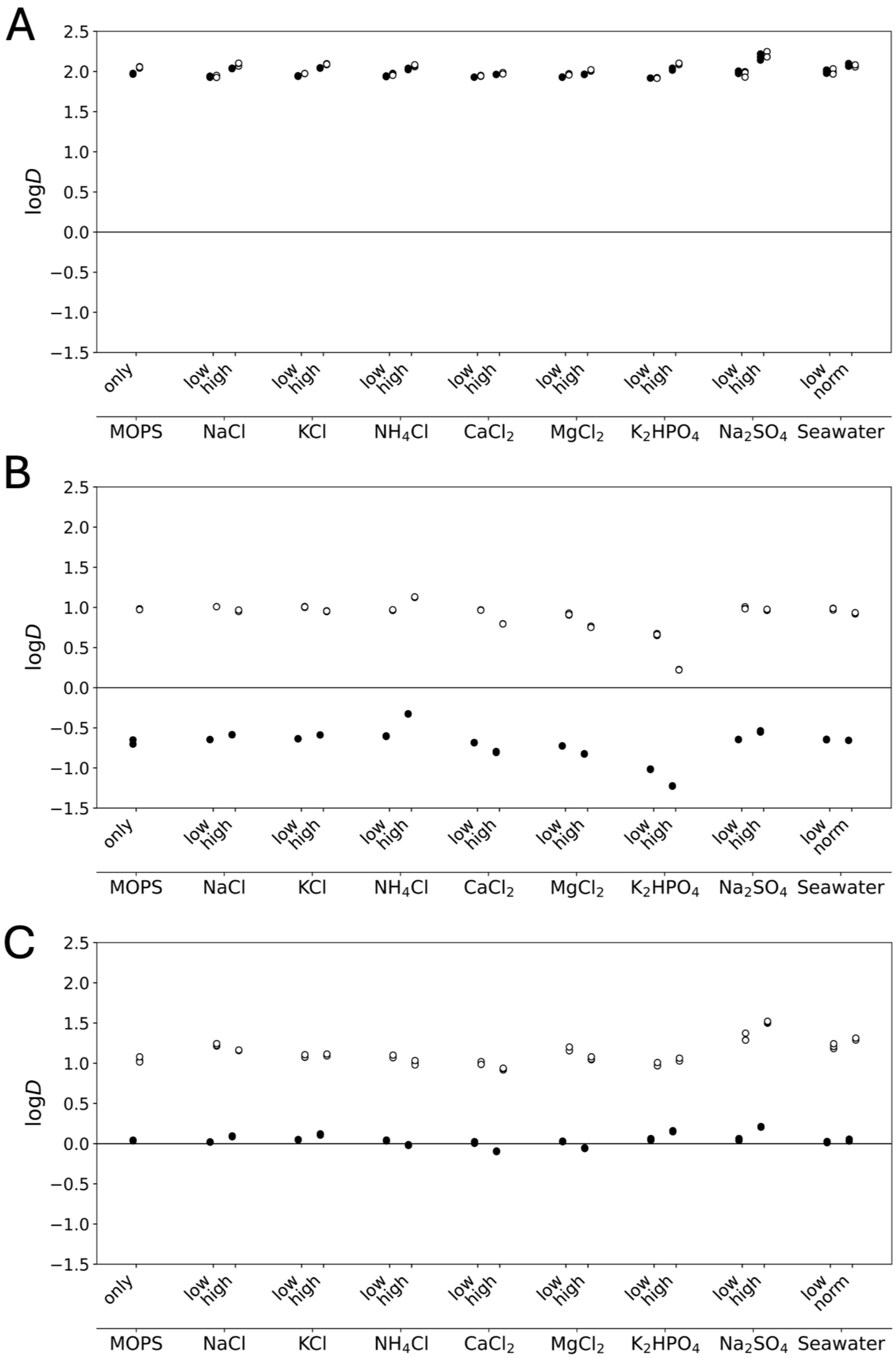

**Figure S18.** LogD measurements in various ionic strength conditions for PCN (A), PCA (B), and PYO (C). Ionic strength equilibrations were carried out at room temperature. In all cases, the aqueous fraction contained 20 mM MOPS at pH 7.2 amended to achieve final concentrations of either 0.1 M or 1 M concentrations of the salts listed. The phosphate salt overwhelmed the MOPS buffer and reached pH 8.5. All other salts were circumneutral at experiment's end. Each condition was conducted in triplicate (all data is shown). A table of exact experimental values is available below.

**Tables of Experimental Values**
In each table below, a line represents an average of experimental triplicates. All experiments were carried out in 20mM MOPS.

Colchicine method validation (pH 7.2)

| Compound | Redox State | pH | $logD_{7.2}$ (average of triplicates) | Standard Error (SE) | 1.96*SE |
|---|---|---|---|---|---|
| Colchicine | N/A | 7.2 | 1.06 | 0.002 | 0.004 |
| Colchicine | N/A | 7.2 | 1.06 | 0.004 | 0.008 |

**Table S6.** Method validation using Colchicine.

Phenazine and 1-OH phenazine (pH 7.2)

| Compound | Redox State | pH | $logD_{7.2}$ (average of triplicates) | Standard Error (SE) | 1.96*SE |
|---|---|---|---|---|---|
| phenazine | oxidized | 7.2 | 2.66 | 0.015 | 0.030 |
| **phenazine** | **reduced** | **7.2** | **2.63** | **0.037** | **0.073** |
| 1-OH phenazine | oxidized | 7.2 | 2.65 | 0.019 | 0.037 |
| 1-OH phenazine | **reduced** | 7.2 | **2.45** | **0.003** | **0.005** |

**Table S7.** Experimental values for phenazine and 1-OH phenazine.

Ionic Strength Experiments (pH 7.2)

| Phenazine | Redox State | Salt | Salt Conc. (M) | $logD_{7.2}$ (average of triplicates) | Standard Error (SE) | 1.96*SE |
|---|---|---|---|---|---|---|
| PCA | oxidized | NaCl | 0.1 | -0.65 | 0.001 | 0.002 |
| PCA | oxidized | NaCl | 1 | -0.59 | 0.001 | 0.002 |
| PCA | oxidized | KCl | 0.1 | -0.64 | 0.002 | 0.003 |
| PCA | oxidized | KCl | 1 | -0.59 | 0.001 | 0.002 |

| | | | | | | |
|---|---|---|---|---|---|---|
| PCA | oxidized | CaCl2 | 0.1 | -0.68 | 0.002 | 0.004 |
| PCA | oxidized | CaCl2 | 1 | -0.80 | 0.004 | 0.008 |
| **PCA** | **reduced** | **NaCl** | **0.1** | **1.01** | **0.001** | **0.001** |
| **PCA** | **reduced** | **NaCl** | **1** | **0.96** | **0.006** | **0.011** |
| **PCA** | **reduced** | **KCl** | **0.1** | **1.01** | **0.003** | **0.007** |
| **PCA** | **reduced** | **KCl** | **1** | **0.95** | **0.003** | **0.006** |
| **PCA** | **reduced** | **CaCl2** | **0.1** | **0.97** | **0.002** | **0.004** |
| **PCA** | **reduced** | **CaCl2** | **1** | **0.80** | **0.001** | **0.001** |
| PCA | oxidized | MgCl2 | 0.1 | -0.72 | 0.001 | 0.003 |
| PCA | oxidized | MgCl2 | 1 | -0.82 | 0.002 | 0.004 |
| PCA | oxidized | NH4Cl | 0.1 | -0.60 | 0.003 | 0.005 |
| PCA | oxidized | NH4Cl | 1 | -0.33 | 0.001 | 0.003 |
| PCA | oxidized | K2HPO4 | 0.1 | -1.02 | 0.002 | 0.005 |
| PCA | oxidized | K2HPO4 | 1 | -1.22 | 0.002 | 0.004 |
| **PCA** | **reduced** | **MgCl2** | **0.1** | **0.92** | **0.007** | **0.015** |
| **PCA** | **reduced** | **MgCl2** | **1** | **0.76** | **0.005** | **0.010** |
| **PCA** | **reduced** | **NH4Cl** | **0.1** | **0.97** | **0.002** | **0.005** |
| **PCA** | **reduced** | **NH4Cl** | **1** | **1.13** | **0.002** | **0.005** |
| **PCA** | **reduced** | **K2HPO4** | **0.1** | **0.66** | **0.007** | **0.014** |
| **PCA** | **reduced** | **K2HPO4** | **1** | **0.22** | **0.002** | **0.004** |
| PCA | oxidized | Na2SO4 | 0.1 | -0.65 | 0.002 | 0.003 |
| PCA | oxidized | Na2SO4 | 1 | -0.54 | 0.005 | 0.010 |
| PCA | oxidized | artificial seawater | 1/3 strength | -0.64 | 0.003 | 0.005 |
| PCA | oxidized | artificial seawater | normal | -0.66 | 0.001 | 0.001 |
| PCA | oxidized | MOPS only | N/A | -0.67 | 0.018 | 0.035 |
| **PCA** | **reduced** | **Na2SO4** | **0.1** | **0.99** | **0.008** | **0.016** |
| **PCA** | **reduced** | **Na2SO4** | **1** | **0.97** | **0.004** | **0.008** |
| **PCA** | **reduced** | **artificial seawater** | **1/3 strength** | **0.98** | **0.007** | **0.013** |
| **PCA** | **reduced** | **artificial seawater** | **normal** | **0.93** | **0.005** | **0.009** |
| **PCA** | **reduced** | **MOPS only** | **N/A** | **0.98** | **0.004** | **0.008** |
| PYO | oxidized | NaCl | 0.1 | 0.02 | 0.001 | 0.003 |
| PYO | oxidized | NaCl | 1 | 0.09 | 0.004 | 0.007 |

| | | | | | | |
|---|---|---|---|---|---|---|
| PYO | oxidized | KCl | 0.1 | 0.05 | 0.002 | 0.004 |
| PYO | oxidized | KCl | 1 | 0.12 | 0.005 | 0.009 |
| PYO | oxidized | CaCl2 | 0.1 | 0.02 | 0.007 | 0.014 |
| PYO | oxidized | CaCl2 | 1 | -0.09 | 0.003 | 0.006 |
| **PYO** | **reduced** | **NaCl** | **0.1** | **1.23** | **0.009** | **0.018** |
| **PYO** | **reduced** | **NaCl** | **1** | **1.16** | **0.002** | **0.005** |
| **PYO** | **reduced** | **KCl** | **0.1** | **1.09** | **0.011** | **0.021** |
| **PYO** | **reduced** | **KCl** | **1** | **1.11** | **0.008** | **0.015** |
| **PYO** | **reduced** | **CaCl2** | **0.1** | **1.00** | **0.011** | **0.021** |
| **PYO** | **reduced** | **CaCl2** | **1** | **0.93** | **0.006** | **0.013** |
| PYO | oxidized | MgCl2 | 0.1 | 0.03 | 0.003 | 0.005 |
| PYO | oxidized | MgCl2 | 1 | -0.06 | 0.003 | 0.006 |
| PYO | oxidized | NH4Cl | 0.1 | 0.04 | 0.004 | 0.008 |
| PYO | oxidized | NH4Cl | 1 | -0.02 | 0.004 | 0.007 |
| PYO | oxidized | K2HPO4 | 0.1 | 0.05 | 0.007 | 0.013 |
| PYO | oxidized | K2HPO4 | 1 | 0.16 | 0.004 | 0.008 |
| **PYO** | **reduced** | **MgCl2** | **0.1** | **1.17** | **0.015** | **0.030** |
| **PYO** | **reduced** | **MgCl2** | **1** | **1.06** | **0.011** | **0.022** |
| **PYO** | **reduced** | **NH4Cl** | **0.1** | **1.08** | **0.010** | **0.020** |
| **PYO** | **reduced** | **NH4Cl** | **1** | **1.01** | **0.016** | **0.031** |
| **PYO** | **reduced** | **K2HPO4** | **0.1** | **0.98** | **0.013** | **0.026** |
| **PYO** | **reduced** | **K2HPO4** | **1** | **1.04** | **0.012** | **0.023** |
| PYO | oxidized | Na2SO4 | 0.1 | 0.05 | 0.008 | 0.015 |
| PYO | oxidized | Na2SO4 | 1 | 0.21 | 0.003 | 0.006 |
| PYO | oxidized | artificial seawater | 1/3 strength | 0.02 | 0.006 | 0.011 |
| PYO | oxidized | artificial seawater | normal | 0.04 | 0.008 | 0.016 |
| PYO | oxidized | MOPS only | N/A | 0.04 | 0.003 | 0.006 |
| **PYO** | **reduced** | **Na2SO4** | **0.1** | **1.32** | **0.028** | **0.056** |
| **PYO** | **reduced** | **Na2SO4** | **1** | **1.51** | **0.007** | **0.013** |
| **PYO** | **reduced** | **artificial seawater** | **1/3 strength** | **1.21** | **0.018** | **0.034** |
| **PYO** | **reduced** | **artificial seawater** | **normal** | **1.30** | **0.007** | **0.014** |
| **PYO** | **reduced** | **MOPS only** | **N/A** | **1.04** | **0.019** | **0.038** |

| | | | | | | |
|---|---|---|---|---|---|---|
| PCN | oxidized | NaCl | 0.1 | 1.94 | 0.005 | 0.009 |
| PCN | oxidized | NaCl | 1 | 2.04 | 0.003 | 0.005 |
| PCN | oxidized | KCl | 0.1 | 1.94 | 0.003 | 0.005 |
| PCN | oxidized | KCl | 1 | 2.05 | 0.002 | 0.004 |
| PCN | oxidized | CaCl2 | 0.1 | 1.93 | 0.001 | 0.001 |
| PCN | oxidized | CaCl2 | 1 | 1.96 | 0.001 | 0.002 |
| **PCN** | **reduced** | **NaCl** | **0.1** | **2.44** | **0.008** | **0.015** |
| **PCN** | **reduced** | **NaCl** | **1** | **2.57** | **0.011** | **0.022** |
| **PCN** | **reduced** | **KCl** | **0.1** | **2.45** | **0.000** | **0.001** |
| **PCN** | **reduced** | **KCl** | **1** | **2.57** | **0.004** | **0.007** |
| **PCN** | **reduced** | **CaCl2** | **0.1** | **2.42** | **0.004** | **0.007** |
| **PCN** | **reduced** | **CaCl2** | **1** | **2.46** | **0.005** | **0.010** |
| PCN | oxidized | MgCl2 | 0.1 | 1.93 | 0.002 | 0.003 |
| PCN | oxidized | MgCl2 | 1 | 1.96 | 0.002 | 0.003 |
| PCN | oxidized | NH4Cl | 0.1 | 1.94 | 0.003 | 0.007 |
| PCN | oxidized | NH4Cl | 1 | 2.03 | 0.006 | 0.011 |
| PCN | oxidized | K2HPO4 | 0.1 | 1.92 | 0.001 | 0.002 |
| PCN | oxidized | K2HPO4 | 1 | 2.03 | 0.009 | 0.017 |
| **PCN** | **reduced** | **MgCl2** | **0.1** | **2.44** | **0.005** | **0.010** |
| **PCN** | **reduced** | **MgCl2** | **1** | **2.49** | **0.006** | **0.011** |
| **PCN** | **reduced** | **NH4Cl** | **0.1** | **2.44** | **0.007** | **0.014** |
| **PCN** | **reduced** | **NH4Cl** | **1** | **2.53** | **0.003** | **0.006** |
| **PCN** | **reduced** | **K2HPO4** | **0.1** | **2.40** | **0.003** | **0.006** |
| **PCN** | **reduced** | **K2HPO4** | **1** | **2.42** | **0.006** | **0.011** |
| PCN | oxidized | Na2SO4 | 0.1 | 1.99 | 0.010 | 0.019 |
| PCN | oxidized | Na2SO4 | 1 | 2.18 | 0.023 | 0.044 |
| PCN | oxidized | artificial seawater | 1/3 strength | 2.00 | 0.012 | 0.024 |
| PCN | oxidized | artificial seawater | normal | 2.08 | 0.011 | 0.022 |
| PCN | oxidized | MOPS only | N/A | 1.97 | 0.004 | 0.007 |
| **PCN** | **reduced** | **Na2SO4** | **0.1** | **2.45** | **0.021** | **0.041** |
| **PCN** | **reduced** | **Na2SO4** | **1** | **2.68** | **0.022** | **0.043** |
| **PCN** | **reduced** | **artificial seawater** | **1/3 strength** | **2.48** | **0.019** | **0.038** |
| | **reduced** | **artificial seawater** | **normal** | **2.55** | **0.008** | **0.015** |

| | | | | | | |
|---|---|---|---|---|---|---|
| **PCN** | | | | | | |
| **PCN** | **reduced** | **MOPS only** | **N/A** | **2.53** | **0.005** | **0.011** |

**Table S8.** Ionic strength experiments were conducted by adding small amounts of concentrated salt solutions to 20mM MOPS buffered phenazine stock solutions. Phosphate solutions overwhelmed the buffer and became more basic (pH 8.5), but this was not expected to interfere with the ionization state of the phenazines used in this portion of the study.

pH Experiments

| Phenazine | Redox State | pH | $logD_{pH}$ (average of triplicates) | Standard Error (SE) | 1.96*SE |
|---|---|---|---|---|---|
| PYO | oxidized | 3.0 | -2.21 | 0.013 | 0.026 |
| PYO | oxidized | 6.0 | -0.12 | 0.003 | 0.005 |
| PYO | oxidized | 9.0 | 0.03 | 0.004 | 0.009 |
| **PYO** | **reduced** | **3.0** | **0.42** | **0.020** | **0.039** |
| **PYO** | **reduced** | **6.0** | **1.13** | **0.100** | **0.196** |
| **PYO** | **reduced** | **9.0** | **1.02** | **0.053** | **0.104** |
| PCA | oxidized | 3 | 2.12 | 0.006 | 0.011 |
| PCA | oxidized | 3 | 2.12 | 0.004 | 0.009 |
| PCA | oxidized | 9 | -2.01 | 0.014 | 0.028 |
| PCA | oxidized | 4 | 2.11 | 0.039 | 0.076 |
| PCA | oxidized | 6 | -0.09 | 0.001 | 0.002 |
| **PCA** | **reduced** | **3** | **3.42** | **0.046** | **0.091** |
| **PCA** | **reduced** | **5.5** | **1.54** | **0.017** | **0.033** |
| **PCA** | **reduced** | **9.5** | **-1.08** | **0.007** | **0.014** |
| **PCA** | **reduced** | **4.5** | **3.35** | **0.049** | **0.097** |
| PCN | oxidized | 3 | 2.13 | 0.010 | 0.020 |
| PCN | oxidized | 9.5 | 1.94 | 0.008 | 0.016 |
| **PCN** | **reduced** | **2.5** | **2.26** | **0.016** | **0.031** |
| **PCN** | **reduced** | **9** | **2.10** | **0.001** | **0.002** |

**Table S9.** For pH experiments, 20mM MOPS starting solutions were overwhelmed with small amounts of concentrated HCl or NaOH. Values from the Ionic Strength MOPS only condition (Table S8) were used to assess LogD at pH 7.2.

**Intracellular pyocyanin concentration estimation**

Goal:
This calculation is designed to estimate a lower bound for intracellular $PYO_{reduced}$ concentrations in a reduced subsample by comparing measured concentrations of extracellular PYO in oxidized and reduced subsamples of the same starter culture.

Calculation:

1. The intracellular concentration of PYO in the reduced subsample is defined as the intracellular amount of PYO in moles divided by the intracellular volume:
$$[PYO]_{reduced,\,intracellular} = \frac{nPYO_{reduced,\,intracellular}}{V_{reduced,\,intracellular}}$$
2. The intracellular volume is the product of the cellular volume (L/cell), the culture density (cells/L) and the subsample volume (L):
$$V_{reduced,\,intracellular} = V_{cell} * \rho_{culture} * V_{subsample}$$
3. To calculate the intracellular moles of PYO in the reduced subsample, we begin with an assumption. Because both subsamples are taken from the same starter culture and are not allowed sufficient time to produce significant additional PYO (45 min for sample preparation), we assume that oxidized and reduced subsamples contain the same total moles of PYO:
$nPYO_{reduced} = nPYO_{oxidized}$
4. Our HPLC measurements show that, for a given starter culture, oxidized subsamples contain significantly more extracellular PYO than reduced subsamples. For simplicity, we will assume that oxidized subsamples contain only extracellular PYO:
$nPYO_{oxidized} = nPYO_{oxidized,\,extracellular}$.
5. And we will assume that reduced subsamples contain both extracellular and intracellular PYO:
$nPYO_{reduced} = nPYO_{reduced,\,extracellular} + nPYO_{reduced,\,intracellular}$ .

This simplifying assumption will ultimately result in an underestimation of intracellular PYO if, as is likely, oxidized subsamples also contain some intracellular PYO.
Because no PYO degradation mechanism is currently known in *P. aeruginosa PA14*, we then assume that the PYO deficit consistently observed in the reduced condition compared to the oxidized condition is due solely to uptake (whether due to passive partitioning and/or active processes). In other words, the difference between measured extracellular amounts in the two subsamples is exactly the amount of intracellular PYO in the reduced condition:

6. $nPYO_{oxidized,\,extracellular} - nPYO_{reduced,\,extracellular} = nPYO_{reduced,\,intracellular}$

Molar amounts of extracellular PYO in an oxidized or reduced subsample can be calculated directly from the concentrations measured by HPLC and the volumes of the subsamples:

7. $[PYO]_{subsample,\,extracellular} * V_{subsample} = nPYO_{subsample,\,extracellular}$

Substituting Equation 7 into Equation 6 gives the total moles of intracellular PYO:

8. 
$$nPYO_{reduced,\,intracellular} = [PYO]_{oxidized,\,extracellular} * V_{oxidized} - [PYO]_{reduced,\,extracellular} * V_{reduced}$$

The intracellular PYO concentration in the reduced subsample is then given by substituting Equation 8 and Equation 2 into Equation 1 as numerator and denominator, respectively:

9.
$$[PYO]_{reduced,\, intracellular} = \frac{[PYO]_{oxidized,\, extracellular} * V_{oxidized} - [PYO]_{reduced,\, extracellular} * V_{reduced}}{V_{cell} * \rho_{culture} * V_{reduced}}$$

Quantities used for calculation:

- A dense culture of WT PA14 cells (measured $OD_{500}$ = 7.5) typically contains about $7x10^9$ cells per mL, or $7x10^{12}$ cells per L [30].
- The volume of each cell is approximately one cubic micron, or one 1fL ($10^{-15}$L) [31].
- The average extracellular concentration in oxidized subsamples was 60μM.
- The average extracellular concentration in reduced subsamples was 35μM.
- The volumes of oxidized and reduced subsamples were 1mL

Substituting, we find the moles of intracellular PYO in the reduced sample (Equation 8) is

$$nPYO_{intracellular,reduced} = (60 \times 10^{-6} M) * (1 \times 10^{-3} L) - (35 \times 10^{-6} M) * (1 \times 10^{-3} L)$$

$$nPYO_{intracellular,\, reduced} = 25 \times 10^{-9} moles.$$

The intracellular volume in the reduced fraction (Equation 2) is

$$V_{intracellular,\, reduced} = V_{cell} * \rho_{culture} * V_{reduced} = \left(1 \times 10^{-15}\ \frac{L}{cell}\right) * \left(7 \times 10^{12}\ \frac{cells}{L}\right) * (1 \times 10^{-3}\ L)\ V_{intracellular,\, reduced} = 7 \times 10^{-6}\ L.$$

The intracellular concentration is then

$$[PYO]_{intracellular,\, reduced} = \frac{nPYO_{intracellular,\, reduced}}{V_{intracellular,\, reduced}} = \frac{25 \times 10^{-9} moles}{7 \times 10^{-6} L} = 3.6 mM.$$

**Membrane fraction PYO retention experiment calculation**

Goal: These calculations are designed to determine the concentration of PYO on a per gram basis in the media and in the recovered membrane fraction of a dense *P. aeruginosa* culture.

Calculations:

The concentration of extracellular PYO found in the media is given by the measured concentration of the diluted media multiplied by the dilution factor (2):

$$16.8\ \mu M * 2 = 33.6\ \mu M$$

If we assume the media has a density of 1g/mL, the concentration on a per gram basis is

$$33.6 \frac{\mu mol}{L} * \frac{1\, L}{1000\, mL} * \frac{1\, mL}{1\, g} = 0.0336 \frac{\mu mol}{g\ media}$$

For the membrane fraction extract, which, after pooling and evaporation, consisted of 4.7mL MeOH, the measured concentration (diluted 1:10) was 125.9μM, making the true concentration in the extract

$$125.9\ \mu M * 10 = 1259.0\ \mu M$$

To find the number of moles per gram wet membrane, we can find the total moles in the 4.7mL extract and divide by the mass of the wet membrane fraction (separated by ultracentrifugation as described above):

$$1259\frac{\mu mol}{L} * \frac{1\,L}{1000\,mL} * 4.7\,mL * \frac{1}{2.75\,g\ wet\ membrane} = 2.15\,\frac{\mu mol}{g\ wet\ membrane}$$

When calculated for the whole cell pellet (mass 7.92g), we find

$$1259\frac{\mu mol}{L} * \frac{1\,L}{1000\,mL} * 4.7\,mL * \frac{1}{27.92\,g\ wet\ pellet} = 0.75\,\frac{\mu mol}{g\ wet\ pellet}$$

    https://book.bionumbers.org/how-big-is-an-e-coli-cell-and-what-is-its-mass/
